# Fractionated brain irradiation profoundly reduces hippocampal immature neuron numbers without affecting spontaneous behavior and cognition in mice

**DOI:** 10.1101/2022.09.01.506166

**Authors:** L.E. Kuil, R. Seigers, M. Loos, M.C. de Gooijer, A. Compter, W. Boogerd, O. Van Tellingen, A.B. Smit, S.B. Schagen

## Abstract

Whole brain radiotherapy (WBRT) is used to treat patients with primary brain tumors, or brain metastasis from various primary tumors to improve tumor control. However, WBRT can lead to cognitive decline. We treated mice with fractionated WBRT (fWBRT) to establish a model system to study the mechanisms underlying cognitive decline. Besides a series of traditional cognitive tests, we also assessed the effect on spontaneous behavior as measured in automated home cages.

Male C57Bl/6j mice (n=11 per group) received bi-lateral fWBRT at a dosage of 4 Gy/day on 5 consecutive days. In line with previous reports, immunohistochemical analysis of doublecortin (DCX) positive cells in the dentate gyrus showed a profound reduction in immature neurons at 4 weeks after fWBRT. Surprisingly, spontaneous behavior as measured in automated home cages was not affected. Moreover, learning and memory measured with traditional tasks - including the novel object recognition task, novel location recognition task, Barnes maze, and fear conditioning - was also not affected at 4-6 weeks after fWBRT. At 10-11 weeks after fWBRT a difference in escape latency during the learning phase, but not in the probe phase of the Barnes maze was observed.

In conclusion, although we confirmed the effect of fWBRT on neurogenesis at 4 weeks after fWBRT, we did not find clear effects on spontaneous behavior in the automated home cage nor on learning abilities as measured by traditional cognitive tasks. The relationship between the neurobiological effects of fWBRT and cognition seems more complex than often assumed and the choice of animal model, cognitive tasks, neurobiological parameters, and experimental set-up might be important factors in these types of experiments.

## Introduction

Whole brain radiotherapy (WBRT) is often used to treat patients with primary brain tumors and patients with brain metastasis from primary tumors of various origin. Where WBRT used to be the standard treatment for brain metastases, current practice increasingly advises the use of stereotactic radiosurgery (SRS) to reduce radiation-induced cognitive impairment ^1^. In fact, from 6 months onwards following WBRT, patients can experience an irreversible and progressive cognitive decline ^2^. Long-term sequelae of WBRT include a slower speed of information processing, memory retrieval deficits, and decreased executive functioning and attention ^3,4^. Many risk factors are associated with these complications following radiotherapy, including age, dose per fraction, cumulative dose, volume of the irradiated brain region and overall treatment time ^2,5^. However, it remains difficult to predict which patients will experience cognitive decline.

Pre-clinical studies have been performed in order to understand the effects of radiotherapy on cognition using various cognitive tasks, with varying results ^6^. Analysis of the brain of irradiated animals revealed alterations including reduced neurogenesis, loss of endothelial cells, loss of oligodendrocytes and microglial activation ^2,7–9^. Overall, the relation between radiation-associated cognitive impairment and radiation-induced neurobiological changes remains poorly understood, partly due to the varying outcomes of cognitive tests using rodents. Automated home cages serve as a sensitive measurement of subtle behavioral changes, independent of any confounding effects such as animal handling ^10^. Previous studies with automated home cages showed that hippocampal lesions resulted in changes in spontaneous behavior, without effects on learning simple spatial memory tasks ^11^. Nonetheless, memory was impaired if an aversive stimulus was used in a reversal learning paradigm ^11^. In addition, we previously showed that treatment with several chemotherapeutic agents affect spontaneous behavior in automated home cages, sometimes in the absence of deviant performance on traditional cognitive tasks ^12^. Preclinical studies examining both biological effects, spontaneous behavior in automated home cages and performance on traditional cognitive tests after irradiation may therefore increase our understanding of the relation between irradiation and cognitive impairment.

To establish a model system to study the mechanism of cognitive decline after fractionated WBRT (fWBRT), male C57Bl/6j mice received 4 Gray (Gy) per day (2×2 Gy bilaterally) on 5 consecutive days. To confirm loss of neurogenesis upon fWBRT, doublecortin (DCX) positive immature neuron numbers were assessed. To explore early and later effects of fWBRT on behavior and cognition, animals were tested starting at 3 and 10 weeks after fWBRT using automated home cages and a set of traditional cognitive tasks.

## Material and methods

### Animal maintenance and irradiation

Adult (11 weeks of age) male C57BL/6j mice (Charles River, France) were housed in groups of 4-5 animals in clear Plexiglas cages on a layer of wood shavings with a fixed 12:12 h light:dark cycle (lights on at 07.00 a.m.) and food and water *ad libitum*. Experiments started 2 weeks after arrival of the animals according to the protocol described below. All experiments were approved by the Animal Experimentation Committee of the Netherlands Cancer Institute (NKI hereafter) and the Vrije Universiteit (VU).

### Irradiation

Irradiation was performed using image-guided radiotherapy on the X-rad 225 μ-IGRT system (Precision X-Ray, North Branford, CT, USA) with a square beam of 7×7 mm. Animals were anesthetized using a mixture of 2.5% isoflurane and air. A sham-irradiated animal and an irradiated animal were paired, so that both animals received equivalent anesthesia. The sham animals (later referred to as sham-control) were left in the induction chamber, whereas the irradiation animals received treatment. First, a CT scan was made to correctly adjust the position of the irradiation beam for each animal. The upper part of the head and the skull was irradiated sparing the skull base, the pituitary gland, the parotid and mandibular salivary glands, the jaw muscles, and cerebellum (see figure 1A for the setup of the cone beam). The animal received 4 Gy, of which 2 Gy per hemisphere, once a day for 5 consecutive days (total dose of 20 Gy) (figure 1A). Subsequently, mice were relocated to the animal facility of the Vrije Universiteit Amsterdam (VU) according to the timelines described below and in tables 1 to 3.

**Figure 1.**
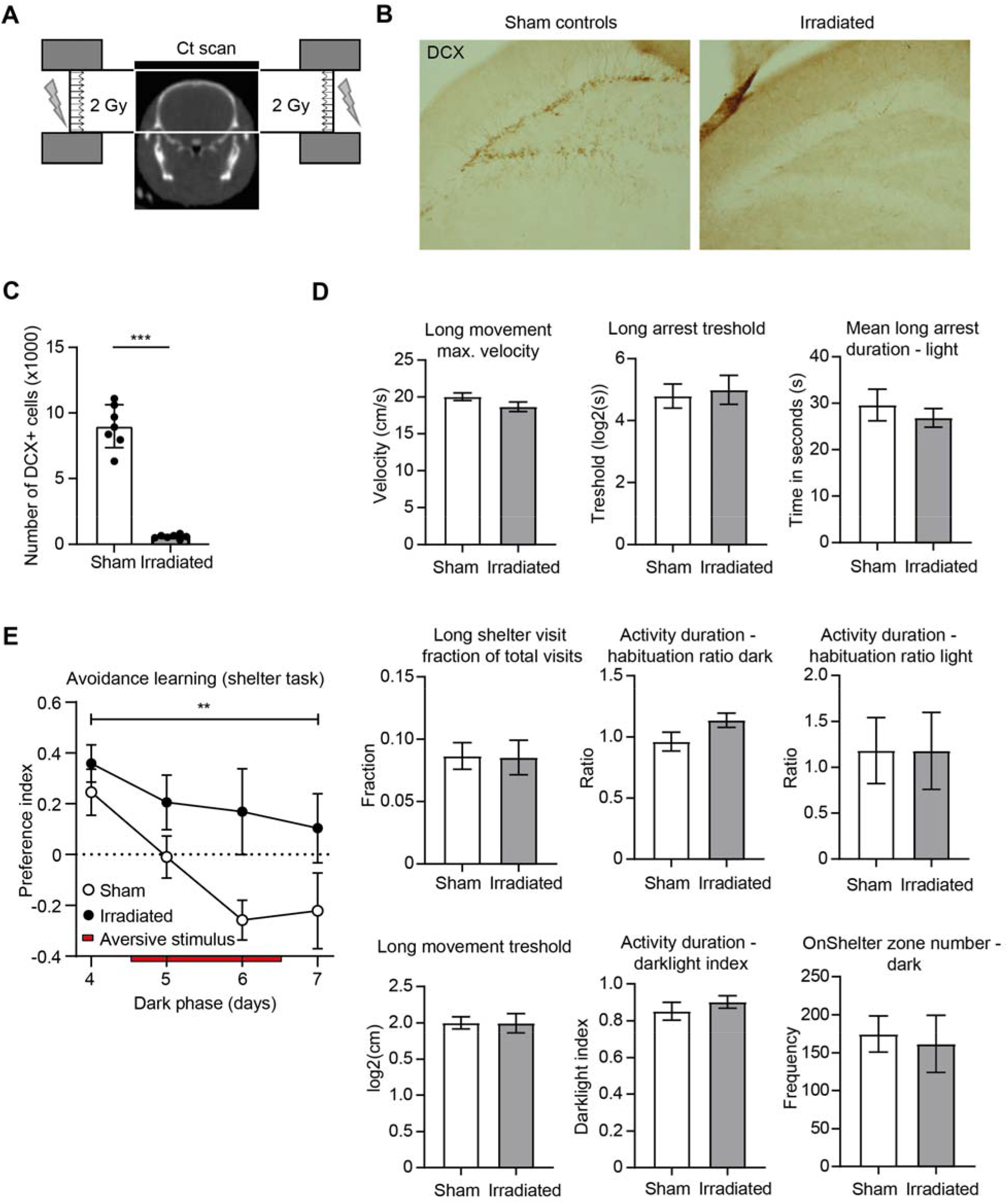
Fractionated brain irradiation reduces the number of immature neurons, but does not change spontaneous behavior in the automated home cage. Scheme for setup of the image-guided radiation therapy: Radiation beams are delivered daily for 5 days in 2 fractions of 2 Gy from both contralateral sides. B. Representative images of DCX staining in the hippocampus. C. Total number of DCX (left graph) positive cells in the dentate gyrus of the hippocampus from sham-control animals (white bar) and irradiated animals (grey bar). Irradiated animals had significantly less DCX (p < 0.001) positive cells than sham-control animals. Each dot represents the results from one animal, error bars represent standard deviation, unpaired students *t*-test. D. Graphs showing a selection of behavioral readouts from the automated home cage, showing no differences between sham-control animals (white bars) and irradiated animals (grey bars) on different parameters. CD. Bars represent the mean with error bars representing the standard error of the mean, unpaired students *t*-test were used for statistical analysis. *** p < 0.001 E. Preference index during the dark phase of day 4 to 7 of the avoidance learning (shelter task) in the automated home cage showing a preference for the entrance at day 4. At day 5 the aversive stimulus is introduced (red bar), showing that sham-controls do not show a preference anymore which continues to decline at day 6 and 7 whereas the irradiated animals retain a preference for the entrance with the aversive stimulus (F= 10.6794, p = 0.002). Dots represent the mean, error bars represent the standard error of the mean, 2-way ANOVA ** p < 0.01.

**Table 1:** Time schedule of behavioral and cognitive assays for the short-term effects of irradiation. Day 0 is the first day of irradiation. Abbreviations: NLR – Novel Location Recognition, NOR – Novel Object Recognition

| Day | -14 | 0-4 | 14 | 22-29 | 31 | 32 | 35-39 | 42-44 |
| --- | --- | --- | --- | --- | --- | --- | --- | --- |
| Activity | Arrival<br>NKI | Irradiation | Transport<br>VU | Automated<br>home cage | Open<br>field | NLR/<br>NOR | Barnes<br>maze | Fear<br>conditioning |

**Table 2:** Time schedule of cognitive assays for the long-term effects of irradiation. Day 0 is the first day of irradiation.

| Day | -14 | 0-4 | 9 | 69-73 | 76-78 |
| --- | --- | --- | --- | --- | --- |
| Activity | Arrival<br>NKI | Irradiation | Transport VU | Barnes maze | Fear conditioning |

**Table 3:** Time schedule of cognitive assays for the effects of repeated anesthesia on fear conditioning. Day 0 is the first day of anesthesia.

| Day | -14 | 0-4 | 14 | 32-34 |
| --- | --- | --- | --- | --- |
| Activity | Arrival<br>NKI/VU | Isoflurane or<br>undisturbed | Transport NKI to VU or<br>undisturbed | Fear conditioning |

### Immunohistochemistry

To explore the effects of irradiation on the brain, 7 control and 7 irradiated animals were relocated to the VU 2 weeks after irradiation started and were sacrificed and perfused 4 weeks after the start of irradiation. From the serial sections, every sixth section from each animal was selected and immunocytochemically stained with markers doublecortin (DCX) using a slightly adapted standard protocol ^13,14^ as described previously ^15^. Primary antibody (DCX: goat-anti-DCX, Santa Cruz, 1:2500), secondary antibodies (rabbit-anti-goat, 1:400, Jackson) and avidin biotinylated peroxidase complex was used (1:400, ABC Elite Kit, Vector Laboratories).

Counting of DCX positive cells in both hemispheres of the dentate gyrus was performed under a light microscope with a magnification of 400x. Quantification was performed in the subgranular layer of the dentate gyrus and counts in both blades were summed. The border of the area that was quantified was defined as the subgranular layer having a thickness of two cell diameters. All cells were counted in the subgranular layer of the dentate gyrus from top to bottom of the 40 μm thick section. Because every sixth section of the brain was stained, the number of positive cells was multiplied by 6 to get the estimated total number of DCX positive cells in the hippocampus. Pictures taken with Zeiss LSM Meta 510 confocal microscope were analyzed using ImageJ.

### Timing of behavioral and cognitive assessments

#### Early-stage observations & cognitive performance (3-6 weeks)

Ten days after the last irradiation, 11 control and 11 irradiated animals were relocated to the VU and behavioral and cognitive experiments started 1 week after relocation, which is 3 weeks after irradiation (table 1). At the VU, animals were housed individually to minimize the stress induced before and after the cognitive tasks (reduces handling and relocation of cages).

#### Later-stage cognitive performance (10-11 weeks)

Five days after the last irradiation, 12 control and 12 irradiated animals were relocated to the VU for cognitive testing. The animals were subjected to the Barnes maze and fear conditioning starting at 10 weeks after irradiation (table 2).

#### Effect of repeated anesthesia and transportation on freezing levels

Low percentage of freezing behavior was observed in the fear conditioning tasks. Therefore, we performed a control experiment to determine the effect of relocation and repeated anesthesia on fear conditioning. This experiment was simultaneous executed at both the NKI and the VU, 8 animals were subjected to 10 minutes isoflurane anesthesia for 5 days, and 8 animals were left undisturbed. Two weeks later, the animals at the NKI were relocated to the VU, while the animals at the VU were left undisturbed. All animals were subjected to the fear conditioning task 4 weeks after the isoflurane treatment (table 3).

### Automated home cage

Measurements in a home cage environment (PhenoTyper model 3000, Noldus Information Technology, Wageningen, the Netherlands) ^16,17^ were performed as previously described ^12^ and started 22 days after irradiation (short-term) and lasted 7 days. The first 3 days were used to analyze spontaneous home cage behavior where 28 measurements of kinematics, 14 measurements of sheltering, 28 measurements of habituation, 15 measurements of DarkLight index, 16 measurements of activity pattern and 14 measurements of activity give a good insight in changes in spontaneous movement in the home cage. Twenty cages were available for the analysis, animals were arbitrarily selected for the automated home cage analysis. One animal in the control group (sham) was excluded from analysis by the quality control after testing, resulting in 9 animals in the control group and 10 animals in the irradiated group.

### Open field

The open field task was executed 31 days after irradiation. The animal was placed in a white Polyvinylchloride box (50×50×50 cm) illuminated with white fluorescent light from above (60 Lux) and was allowed to explore the arena freely for 10 minutes. The box was cleaned with 70% ethanol in between animals. The box was divided into 2 zones, an outer zone and an inner zone (25x25 cm). Time spent in the inner zone (s) was analyzed with Biobserve (Biobserve GmbH). The time in the inner zone was taken as measure of anxiety as described previously ^12^.

### Novel location recognition (NLR) and novel object recognition (NOR)

The NLR and NOR tasks were given 32 days after irradiation and were executed in the open field box to explore recollection-like object memory as described previously with an inter-trial time of 2 hours between acquisition and NLR and an inter-trial time of 2 hours between NLR and NOR ^12^. One animal was excluded from analysis of the NLR specifically, as it interacted with neither object during the testing phase (Sham control group, animal 51). Measurements during the third minute of the testing period were used for statistics and figures, since at this time sufficient interaction with the objects was observed.

### Barnes maze

Spatial learning and memory in the Barnes maze were assessed 35 days (short-term) or 69 days (longer-term) after irradiation as described previously ^12^. Barnes maze training consisted of 2 sessions per day for 5 days. The mouse was allowed to explore the Barnes maze freely for a maximum of 5 minutes or until it found the escape hole. If the animal did not find the escape hole within these 5 minutes, it was guided by hand. The escape hole was removed for the last session on day 5 and replaced by a cylinder equal to the other cylinders, and this probe trial lasted for 5 minutes. Time spent (in seconds) in the zone where the escape hole used to be was analyzed providing information on how well the animals had learned to spatially locate the escape hole. Data from the first minute of the probe trial was used for analysis.

### Fear conditioning

The fear conditioning protocol started 42 days (short-term) or 76 days (longer-term) after irradiation and lasted for 3 days as described previously ^12^. The animals in experiment 4 (effect of repeated anesthesia and transportation on freezing levels) were tested with the same protocol 4 weeks after anesthesia exposure. Briefly, inter-trial time was 24 hours for contextual fear conditioning and another 24 hours later for generalized and cued fear. Freezing was defined as no movement (velocity < 1cm/s) for at least 3 seconds.

## Statistics

The learning phase of the Barnes maze was analyzed using repeated measures ANOVA with a Geisser-Greenhouse correction and a Dunnett’s multiple comparison test. The open field, NLR, NOR, probe trial of the Barnes maze, DCX quantification were analyzed using an unpaired *t*-test. The fear conditioning with two groups (short-term and longer-term experiments) was analyzed using multiple unpaired *t*-tests with a Bonferroni-Dunn correction for multiple comparisons. The fear conditioning task in the anesthesia control experiment was analyzed two-way ANOVA with a Tukey’s multiple comparisons test. For all statistical tests, a p<0.05 was considered to be statistically significant.

## Results

### Immunohistochemistry shows loss of neurogenesis after irradiation

To determine the presence of immature neurons in the hippocampus, DCX positive cells in the dentate gyrus were visualized and quantified 4 weeks after fWBRT. Irradiated animals showed significant fewer DCX positive cells than sham-control animals, indicative of severe radiotherapy-induced loss of neurogenesis in the dentate gyrus (p < 0.001, figure 1B-C).

### Early effects of irradiation on behavior and cognition

At 3 weeks after fWBRT the sham-control and irradiated animals were first subjected to automated home cages and subsequently to a sequential series of cognitive tasks, which started at 4 weeks after irradiation (Table 1). None of the parameters of spontaneous behavior as measured in the automated home cage (kinematics, sheltering, habituation, DarkLight index, activity pattern or activity) were altered in irradiated animals compared with sham-control animals (Figure 1D; supplemental data file 1). After the spontaneous behavior measured in the first three days, an avoidance learning paradigm was started from day 4. In this task, the preference for one of the two entrances to the shelter is measured. After establishing this, when the mouse uses the preferred entrance a bright light in the shelter will be turned on (day 5-6). If mice are able to learn, they will lose their preference for this entrance to avoid this bright light by either not using any entrances (sleeping outside of the shelter) or using the other entrance (output measurement). The behavior during this task is visualized as a multi-day preference index curve during the dark phase, where the sham-controls show a reduction in their preference index upon the presence of the aversive stimulus (red bar) and maintained this at day 7 (figure 1E). While sham-control animals reduce their entrances through the preferred entrance and develop a preference for the other entrance, the irradiated mice retain a preference for the aversive entrance (figure 1E).

Following the automated home cage experiments animals were allowed to acclimate for one day before being subjected to traditional cognitive tasks (Table 1). In the open field test no difference was seen in anxiety, measured by the time spend in the inner zone of the open field between the 2 groups (figure 2A). The novel location recognition (NLR) test, which is dependent on hippocampal learning, showed no changes in discrimination index after irradiation (figure 2B). The novel object recognition (NOR) test measures peri-postrhinal cortex-dependent memory and did not show learning in the sham-control group since the discrimination index was below 0,5 (figure 2C). In contrast, irradiated animals did show a preference for the novel object (discrimination index > 0,5), but the difference between groups was not significant (figure 2C). The Barnes maze is used to measure spatial learning, as contextual cues can be used to learn the location of the target hole. The escape latency during the learning phase and time spent in the escape zone during the probe phase did not differ between sham-controls and irradiated animals in the Barnes maze (figure 2D-E), suggesting there is no impairment in spatial memory. During the learning phase there was a significant time effect (repeated measures ANOVA F_2,42_=39.83, p < 0.0001), showing improvement of escape latencies over time. Multiple comparison analysis showed significant improvements between day 1 and all other days for both groups (p < 0.01; Table S1). The highest percentage time spent freezing was found in the context and cued fear conditioning set-ups, but fWBRT did not cause a significant drop in time spent freezing by any conditioning (figure 2F). This shows that context and cued fear was not affected. In the novel-fear set-up the time spent freezing was very low already in the sham-control group, which complicates the detection of differences in this set-up (figure 2F).

**Figure 2.**
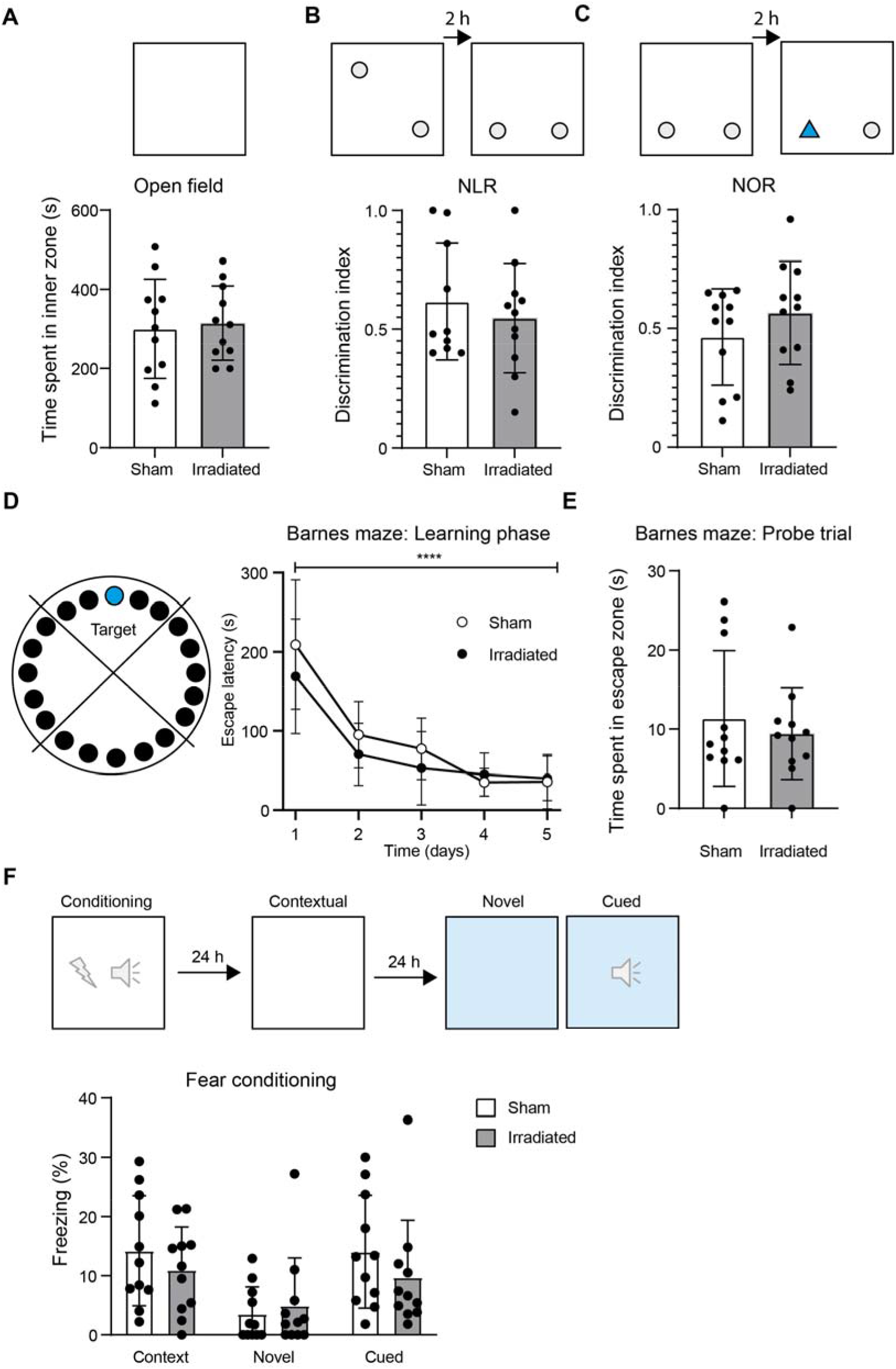
Fractionated whole brain irradiation did not result in cognitive deficits from 4 to 6 weeks after irradiation. A. Time spent in the inner zone of the open field for sham-control animals (white bar) and irradiated animals (grey bar) 31 days after irradiation did not significantly differ between groups. B. Discrimination index in the novel location recognition 26 days after irradiation did not significantly differ between groups. C. Discrimination index in the novel object recognition 26 days after irradiation did not significantly differ between groups. Note that the sham-control animals, on average, did not differentiate between the objects (discrimination index < 0,5). D. Escape latencies during the learning phase in the Barnes maze 35-39 days after irradiation. The escape latencies of the 2 trials per animal per day are set as average. No differences were observed between sham-control animals (open circles) and irradiated animals (black circles), but a significant time effect was found (repeated measures ANOVA F_2,42_=39.83, p < 0.0001). E. Time spent in the zone were the escape hole used to be during the first minute of the probe trial for shamcontrol animals (white bar) and irradiated animals (grey bar) at 39 days after irradiation. F. Percentage freezing in the fear conditioning during contextual, novel, and cued fear for sham-control animals (white bars) and irradiated animals (grey bars) 42-44 days after irradiation. A-C, E-F Each dot represents the results from one animal, error bars represent standard deviation. D. Dots represent the mean, error bars represent the standard deviation. Statistics were performed using unpaired t-tests (A-C, E), repeated measures ANOVA (D) Unpaired t-test with Bonferroni-Dunn correction for multiple comparisons (E).

### Longer-term effects of irradiation on cognition

Since cognitive decline after irradiation appears to be a delayed effect in patients, often presenting from 6 months after fWBRT onwards ^2^, we performed spatial learning and fear conditioning tests starting at 10 weeks after irradiation (Table 2). There was an overall significant group effect (repeated measures ANOVA: F_1,22_=7.406, p = 0.012, figure 3A) and time effect (repeated measures ANOVA: F_2,54_=17,96, p < 0.0001, figure 3A) on escape latency during the learning phase of the Barnes maze. This shows that irradiated animals have a significantly longer escape latency, but over time show significant improvement suggesting learning (Day 1 versus 4 p = 0.01; day 1 versus 5 p = 0.007). The sham-control group also showed significant improvement between day 1 and day 4 (p = 0.005) and 5 (p = 0.002). During the probe trial no difference was observed in the time spent in the correct quadrant (figure 3B). In both the novel and cued fear conditioning set-ups the sham-control mice already exhibited low percentages of time spent freezing ^18–21^, which complicates interpretation (figure 3C). In contrast, sham-control mice spent more time freezing in the contextual fear condition, allowing the potential observation of a negative impact of fWBRT on fear conditioning (figure 3C). However, also in this conditioning set-up no differences were found in the time spent freezing between sham-controls and irradiated animals (figure 3C).

**Figure 3.**
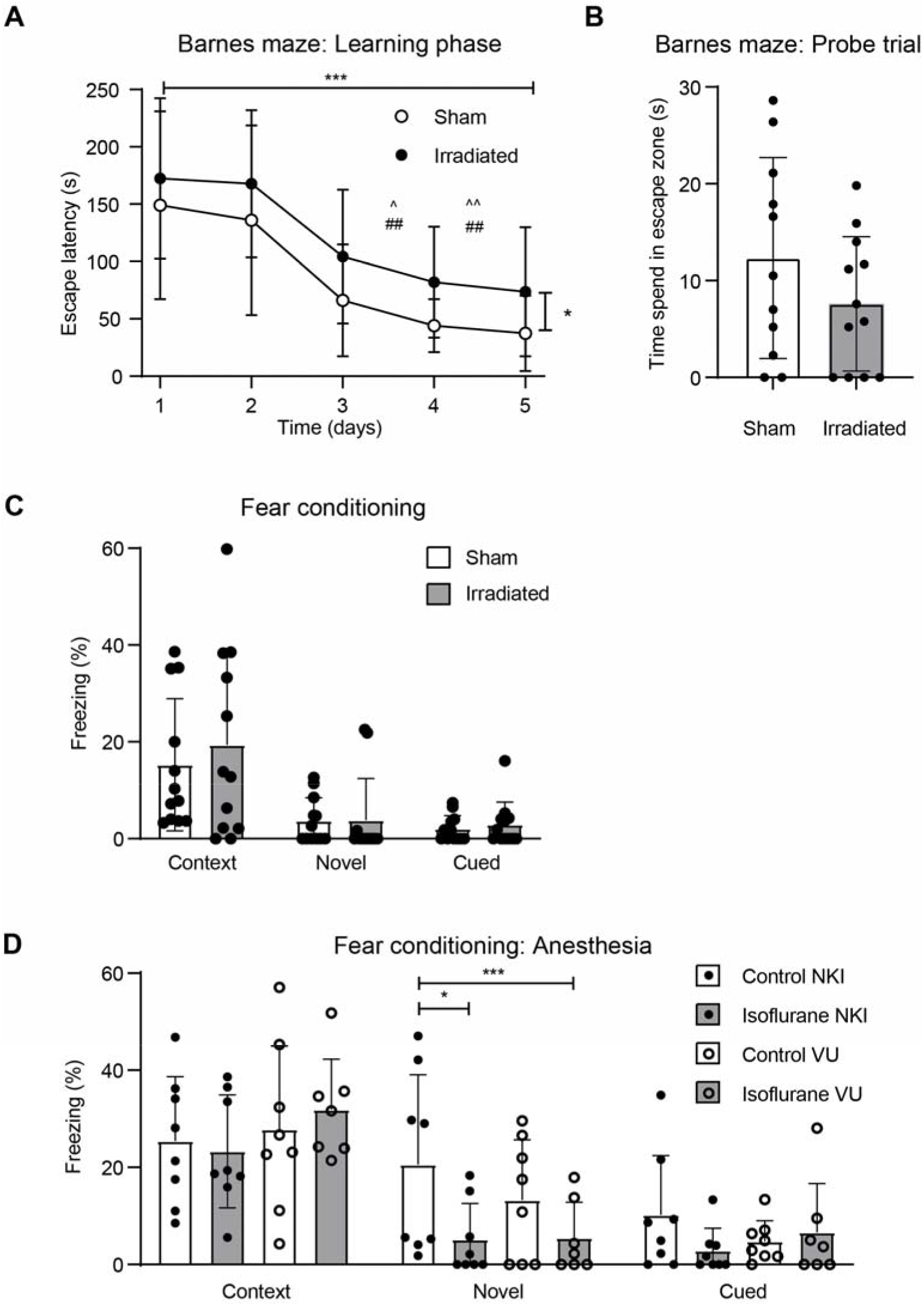
Cognitive assays on animals that underwent repeated anesthesia and animals 10 weeks after fractionated irradiation. A. Escape latency in the Barnes maze for sham-control animals (open circles) and irradiated animals (black circles) 10 weeks after irradiation. The escape latencies of the 2 trials per animal per day are set as average. There was an overall significant group effect (repeated measures ANOVA: F_1,22_=7.406, p = 0.01) and time effect (repeated measures ANOVA: F_2,54_=17,96, p < 0.0001) in learning phase. ^ = Irradiated group p < 0.05 compared to day 1, ^^ = Irradiated group p <0.01 compared to irradiated day 1, ## = Sham-control group p <0.01 compared to sham-control day 1. Each dot represents the mean and error bars represent the standard deviation. B. Time spent in the zone were the escape hole used to be during the first minute of the probe trial for sham-control animals (white bar) and irradiated animals (grey bar) 10 weeks after irradiation. No statistical differences were found, students *t*-test. C. Percentage freezing in the fear conditioning during contextual, novel, and cued fear for control animals (white bar) and irradiated animals (grey bar) 10 weeks after irradiation showed no differences between groups. Unpaired *t*-test with Bonferroni-Dunn correction for multiple comparisons D. Percentage freezing in the fear conditioning during contextual, novel, and cued fear for control animals at the NKI (white bar with black dot), control animals at the VU (white bars with open circle), isoflurane anesthesia animals at the NKI (grey bar with black dot), and isoflurane anesthesia animals at the VU (grey bar with open circle). A main effect was found on novel fear (F (2, 81) = 27,38; p < 0,0001), which was explained by the effect of Isoflurane on novel fear conditioning in the NKI population specifically (p = 0.047). Each dot represents one animal, error bars represent standard deviation, two-way ANOVA with a Tukey’s multiple comparisons test was used for statistical analysis. B-D. All dots represent one animal, error bars represent the standard deviation.

### Repeated anesthesia reduces freezing levels in the novel fear conditioning specifically

Since low percentages of freezing behavior were observed in our fear conditioning experiments, we set up an experiment to examine whether the stressors in this experiment (repeated isoflurane anesthesia and transportation) may have affected our experimental results in the fear conditioning task. Stress is known to have a negative effect on cognition ^22^. Control animals were left undisturbed (VU control), only underwent transportation (NKI control) and experimental animals were anesthetized repeatedly with isoflurane with transportation (NKI isoflurane) or without transportation (VU isoflurane). The animals were subjected to contextual, novel and cued fear conditioning 4 weeks after repeated exposure to isoflurane anesthesia (Table 3). No difference was seen in contextual or cued fear between animals that were subjected to anesthesia and those that were not, nor between animals from the NKI and the VU. However, a main effect was found in the novel fear conditioning (F_2,81_ = 27,38; p < 0,0001), which was explained by the effect of Isoflurane on novel fear conditioning in the NKI population (p = 0.047) (figure 3D). The anesthetized animals at the VU also showed a reduced percentage of freezing behavior, although not statistically significant (Figure 3D). Overall, repeated anesthesia reduces freezing behavior in novel fear conditioning, but not contextual fear conditioning.

## Discussion

Neurogenesis has repeatedly shown to be decreased after brain irradiation independent of age, sex or species used ^23–33^, which is in line with our findings showing a reduction of immature neurons in the dentate gyrus. However, we did not detect changes in spontaneous behavior, suggesting that the irradiated animals do not show abnormalities in activity, mobility and behavioral patterns. This is in contrast to our previous findings using chemotherapeutics where we did detect changes in spontaneous behavior, as well as cognition ^12^ and that of others that reported changes in spontaneous behavior during natural aging ^34^. The effect of fWBRT on cognition is also limited in our experiments, since only defects in avoidance learning in the automated home cage 3 weeks after fWBRT and subtle differences in the learning phase of the Barnes maze at 10 weeks after fWBRT were detected. A recent study corroborates this pattern of findings as they reported significant loss of neurogenesis but no changes in contextual fear, spatial memory (Morris water maze), NOR and open field task ^35^.

Overall, whereas effects of WBRT on neurobiology seem clear and robust, the results from cognitive studies using WBRT in rodents are inconclusive. Some studies failed to detect changes in spatial memory (mainly Morris water maze) upon irradiation ^23,24,36,37^, however, others could find changes ^25,38-44^. Some interesting observations were made with respect to (1) the use of deviating spatial strategies in spatial memory tasks by irradiated animals ^45^, (2) enhanced extinction of memory upon irradiation and (3) deficits in cognitive flexibility in irradiated animals ^46^. However, these types of changes might not be measured by the traditional cognitive tasks and therefore the effects of irradiation on cognition might not always be detected. Therefore, more complex learning tasks including for example reversal learning might give better insight in impairments after WBRT. One can choose to perform the novel object recognition (NOR) task and the novel location recognition (NLR) tasks using a short retention interval (minutes) as a measurement of hippocampus-independent, prefrontal cortex-dependent memory, or long retention interval (hours) for hippocampal-dependent learning. While the short-term interval version resulted in defects after irradiation ^23,24,41,47–52^, the hippocampal-dependent version of the NOR was not always affected after irradiation ^53–55^. Interestingly, performance on the NOR was found to be dependent on dose, dosing-regimen and timing ^53,54,56^. In addition, male C57BL/6 mice only perform well at the NOR at 3 months, but not at older age ^57^ and short timeframes are required to capture NOR ^57^. Therefore, we only analyzed one minute of the five-minute test phase for analysis, but still the sham-controls did not show preference. Moreover, various mouse strains perform well on the 1 hour retention version of the NOR but failed to recognize or remember the familiar object in the 24 hour retention version ^58^. Regardless of differences in dosing regimen, the effects of irradiation and/or loss of neurogenesis on fear conditioning are unclear; reporting defects ^59–61^, or no changes ^35,53,62^. Drew *et al*. found that defects were only observed in single-trial contextual fear conditioning (delayed shock) after low-dose irradiation, but not in the repeated-trial contextual fear conditioning ^63^. This once again shows that irradiation seems to induce specific behavioral abnormalities that regular cognitive tasks might not always measure.

## Limitations

First of all, our behavioral and cognitive assessments were performed between 3 and 11 weeks after irradiation, which might still be too short after irradiation to develop behavioral and cognitive abnormalities, since effects are usually seen only after 6 months in patients ^2^. Second, in the avoidance learning task in the automated home cage there are some factors that influence performance in this task. For example, the extremely bright light that shines upon entry through the preferred entrance might interfere with the circadian rhythm of the mice. Also, the mice can choose to sleep outside of the shelter, thus stopping using the entrances altogether, which is not accounted for in the readout. Third, in the current study the sham-control animals did not learn the NOR task, since they showed a preference index below 0,5 on average, meaning that they did not show a preference for one of the objects. The change in shape of an object might not be as meaningful to the mice, since rodents mostly rely on tactile stimuli. Last, in the fear conditioning it appears that the animals in our cognitive studies showed a relatively low percentage of time spent freezing, which hampers the detection of differences between groups. Our control experiment showed reduced freezing behavior specifically in the novel fear task after repeated anesthesia. Therefore, it is particularly hard to detect reduced freezing in this specific part of the paradigm, which likely hampered our assessments.

## Conclusion

In our animal model the number of immature neurons in the dentate gyrus was strongly reduced 4 weeks after fWBRT. Spontaneous behavior and cognition were not affected at any time point, except for the learning phase of the Barnes maze at 10 weeks after fWBRT. No deficits in the open field, novel object recognition, novel location recognition, the probe trial of the Barnes maze, and fear conditioning were observed. The data in this paper underscores the complexity of the relation between irradiation, neurogenesis and cognition.

## Supplemental data file 1.

Report of the full outcome list from the automated home cage.

