## Supplemental data file 1 for "Fractionated brain irradiation profoundly reduces hippocampal immature neuron numbers without affecting spontaneous behavior and cognition in mice"

Sylics

**Strain: C57BL/6J**

| Gene | Locus | Official name | Aliases | Ensembl ID | MGI ID | JAX ID |
|---|---|---|---|---|---|---|
| C57BL/6J | | | | | | |

**General MGI ID:**
**General JAX ID:**
**Comments:**

 *© Sylics BV*

### HTP Data

 *© Sylics BV*

### Avoidance learning (shelter task)

The PhenoTyper is equipped with a shelter compartment, with two entrances, in which mice typically spend 80% of their time (resting/sleeping). During the first 4 days, mice develop a preference to enter the shelter through one of the two entrances. The preference index is calculated by : [(number of entries through the preferred entrance) - (number of entries through
non-preferred entrance)]/(total number of entries).
Avoidance learning is studied by automatically applying a mild aversive stimulus (shelter illumination with bright light) during days 5 and 6 each time mice entered the shelter using their preferred entrance, but not when using the other entrance. A reduction in the preference index indicates that a mouse is establishing a specific association between its preferred entrance and the aversive stimulus. During day 7 sanctioning (i.e. shelter illumination) is discontinued, and the stability of the change in preference can be determined.
Avoidance learning is best studied during the Dark phase when shelter illumination is a stronger stimulus than during the light phase. For a detailed explanation see *Maroteaux et al.* (http://www.ncbi.nlm.nih.gov/pubmed/22846151)
The behavior during this task can be visualized in the multi-day preference index curve (plotting Dark phase 4 -7), see C57BL/6J mice as an example:
mousedata.sylics.com/?page=strainhtp&loaded=true&s=11&htpid=243&e=2&batch=1&t=&texp=&list=batches&tdose=

#### Preference index

*Preference index*: The preference index was determined as follows: [(number of entries through the preferred entrance) - (number of entries through non-preferred entrance)]/(total number of entries).

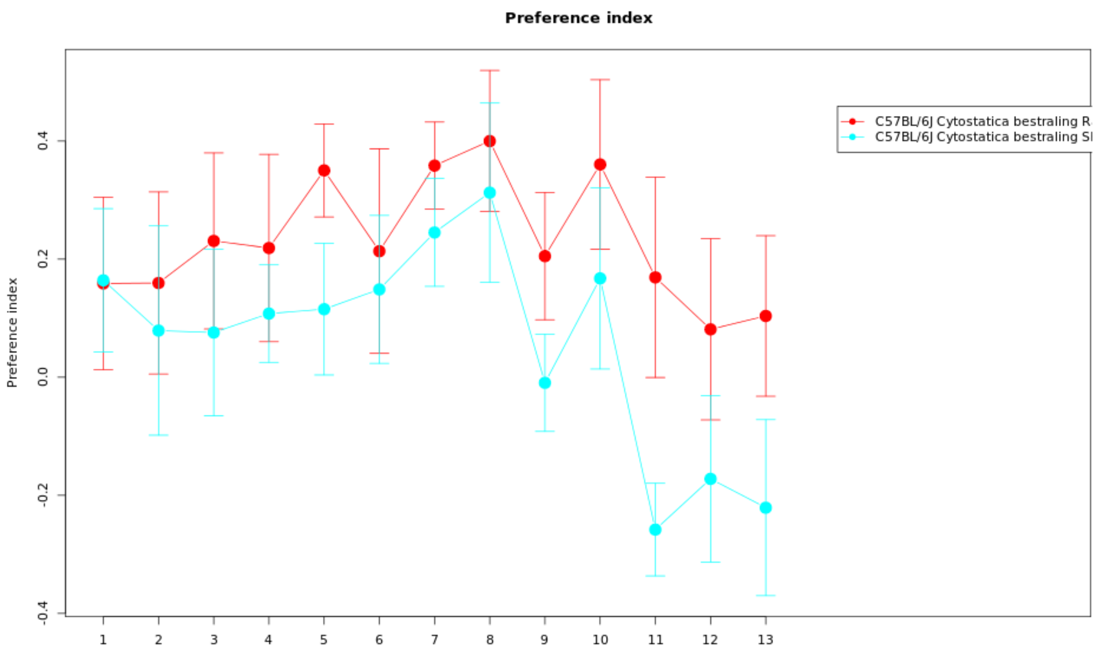

Per data point info

| Genotype | Mean | Median | SEM | N |
|---|---|---|---|---|
| (Gr 1): C57BL/6J Cytostatica bestraling Radiation (batch 22) (1) | 0.1589 | 0.252 | 0.1458 | 10 |
| (Gr 1): C57BL/6J Cytostatica bestraling Radiation (batch 22) (2) | 0.1596 | 0.0732 | 0.1542 | 10 |
| (Gr 1): C57BL/6J Cytostatica bestraling Radiation (batch 22) (3) | 0.2308 | 0.2989 | 0.1491 | 10 |
| (Gr 1): C57BL/6J Cytostatica bestraling Radiation (batch 22) (4) | 0.2188 | 0.1774 | 0.1587 | 10 |
| (Gr 1): C57BL/6J Cytostatica bestraling Radiation (batch 22) (5) | 0.3503 | 0.333 | 0.0788 | 10 |
| (Gr 1): C57BL/6J Cytostatica bestraling Radiation (batch 22) (6) | 0.2135 | 0.3068 | 0.1732 | 10 |
| (Gr 1): C57BL/6J Cytostatica bestraling Radiation (batch 22) (7) | 0.3584 | 0.3263 | 0.0738 | 10 |
| (Gr 1): C57BL/6J Cytostatica bestraling Radiation (batch 22) (8) | 0.4 | 0.3297 | 0.1196 | 10 |
| (Gr 1): C57BL/6J Cytostatica bestraling Radiation (batch 22) (9) | 0.2052 | 0.2711 | 0.1079 | 10 |
| (Gr 1): C57BL/6J Cytostatica bestraling Radiation (batch 22) (10) | 0.3604 | 0.4248 | 0.1435 | 10 |
| (Gr 1): C57BL/6J Cytostatica bestraling Radiation (batch 22) (11) | 0.1691 | 0.197 | 0.1695 | 10 |
| (Gr 1): C57BL/6J Cytostatica bestraling Radiation (batch 22) (12) | 0.0811 | -0.1404 | 0.1534 | 10 |
| (Gr 1): C57BL/6J Cytostatica bestraling Radiation (batch 22) (13) | 0.1037 | 0.2418 | 0.136 | 10 |
| (Gr 2): C57BL/6J Cytostatica bestraling Sham (batch 22) (1) | 0.1641 | 0.3605 | 0.1214 | 9 |
| (Gr 2): C57BL/6J Cytostatica bestraling Sham (batch 22) (2) | 0.079 | 0.1538 | 0.1772 | 9 |
| (Gr 2): C57BL/6J Cytostatica bestraling Sham (batch 22) (3) | 0.0757 | 0.1346 | 0.1412 | 9 |
| (Gr 2): C57BL/6J Cytostatica bestraling Sham (batch 22) (4) | 0.1077 | 0.1667 | 0.0825 | 9 |
| (Gr 2): C57BL/6J Cytostatica bestraling Sham (batch 22) (5) | 0.1152 | 0.25 | 0.1114 | 9 |
| (Gr 2): C57BL/6J Cytostatica bestraling Sham (batch 22) (6) | 0.1487 | 0.2727 | 0.1253 | 9 |
| (Gr 2): C57BL/6J Cytostatica bestraling Sham (batch 22) (7) | 0.2451 | 0.1368 | 0.0913 | 9 |
| (Gr 2): C57BL/6J Cytostatica bestraling Sham (batch 22) (8) | 0.3126 | 0.3333 | 0.1516 | 9 |
| (Gr 2): C57BL/6J Cytostatica bestraling Sham (batch 22) (9) | -0.0095 | -0.0517 | 0.0824 | 9 |
| (Gr 2): C57BL/6J Cytostatica bestraling Sham (batch 22) (10) | 0.1674 | 0 | 0.1536 | 9 |

| | | | | |
|---|---|---|---|---|
| (Gr 2): C57BL/6J Cytostatica bestraling Sham (batch 22) (11) | -0.2581 | -0.25 | 0.0783 | 9 |
| (Gr 2): C57BL/6J Cytostatica bestraling Sham (batch 22) (12) | -0.1722 | -0.2364 | 0.1407 | 9 |
| (Gr 2): C57BL/6J Cytostatica bestraling Sham (batch 22) (13) | -0.221 | -0.4351 | 0.1491 | 9 |

2-Way ANOVA statistics

| 2-way ANOVA | Df | Sum Sq | Mean Sq | F value | Pr(>F) |
|---|---|---|---|---|---|
| genotype | 1 | 1.7767 | 1.7767 | 10.4625 | 0.0014 |
| parametername | 1 | 0.7163 | 0.7163 | 4.218 | 0.0411 |
| genotype:parametername | 1 | 0.4501 | 0.4501 | 2.6508 | 0.1048 |
| Residuals | 242 | 41.0945 | 0.1698 | NA | NA |

| Mice excluded by Quality Control |
|---|
| PH06469 (C57BL/6J - ) |

*© Sylics BV*

Sylics

Aversion index

*Aversion index*: The aversion index was defined as follows: [(time spent in the illuminated shelter after entering through the sanctioned entrance) - (time spent in the dark shelter after entering through the non-sanctioned entrance)]/(total time spent in shelter)

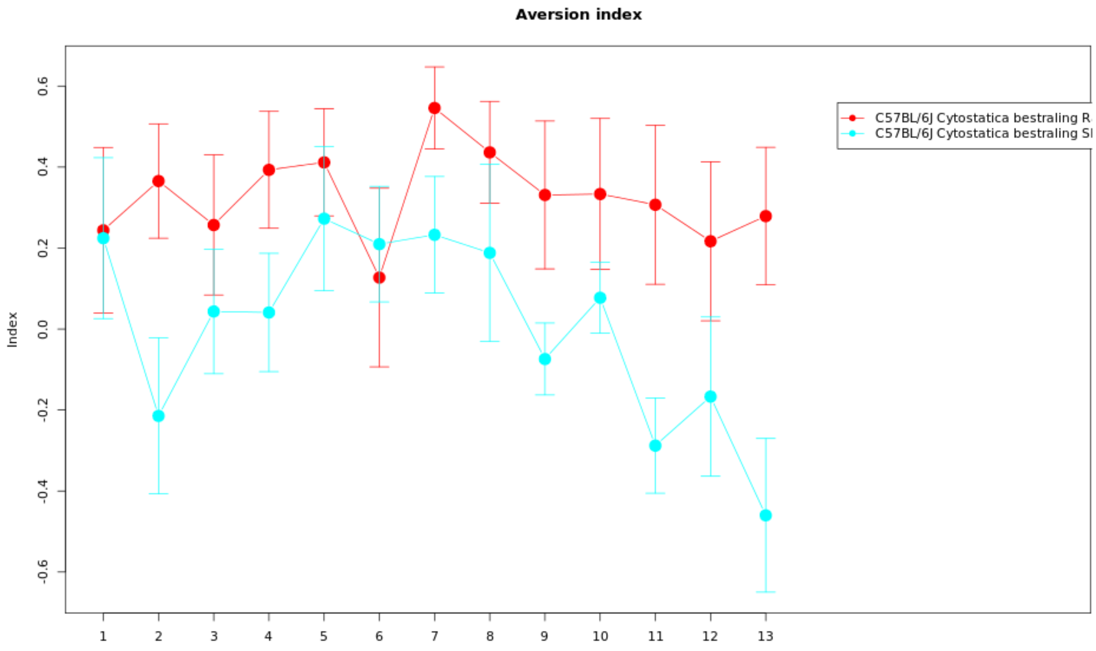

Per data point info

| Genotype | Mean | Median | SEM | N |
|---|---|---|---|---|
| (Gr 1): C57BL/6J Cytostatica bestraling Radiation (batch 22) (1) | 0.2439 | 0.3737 | 0.2046 | 10 |
| (Gr 1): C57BL/6J Cytostatica bestraling Radiation (batch 22) (2) | 0.3655 | 0.391 | 0.1413 | 10 |
| (Gr 1): C57BL/6J Cytostatica bestraling Radiation (batch 22) (3) | 0.257 | 0.2916 | 0.1729 | 10 |
| (Gr 1): C57BL/6J Cytostatica bestraling Radiation (batch 22) (4) | 0.3931 | 0.5075 | 0.1443 | 10 |
| (Gr 1): C57BL/6J Cytostatica bestraling Radiation (batch 22) (5) | 0.4116 | 0.5384 | 0.1322 | 10 |
| (Gr 1): C57BL/6J Cytostatica bestraling Radiation (batch 22) (6) | 0.1272 | 0.2206 | 0.2205 | 10 |
| (Gr 1): C57BL/6J Cytostatica bestraling Radiation (batch 22) (7) | 0.5458 | 0.6578 | 0.1015 | 10 |
| (Gr 1): C57BL/6J Cytostatica bestraling Radiation (batch 22) (8) | 0.4363 | 0.4764 | 0.1258 | 10 |
| (Gr 1): C57BL/6J Cytostatica bestraling Radiation (batch 22) (9) | 0.3311 | 0.5672 | 0.1825 | 10 |
| (Gr 1): C57BL/6J Cytostatica bestraling Radiation (batch 22) (10) | 0.3335 | 0.5955 | 0.1865 | 10 |
| (Gr 1): C57BL/6J Cytostatica bestraling Radiation (batch 22) (11) | 0.3069 | 0.4847 | 0.1964 | 10 |
| (Gr 1): C57BL/6J Cytostatica bestraling Radiation (batch 22) (12) | 0.2168 | 0.2356 | 0.1964 | 10 |
| (Gr 1): C57BL/6J Cytostatica bestraling Radiation (batch 22) (13) | 0.2789 | 0.3313 | 0.1699 | 10 |
| (Gr 2): C57BL/6J Cytostatica bestraling Sham (batch 22) (1) | 0.2245 | 0.3588 | 0.1986 | 9 |
| (Gr 2): C57BL/6J Cytostatica bestraling Sham (batch 22) (2) | -0.2144 | -0.1887 | 0.1926 | 9 |
| (Gr 2): C57BL/6J Cytostatica bestraling Sham (batch 22) (3) | 0.0435 | -0.0316 | 0.1538 | 9 |
| (Gr 2): C57BL/6J Cytostatica bestraling Sham (batch 22) (4) | 0.0409 | 0.1035 | 0.146 | 9 |
| (Gr 2): C57BL/6J Cytostatica bestraling Sham (batch 22) (5) | 0.2726 | 0.4735 | 0.1779 | 9 |
| (Gr 2): C57BL/6J Cytostatica bestraling Sham (batch 22) (6) | 0.2097 | 0.156 | 0.1427 | 9 |
| (Gr 2): C57BL/6J Cytostatica bestraling Sham (batch 22) (7) | 0.2327 | 0.1191 | 0.144 | 9 |
| (Gr 2): C57BL/6J Cytostatica bestraling Sham (batch 22) (8) | 0.1882 | 0.0188 | 0.2189 | 9 |
| (Gr 2): C57BL/6J Cytostatica bestraling Sham (batch 22) (9) | -0.074 | -0.1404 | 0.0888 | 9 |

 © Sylics BV

| | | | | |
|---|---|---|---|---|
| (Gr 2): C57BL/6J Cytostatica bestraling Sham (batch 22) (10) | 0.0773 | 0.0787 | 0.0873 | 9 |
| (Gr 2): C57BL/6J Cytostatica bestraling Sham (batch 22) (11) | -0.2877 | -0.332 | 0.1176 | 9 |
| (Gr 2): C57BL/6J Cytostatica bestraling Sham (batch 22) (12) | -0.1667 | -0.2024 | 0.1967 | 9 |
| (Gr 2): C57BL/6J Cytostatica bestraling Sham (batch 22) (13) | -0.4599 | -0.6908 | 0.1898 | 9 |

2-Way ANOVA statistics

| 2-way ANOVA | Df | Sum Sq | Mean Sq | F value | Pr(>F) |
|---|---|---|---|---|---|
| genotype | 1 | 6.153 | 6.153 | 23.1579 | 0 |
| parametername | 1 | 0.9014 | 0.9014 | 3.3926 | 0.0667 |
| genotype:parametername | 1 | 0.7686 | 0.7686 | 2.8928 | 0.0903 |
| Residuals | 242 | 64.2994 | 0.2657 | NA | NA |

| Mice excluded by Quality Control |
|---|
| PH06469 (C57BL/6J - ) |

 *© Sylics BV*

Preference index - Dark phase

*Preference index on Dark 4*:

*Preference index on Dark 5*:

*Preference index on Dark 6*:

*Preference index on Dark 7*:

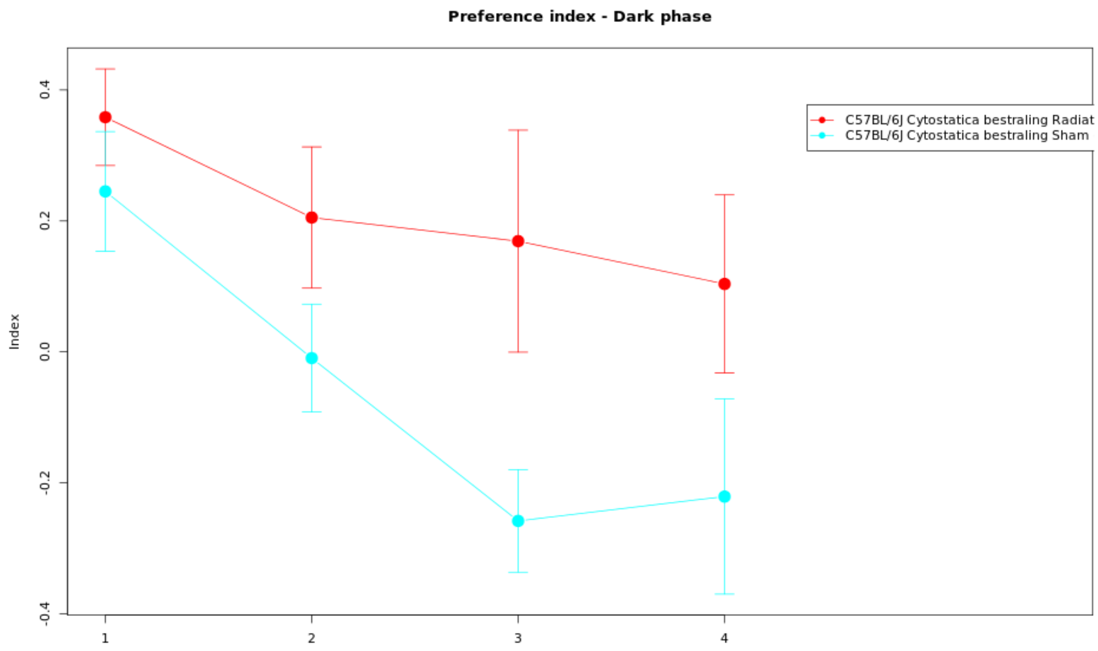

Per data point info

| Genotype | Mean | Median | SEM | N |
|---|---|---|---|---|
| (Gr 1): C57BL/6J Cytostatica bestraling Radiation (batch 22) (1) | 0.3584 | 0.3263 | 0.0738 | 10 |
| (Gr 1): C57BL/6J Cytostatica bestraling Radiation (batch 22) (2) | 0.2052 | 0.2711 | 0.1079 | 10 |
| (Gr 1): C57BL/6J Cytostatica bestraling Radiation (batch 22) (3) | 0.1691 | 0.197 | 0.1695 | 10 |
| (Gr 1): C57BL/6J Cytostatica bestraling Radiation (batch 22) (4) | 0.1037 | 0.2418 | 0.136 | 10 |
| (Gr 2): C57BL/6J Cytostatica bestraling Sham (batch 22) (1) | 0.2451 | 0.1368 | 0.0913 | 9 |
| (Gr 2): C57BL/6J Cytostatica bestraling Sham (batch 22) (2) | -0.0095 | -0.0517 | 0.0824 | 9 |
| (Gr 2): C57BL/6J Cytostatica bestraling Sham (batch 22) (3) | -0.2581 | -0.25 | 0.0783 | 9 |
| (Gr 2): C57BL/6J Cytostatica bestraling Sham (batch 22) (4) | -0.221 | -0.4351 | 0.1491 | 9 |

2-Way ANOVA statistics

| 2-way ANOVA | Df | Sum Sq | Mean Sq | F value | Pr(>F) |
|---|---|---|---|---|---|
| genotype | 1 | 1.3801 | 1.3801 | 10.6794 | 0.0017 |
| parametername | 1 | 1.3705 | 1.3705 | 10.605 | 0.0017 |
| genotype:parametername | 1 | 0.1699 | 0.1699 | 1.3145 | 0.2554 |
| Residuals | 71 | 9.1752 | 0.1292 | NA | NA |

| Mice excluded by Quality Control |
|---|
| PH06469 (C57BL/6J - ) |

© Sylics BV

### Spontaneous behavior (Group 1: kinematics)

With respect to spontaneous behaviors in the first three days in the PhenoTyper, 6 groups of behavioral parameters are defined as described below. The first two groups describe specific behavioral elements related to kinematics of mice (description of movement characteristics, group1) and sheltering behavior (group 2). These behavioral parameters were analyzed with respect to temporal aspects, in particular over 4 different time scales, i.e., habituation effects across multiple days (group 3), effects of DarkLight phase across 24h (group 4), differences in the pattern of behavior in the few hours before and after phase shifts (group5), and differences in activity bout properties on the sub-minute time scale (group 6).

**Group 1: kinematics**

Movement segments:

Mice make short movements, such as turning or rearing against the wall, as well as long movements when mice travel from one location in the cage to the next. These two types of movement, i.e. short and long, can be distinguished by plotting the frequency distribution of all move segment distances over three days in the cage, yielded a bimodal distribution. For instance, see the "histogram of movement distance" at the bottom of this page for C57BL/6J mice:
http://mousedata.sylics.com/?page=strainhtp&loaded=true&s=11&htpid=53&e=9&batch=1&t=&texp=&list=batches&tdose=

An animal-centered threshold that was defined by the intersection of the two Gaussian distance curves obtained after Gaussian mixture model fitting of data of each individual mouse.

Arrest segments:

Most arrest durations are relatively short, and occur in between move segments when mice reorient themselves. In addition, mice infrequently made long arrests when they were consuming food or water, or when they were resting outside their shelter. An animal-centered threshold was set distinguishing the 90% shortest arrests (short) from the 10% longest arrests (long), separating brief from long arrests. For instance, see the "histogram of arrest durations" at the bottom of this page for C57BL/6J mice:
http://mousedata.sylics.com/?page=strainhtp&loaded=true&s=11&htpid=55&e=9&batch=1&t=&texp=&list=batches&tdose=

 *© Sylics BV*

Long movement fraction of total movement

*Long movement fraction of total movement*: The fraction of movement segments with distance larger than long movement threshold

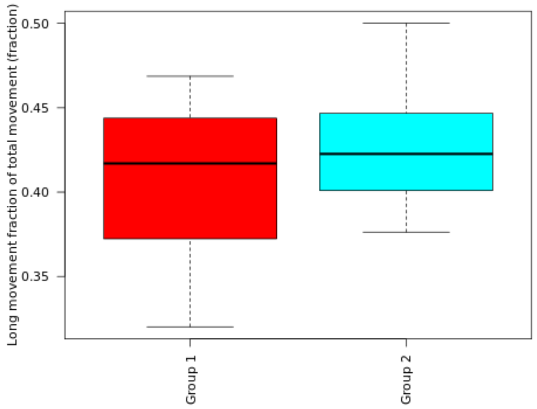

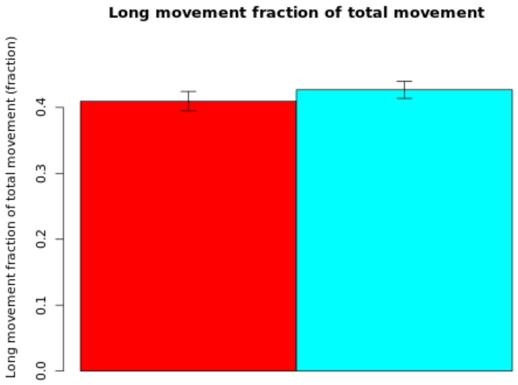

**Secondary plot:**

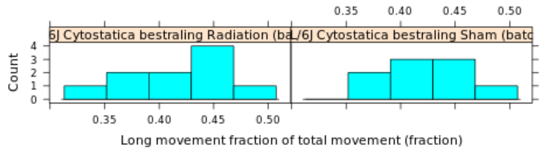

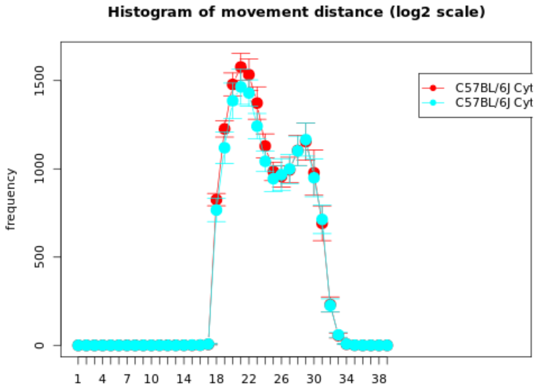

| Genotype | Mean | Median | SEM | N |
|---|---|---|---|---|
| (Gr 1): C57BL/6J Cytostatica bestraling Radiation (batch 22) | 0.4096 | 0.417 | 0.0147 | 10 |
| (Gr 2): C57BL/6J Cytostatica bestraling Sham (batch 22) | 0.4268 | 0.4226 | 0.0129 | 9 |

| | |
|---|---|
| **T-test:** | -0.8788 |
| **P-value:** | 0.3918 |
| **Df:** | 16.9173 |
| **Mann-Whitney U test:** | 0.6038 |

| Mice excluded by Quality Control |
|---|
| PH06469 (C57BL/6J - ) |

© Sylics BV

Long movement max. velocity

*Long movement max. velocity*: Average velocity of the 95th percentile fastest long movement segments

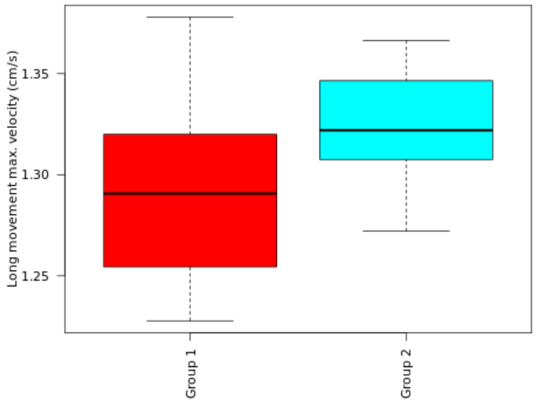

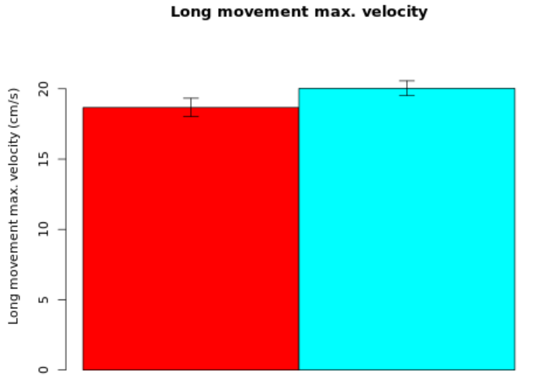

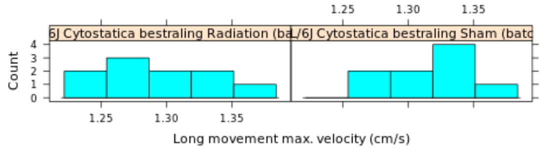

#### Secondary plot:

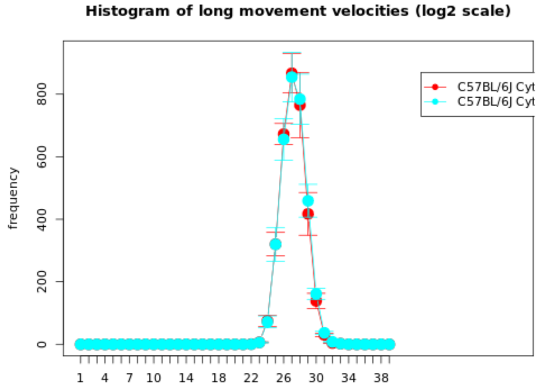

**NOTE: The statistics and boxplot are performed on log10 transformed data**

| Genotype | Mean | Median | SEM | N |
|---|---|---|---|---|
| (Gr 1): C57BL/6J Cytostatica bestraling Radiation (batch 22) | 18.6673 | 0 | Up: 0.6594<br>Down: 0.638 | 10 |
| (Gr 2): C57BL/6J Cytostatica bestraling Sham (batch 22) | 20.0328 | 0 | Up: 0.5328<br>Down: 0.5196 | 9 |

| | |
|---|---|
| **T-test:** | -1.6218 |
| **P-value:** | 0.1241 |
| **Df:** | 16.2754 |
| **Mann-Whitney U test:** | 0.1333 |

| Mice excluded by Quality Control |
|---|
| PH06469 (C57BL/6J - ) |

#### Long arrest threshold

*Long arrest threshold*: Cut-off value to separate short and long arrests

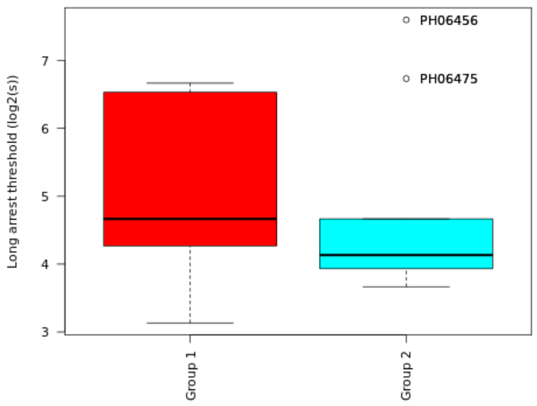

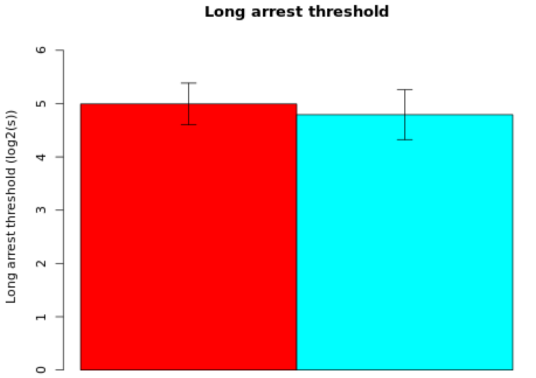

##### Secondary plot:

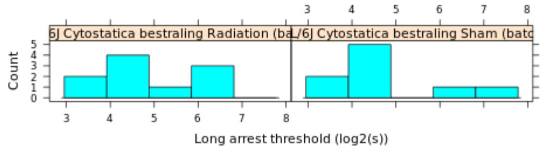

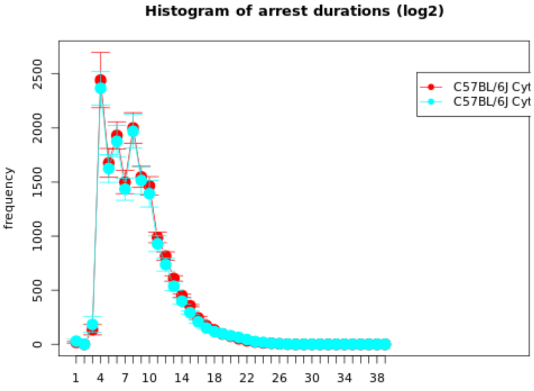

| Genotype | Mean | Median | SEM | N |
|---|---|---|---|---|
| (Gr 1): C57BL/6J Cytostatica bestraling Radiation (batch 22) | 4.9933 | 4.6667 | 0.3912 | 10 |
| (Gr 2): C57BL/6J Cytostatica bestraling Sham (batch 22) | 4.7926 | 4.1333 | 0.4675 | 9 |

| | |
|---|---|
| **T-test:** | 0.3293 |
| **P-value:** | 0.7462 |
| **Df:** | 16.106 |
| **Mann-Whitney U test:** | 0.5948 |

| Mice excluded by Quality Control |
|---|
| PH06469 (C57BL/6J - ) |

© Sylics BV

Short arrest duration - dark

*Short arrest duration - dark*: Cummulative duration of short arrests during the dark phase

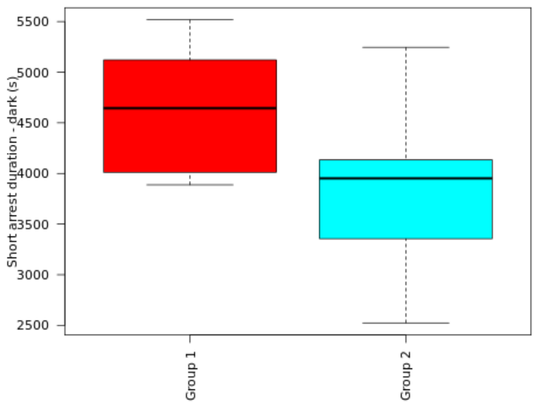

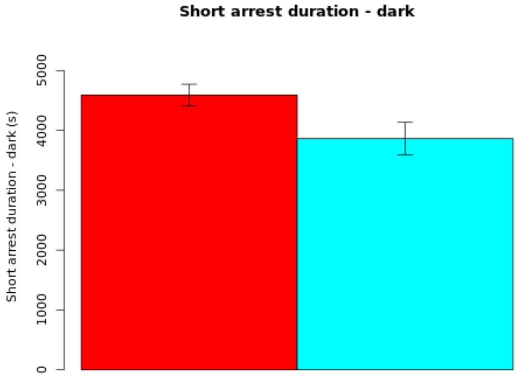

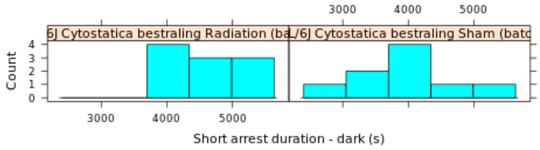

**Secondary plot:**

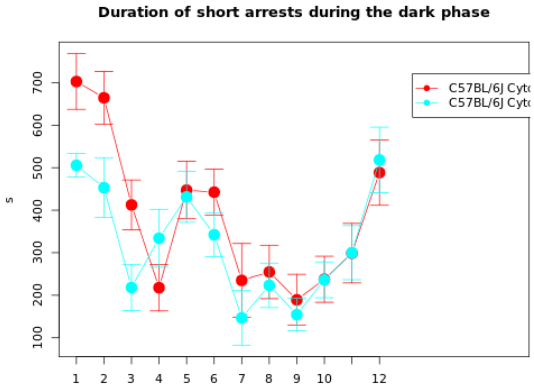

| Genotype | Mean | Median | SEM | N |
|---|---|---|---|---|
| (Gr 1): C57BL/6J Cytostatica bestraling Radiation (batch 22) | 4591.3053 | 4644.0867 | 179.8608 | 10 |
| (Gr 2): C57BL/6J Cytostatica bestraling Sham (batch 22) | 3863.76 | 3951.3867 | 270.4852 | 9 |

| | |
|---|---|
| **T-test:** | 2.2398 |
| **P-value:** | 0.0416 |
| **Df:** | 14.1753 |
| **Mann-Whitney U test:** | 0.0535 |

| Mice excluded by Quality Control |
|---|
| PH06469 (C57BL/6J - ) |

 © Sylics BV

Mean short arrest duration - dark

*Mean short arrest duration - dark*: Mean duration per short arrest during the dark phase

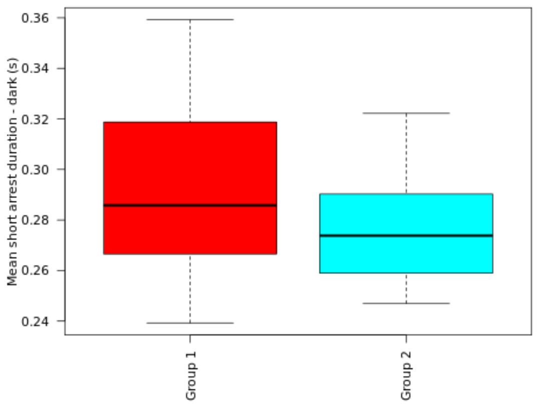

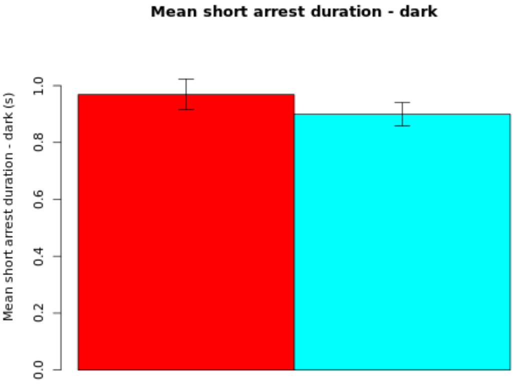

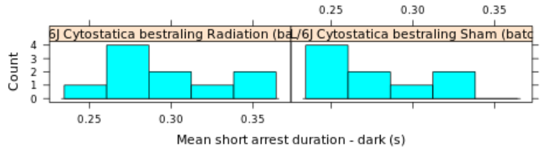

**NOTE: The statistics and boxplot are performed on log10 transformed data**

| Genotype | Mean | Median | SEM | N |
|---|---|---|---|---|
| (Gr 1): C57BL/6J Cytostatica bestraling Radiation (batch 22) | 0.9687 | 0 | Up: 0.0544<br>Down: 0.0529 | 10 |
| (Gr 2): C57BL/6J Cytostatica bestraling Sham (batch 22) | 0.8993 | 0 | Up: 0.0412<br>Down: 0.0403 | 9 |

| | |
|---|---|
| **T-test:** | 1.0336 |
| **P-value:** | 0.3163 |
| **Df:** | 16.4849 |
| **Mann-Whitney U test:** | 0.3562 |

| Mice excluded by Quality Control |
|---|
| PH06469 (C57BL/6J - ) |

© Sylics BV

Short arrest number - dark

*Short arrest number - dark*: Cummulative number of short arrests during the dark phase

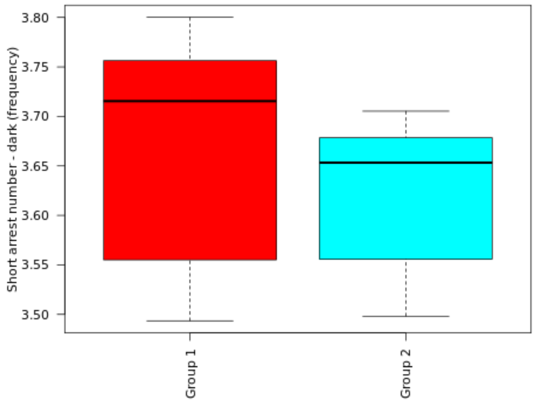

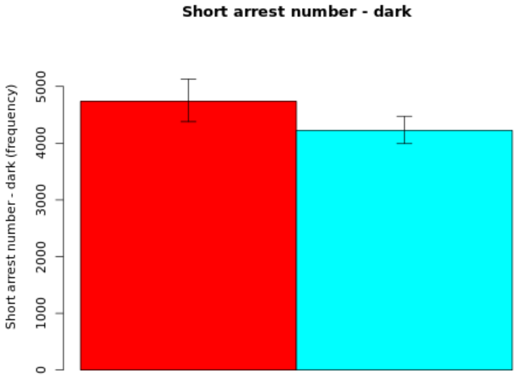

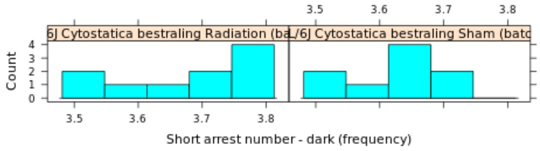

**Secondary plot:**

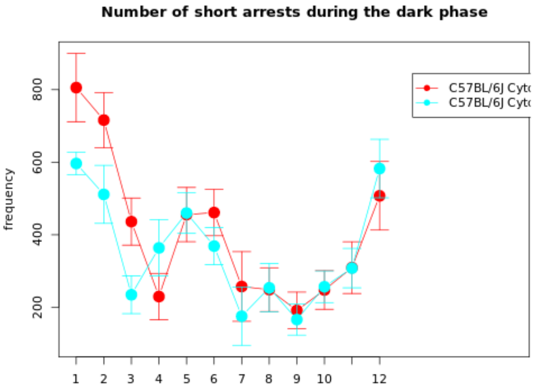

**NOTE: The statistics and boxplot are performed on log10 transformed data**

| Genotype | Mean | Median | SEM | N |
|---|---|---|---|---|
| (Gr 1): C57BL/6J Cytostatica bestraling Radiation (batch 22) | 4739.9883 | 0 | Up: 390.3067<br>Down: 360.6184 | 10 |
| (Gr 2): C57BL/6J Cytostatica bestraling Sham (batch 22) | 4226.0999 | 0 | Up: 248.4609<br>Down: 234.6676 | 9 |

| | |
|---|---|
| **T-test:** | 1.1758 |
| **P-value:** | 0.2569 |
| **Df:** | 15.9517 |
| **Mann-Whitney U test:** | 0.211 |

| Mice excluded by Quality Control |
|---|
| PH06469 (C57BL/6J - ) |

© Sylics BV

Long arrest duration - dark

*Long arrest duration - dark*: Cummulative duration of long arrests during the dark phase

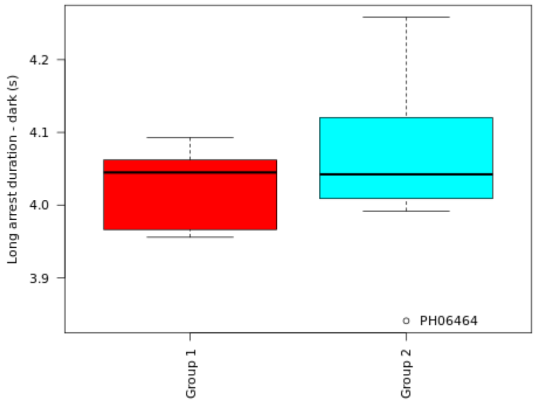

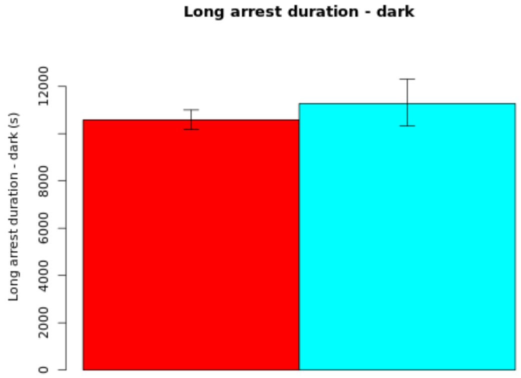

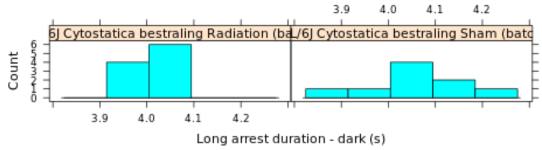

**Secondary plot:**

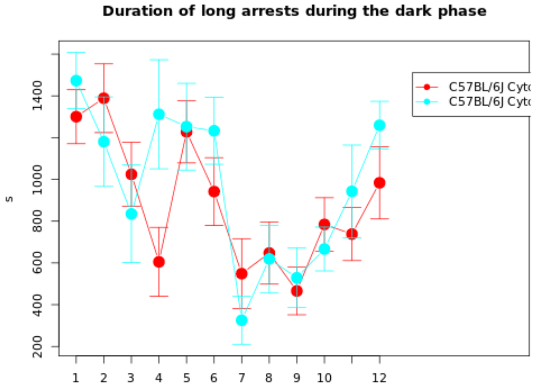

**NOTE: The statistics and boxplot are performed on log10 transformed data**

| Genotype | Mean | Median | SEM | N |
|---|---|---|---|---|
| (Gr 1): C57BL/6J Cytostatica bestraling Radiation (batch 22) | 10585.6425 | 0 | Up: 418.81<br>Down: 402.8722 | 10 |
| (Gr 2): C57BL/6J Cytostatica bestraling Sham (batch 22) | 11277.134 | 0 | Up: 1033.3233<br>Down: 946.5946 | 9 |

| | |
|---|---|
| **T-test:** | -0.66 |
| **P-value:** | 0.5228 |
| **Df:** | 11.0635 |
| **Mann-Whitney U test:** | 0.4967 |

| Mice excluded by Quality Control |
|---|
| PH06469 (C57BL/6J - ) |

 © Sylics BV

Mean long arrest duration - dark

*Mean long arrest duration - dark*: Mean duration per long arrest during the dark phase

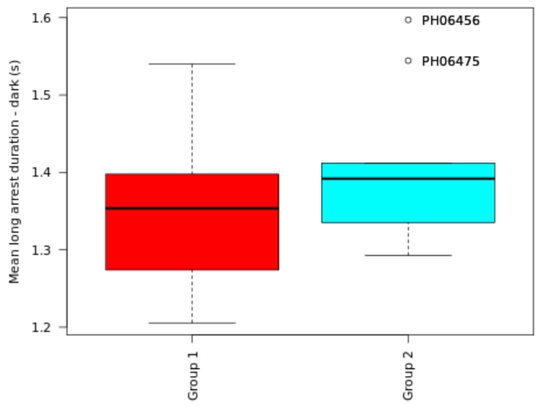

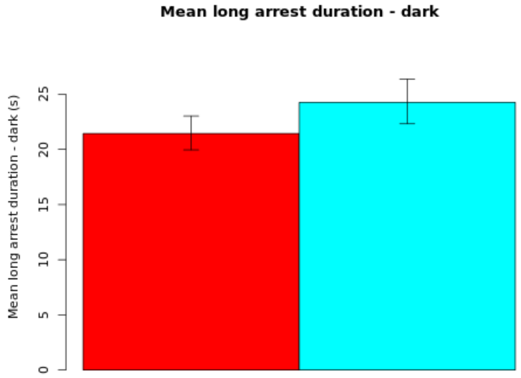

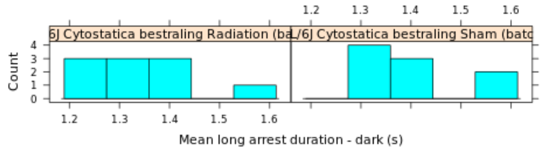

**NOTE: The statistics and boxplot are performed on log10 transformed data**

| Genotype | Mean | Median | SEM | N |
|---|---|---|---|---|
| (Gr 1): C57BL/6J Cytostatica bestraling Radiation (batch 22) | 21.4398 | 0 | Up: 1.5869<br>Down: 1.4821 | 10 |
| (Gr 2): C57BL/6J Cytostatica bestraling Sham (batch 22) | 24.2649 | 0 | Up: 2.1083<br>Down: 1.9459 | 9 |

| | |
|---|---|
| **T-test:** | -1.1259 |
| **P-value:** | 0.2766 |
| **Df:** | 16.2327 |
| **Mann-Whitney U test:** | 0.3562 |

| Mice excluded by Quality Control |
|---|
| PH06469 (C57BL/6J - ) |

 *© Sylics BV*

Long arrest number - dark

*Long arrest number - dark*: Cummulative number of long arrests during the dark phase

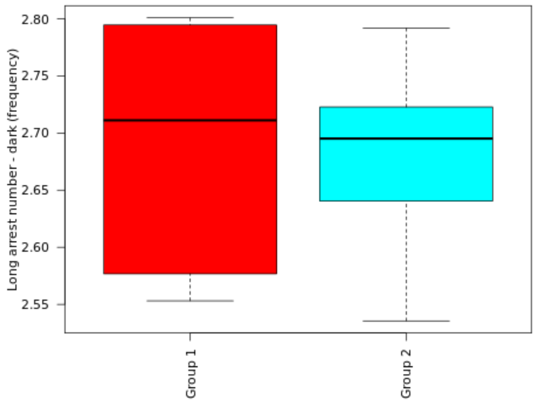

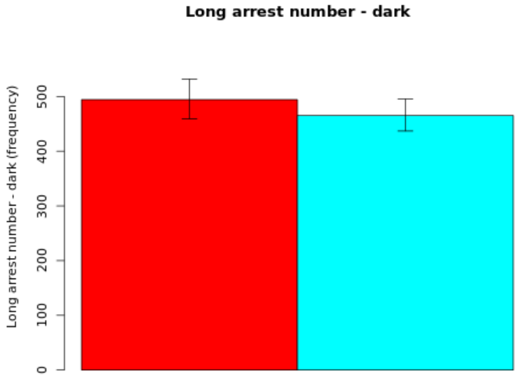

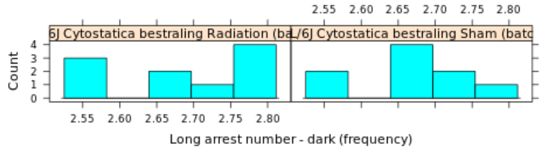

**Secondary plot:**

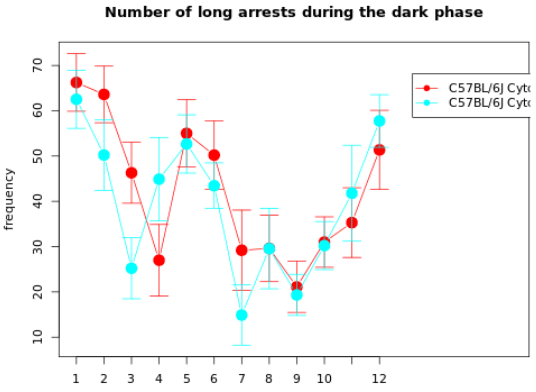

**NOTE: The statistics and boxplot are performed on log10 transformed data**

| Genotype | Mean | Median | SEM | N |
|---|---|---|---|---|
| (Gr 1): C57BL/6J Cytostatica bestraling Radiation (batch 22) | 494.2582 | 0 | Up: 37.5425<br>Down: 34.8972 | 10 |
| (Gr 2): C57BL/6J Cytostatica bestraling Sham (batch 22) | 465.2473 | 0 | Up: 30.1959<br>Down: 28.3593 | 9 |

| | |
|---|---|
| **T-test:** | 0.6267 |
| **P-value:** | 0.5392 |
| **Df:** | 16.8562 |
| **Mann-Whitney U test:** | 0.4002 |

| Mice excluded by Quality Control |
|---|
| PH06469 (C57BL/6J - ) |

 © Sylics BV

Long movement distance - dark

*Long movement distance - dark*: Cummulative long movement distance during the dark phase

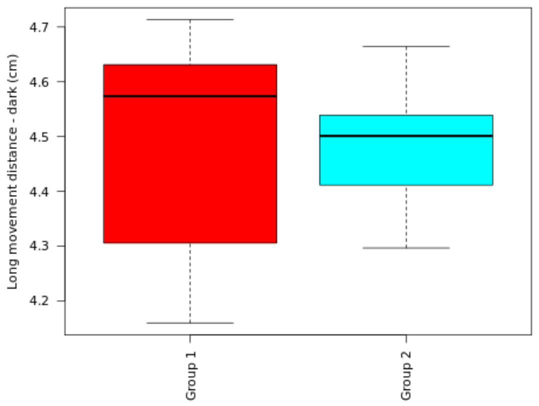

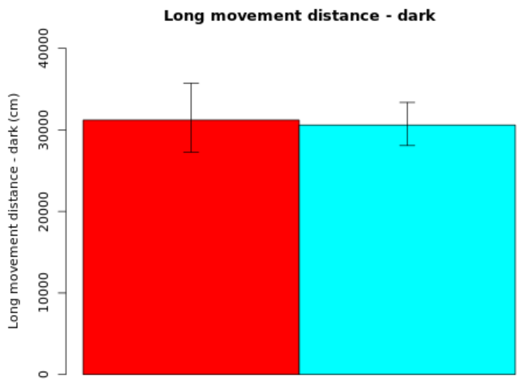

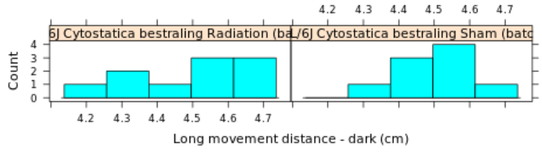

**Secondary plot:**

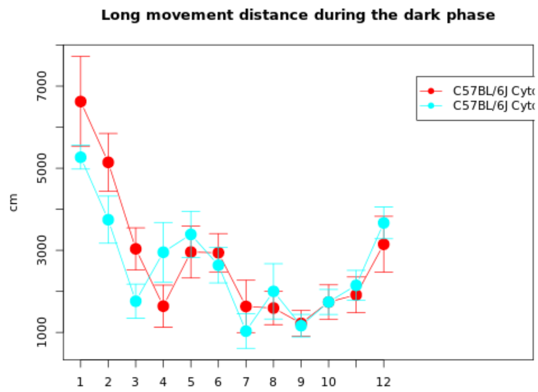

**NOTE: The statistics and boxplot are performed on log10 transformed data**

| Genotype | Mean | Median | SEM | N |
|---|---|---|---|---|
| (Gr 1): C57BL/6J Cytostatica bestraling Radiation (batch 22) | 31208.1139 | 0 | Up: 4514.75<br>Down: 3944.1799 | 10 |
| (Gr 2): C57BL/6J Cytostatica bestraling Sham (batch 22) | 30597.1743 | 0 | Up: 2755.4935<br>Down: 2527.85 | 9 |

| | |
|---|---|
| **T-test:** | 0.1233 |
| **P-value:** | 0.9035 |
| **Df:** | 15.0211 |
| **Mann-Whitney U test:** | 0.6038 |

| Mice excluded by Quality Control |
|---|
| PH06469 (C57BL/6J - ) |

Mean long movement distance - dark

*Mean long movement distance - dark*: Mean distance per long movement during the dark phase

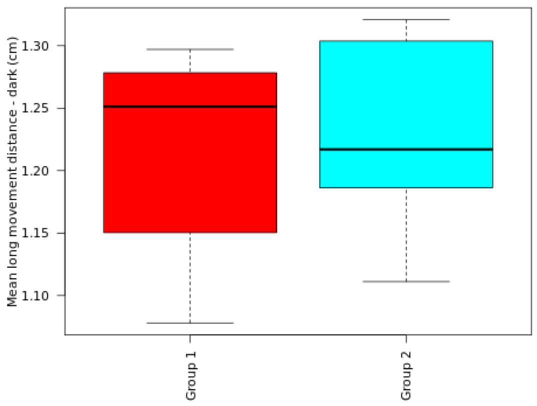

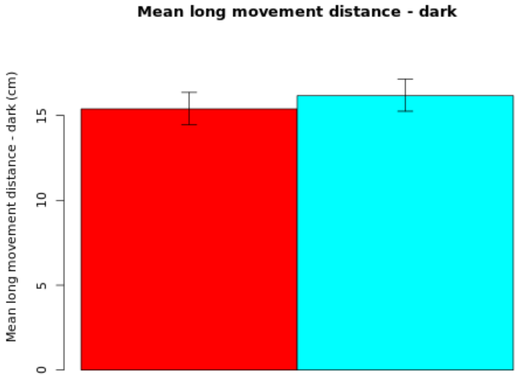

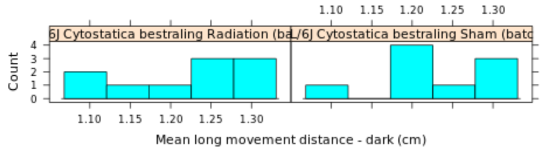

**NOTE: The statistics and boxplot are performed on log10 transformed data**

| Genotype | Mean | Median | SEM | N |
|---|---|---|---|---|
| (Gr 1): C57BL/6J Cytostatica bestraling Radiation (batch 22) | 15.3866 | 0 | Up: 0.9727<br>Down: 0.9182 | 10 |
| (Gr 2): C57BL/6J Cytostatica bestraling Sham (batch 22) | 16.1734 | 0 | Up: 0.9737<br>Down: 0.9215 | 9 |

| | |
|---|---|
| **T-test:** | -0.5878 |
| **P-value:** | 0.5644 |
| **Df:** | 16.9965 |
| **Mann-Whitney U test:** | 0.447 |

| Mice excluded by Quality Control |
|---|
| PH06469 (C57BL/6J - ) |

© Sylics BV

Long movement number - dark

*Long movement number - dark*: Cummulative long movement number during the dark phase

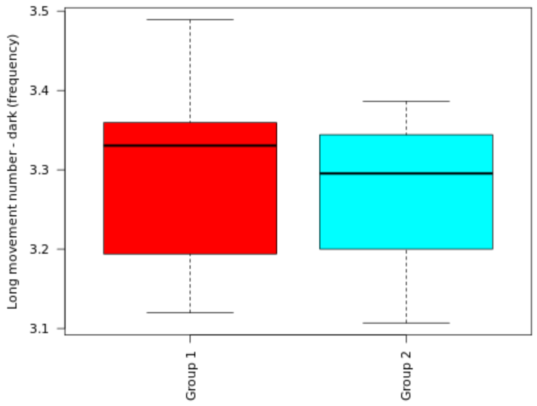

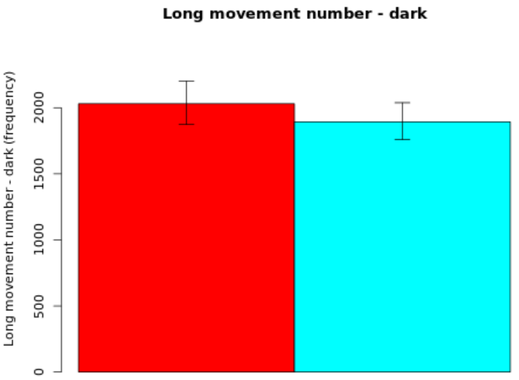

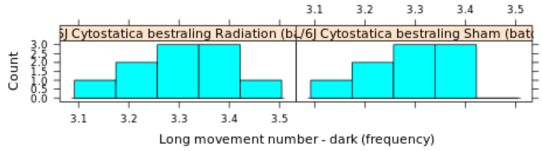

**Secondary plot:**

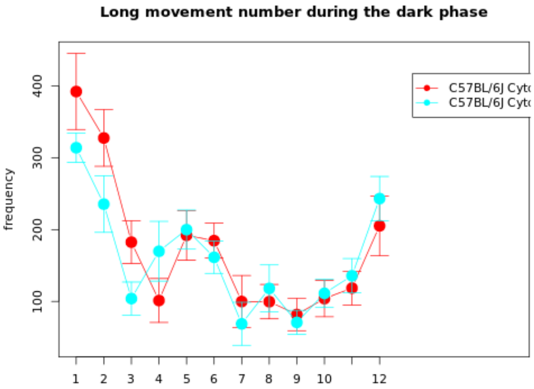

**NOTE: The statistics and boxplot are performed on log10 transformed data**

| Genotype | Mean | Median | SEM | N |
|---|---|---|---|---|
| (Gr 1): C57BL/6J Cytostatica bestraling Radiation (batch 22) | 2030.4924 | 0 | Up: 171.1397<br>Down: 157.8425 | 10 |
| (Gr 2): C57BL/6J Cytostatica bestraling Sham (batch 22) | 1893.3859 | 0 | Up: 144.394<br>Down: 134.1674 | 9 |

| | |
|---|---|
| **T-test:** | 0.6395 |
| **P-value:** | 0.531 |
| **Df:** | 16.9765 |
| **Mann-Whitney U test:** | 0.6038 |

| Mice excluded by Quality Control |
|---|
| PH06469 (C57BL/6J - ) |

 © Sylics BV

Short movement distance - dark

*Short movement distance - dark*: Cummulative short movement distance during the dark phase

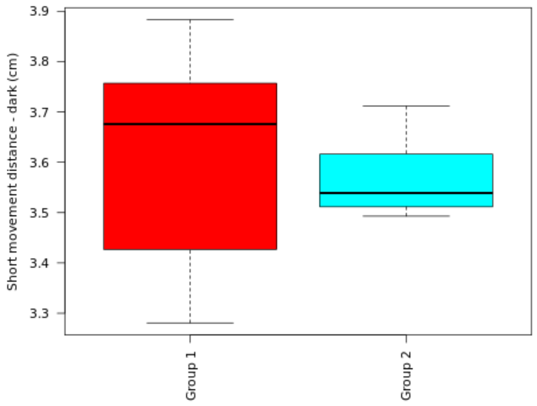

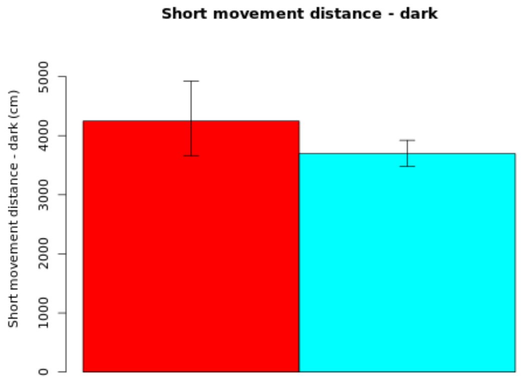

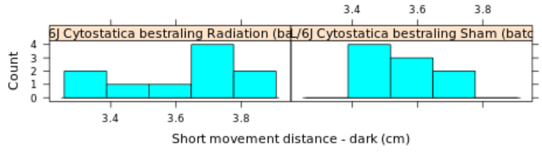

#### Secondary plot:

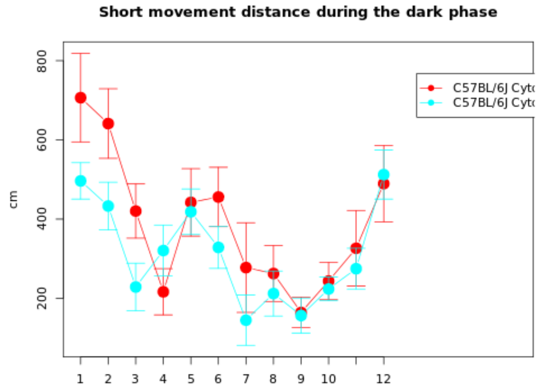

**NOTE: The statistics and boxplot are performed on log10 transformed data**

| Genotype | Mean | Median | SEM | N |
|---|---|---|---|---|
| (Gr 1): C57BL/6J Cytostatica bestraling Radiation (batch 22) | 4246.712 | 0 | Up: 676.8664<br>Down: 583.8334 | 10 |
| (Gr 2): C57BL/6J Cytostatica bestraling Sham (batch 22) | 3695.3427 | 0 | Up: 226.0568<br>Down: 213.0287 | 9 |

| | |
|---|---|
| **T-test:** | 0.8726 |
| **P-value:** | 0.4003 |
| **Df:** | 11.7904 |
| **Mann-Whitney U test:** | 0.3154 |

| Mice excluded by Quality Control |
|---|
| PH06469 (C57BL/6J - ) |

*© Sylics BV*

Mean short movement distance - dark

*Mean short movement distance - dark*: Mean distance per short movement during the dark phase

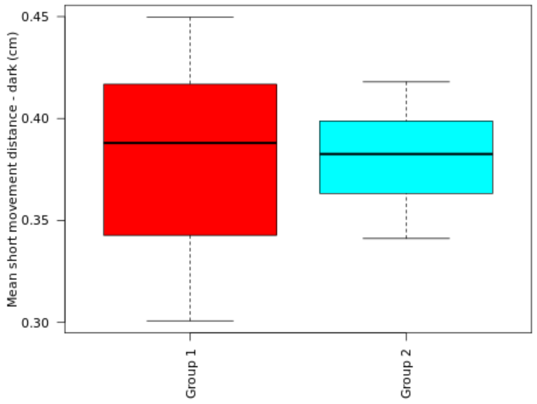

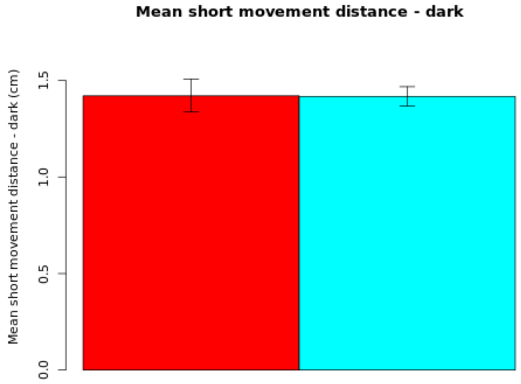

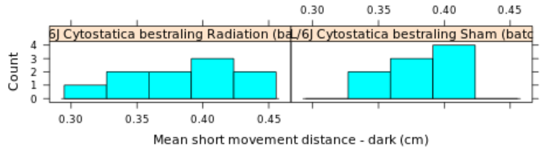

**NOTE: The statistics and boxplot are performed on log10 transformed data**

| Genotype | Mean | Median | SEM | N |
|---|---|---|---|---|
| (Gr 1): C57BL/6J Cytostatica bestraling Radiation (batch 22) | 1.4203 | 0 | Up: 0.0868<br>Down: 0.0838 | 10 |
| (Gr 2): C57BL/6J Cytostatica bestraling Sham (batch 22) | 1.417 | 0 | Up: 0.0509<br>Down: 0.0498 | 9 |

| | |
|---|---|
| **T-test:** | 0.0335 |
| **P-value:** | 0.9737 |
| **Df:** | 14.4112 |
| **Mann-Whitney U test:** | 0.9682 |

| Mice excluded by Quality Control |
|---|
| PH06469 (C57BL/6J - ) |

 © Sylics BV

Short movement number - dark

*Short movement number - dark*: Cummulative short movement number during the dark phase

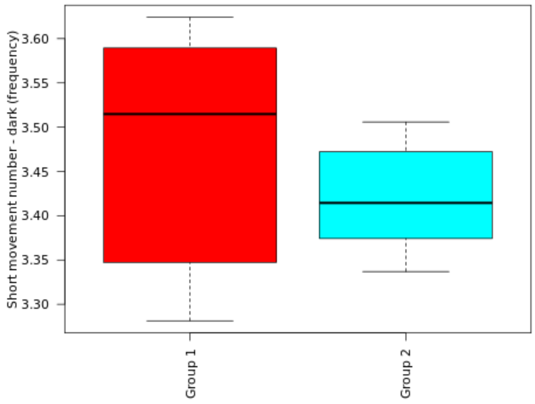

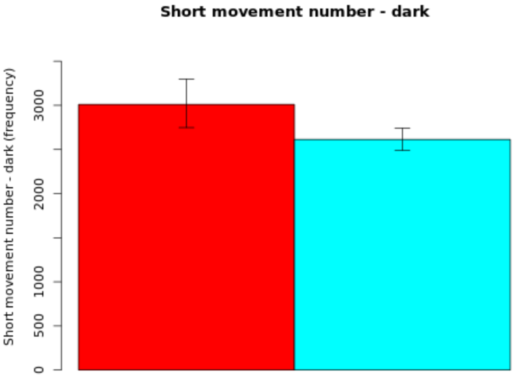

##### Secondary plot:

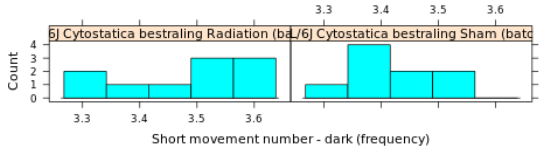

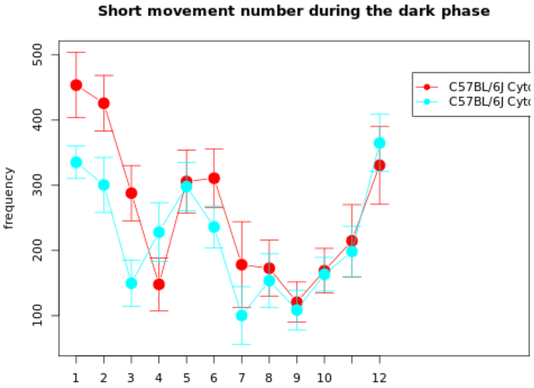

**NOTE: The statistics and boxplot are performed on log10 transformed data**

| Genotype | Mean | Median | SEM | N |
|---|---|---|---|---|
| (Gr 1): C57BL/6J Cytostatica bestraling Radiation (batch 22) | 3010.6219 | 0 | Up: 287.4255<br>Down: 262.3839 | 10 |
| (Gr 2): C57BL/6J Cytostatica bestraling Sham (batch 22) | 2613.3336 | 0 | Up: 129.3188<br>Down: 123.2235 | 9 |

| | |
|---|---|
| **T-test:** | 1.3715 |
| **P-value:** | 0.1925 |
| **Df:** | 13.5576 |
| **Mann-Whitney U test:** | 0.2428 |

| Mice excluded by Quality Control |
|---|
| PH06469 (C57BL/6J - ) |

 © Sylics BV

Short arrest duration - light

*Short arrest duration - light*: Cummulative duration of short arrests during the light phase

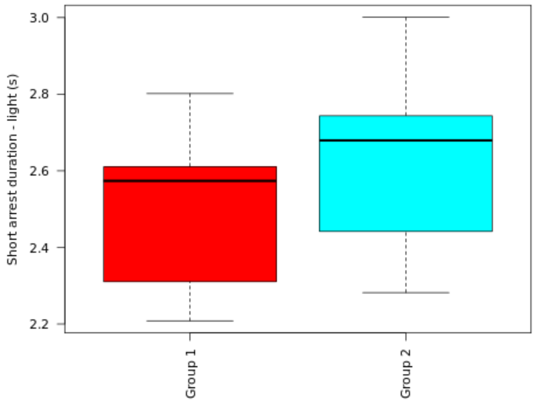

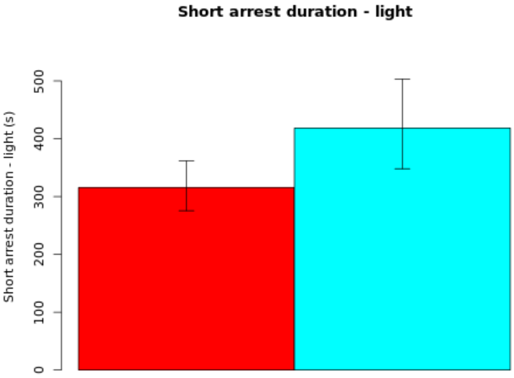

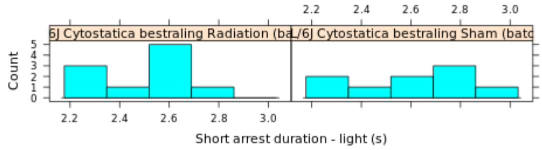

#### Secondary plot:

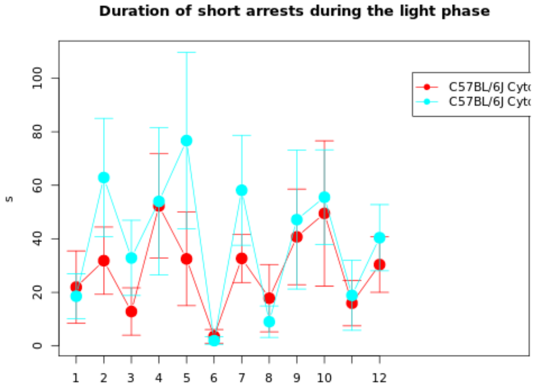

##### NOTE: The statistics and boxplot are performed on log10 transformed data

| Genotype | Mean | Median | SEM | N |
|---|---|---|---|---|
| (Gr 1): C57BL/6J Cytostatica bestraling Radiation (batch 22) | 315.7366 | 0 | Up: 46.0064<br>Down: 40.1714 | 10 |
| (Gr 2): C57BL/6J Cytostatica bestraling Sham (batch 22) | 418.545 | 0 | Up: 84.6444<br>Down: 70.4341 | 9 |

| | |
|---|---|
| **T-test:** | -1.2307 |
| **P-value:** | 0.2373 |
| **Df:** | 15.1042 |
| **Mann-Whitney U test:** | 0.1823 |

| Mice excluded by Quality Control |
|---|
| PH06469 (C57BL/6J - ) |

© Sylics BV

Mean short arrest duration - light

*Mean short arrest duration - light*: Mean duration per short arrest during the light phase

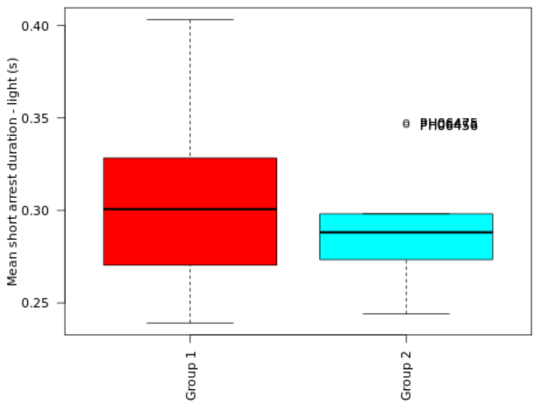

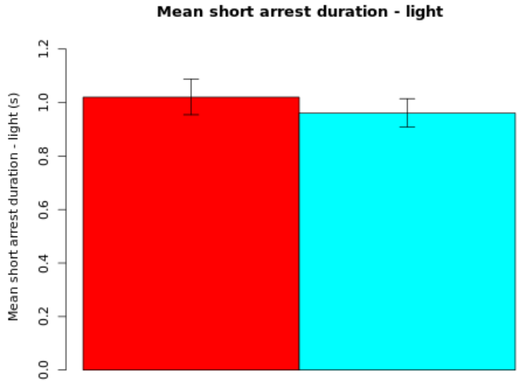

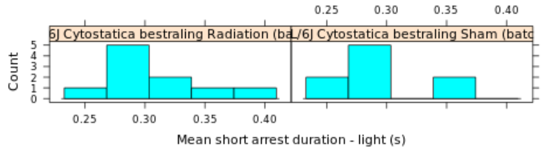

**NOTE: The statistics and boxplot are performed on log10 transformed data**

| Genotype | Mean | Median | SEM | N |
|---|---|---|---|---|
| (Gr 1): C57BL/6J Cytostatica bestraling Radiation (batch 22) | 1.0201 | 0 | Up: 0.0678<br>Down: 0.0656 | 10 |
| (Gr 2): C57BL/6J Cytostatica bestraling Sham (batch 22) | 0.9608 | 0 | Up: 0.0537<br>Down: 0.0523 | 9 |

| | |
|---|---|
| **T-test:** | 0.699 |
| **P-value:** | 0.4942 |
| **Df:** | 16.6834 |
| **Mann-Whitney U test:** | 0.4967 |

| Mice excluded by Quality Control |
|---|
| PH06469 (C57BL/6J - ) |

 *© Sylics BV*

Short arrest number - light

*Short arrest number - light*: Cummulative number of short arrests during the light phase

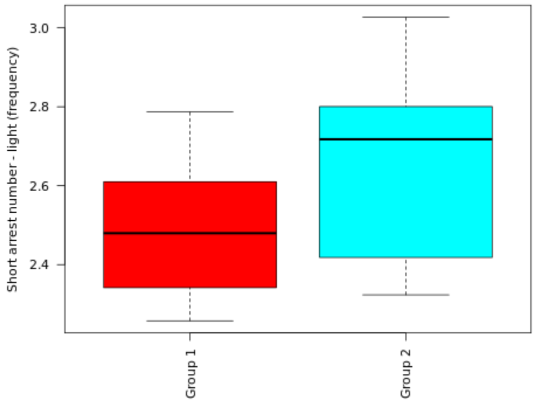

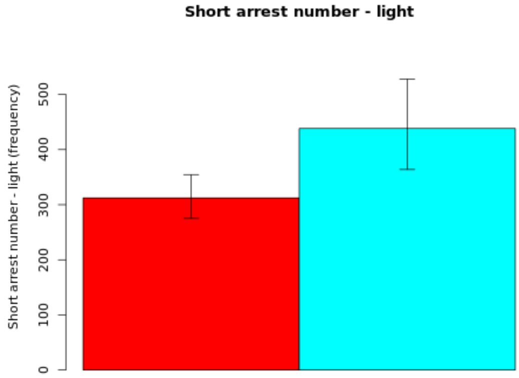

#### Secondary plot:

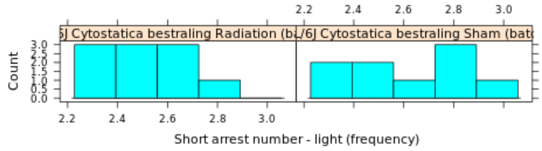

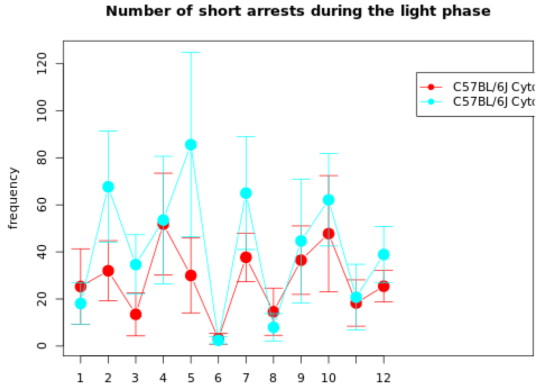

**NOTE: The statistics and boxplot are performed on log10 transformed data**

| Genotype | Mean | Median | SEM | N |
|---|---|---|---|---|
| (Gr 1): C57BL/6J Cytostatica bestraling Radiation (batch 22) | 312.3225 | 0 | Up: 42.2539<br>Down: 37.2327 | 10 |
| (Gr 2): C57BL/6J Cytostatica bestraling Sham (batch 22) | 438.2676 | 0 | Up: 89.6382<br>Down: 74.4464 | 9 |

| | |
|---|---|
| **T-test:** | -1.5037 |
| **P-value:** | 0.1543 |
| **Df:** | 14.3929 |
| **Mann-Whitney U test:** | 0.1564 |

| Mice excluded by Quality Control |
|---|
| PH06469 (C57BL/6J - ) |

© Sylics BV

Long arrest duration - light

*Long arrest duration - light*: Cummulative duration of long arrests during the light phase

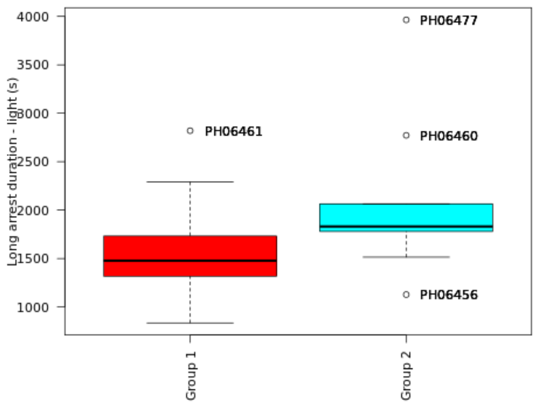

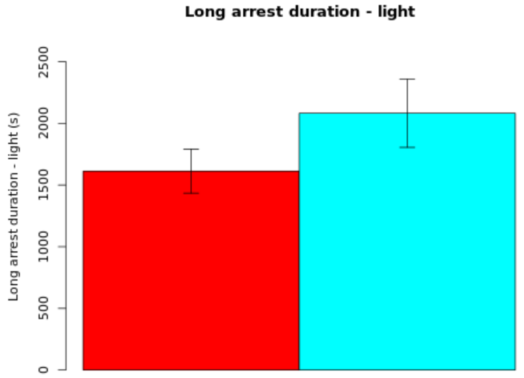

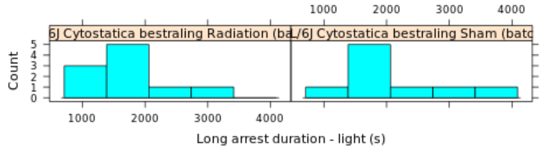

**Secondary plot:**

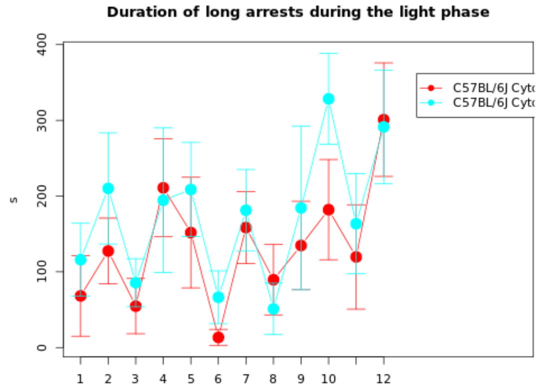

| Genotype | Mean | Median | SEM | N |
|---|---|---|---|---|
| (Gr 1): C57BL/6J Cytostatica bestraling Radiation (batch 22) | 1612.3427 | 1477.5333 | 178.5788 | 10 |
| (Gr 2): C57BL/6J Cytostatica bestraling Sham (batch 22) | 2081.8281 | 1830.5467 | 277.0152 | 9 |

| | |
|---|---|
| **T-test:** | -1.4245 |
| **P-value:** | 0.1764 |
| **Df:** | 13.8974 |
| **Mann-Whitney U test:** | 0.0947 |

| Mice excluded by Quality Control |
|---|
| PH06469 (C57BL/6J - ) |

Mean long arrest duration - light

*Mean long arrest duration - light*: Mean duration per long arrest during the light phase

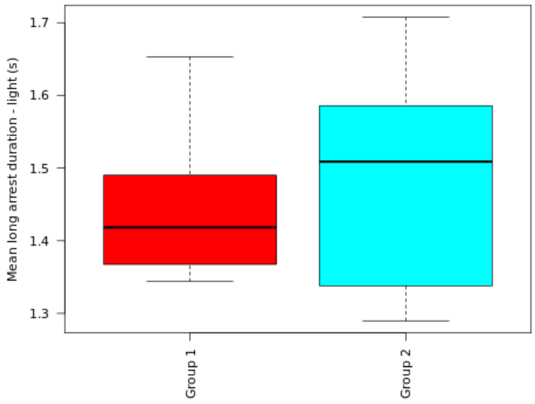

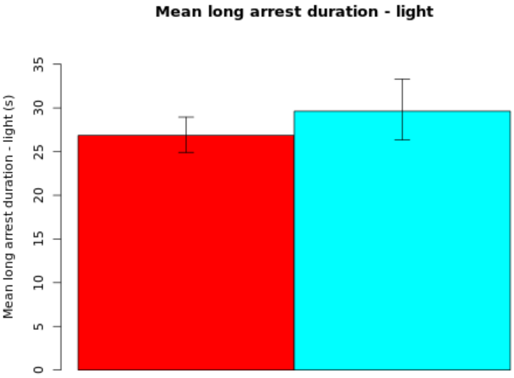

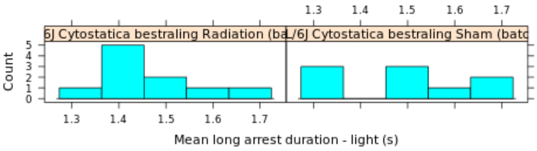

**NOTE: The statistics and boxplot are performed on log10 transformed data**

| Genotype | Mean | Median | SEM | N |
|---|---|---|---|---|
| (Gr 1): C57BL/6J Cytostatica bestraling Radiation (batch 22) | 26.8548 | 0 | Up: 2.0924<br>Down: 1.9462 | 10 |
| (Gr 2): C57BL/6J Cytostatica bestraling Sham (batch 22) | 29.624 | 0 | Up: 3.6772<br>Down: 3.283 | 9 |

| | |
|---|---|
| **T-test:** | -0.7044 |
| **P-value:** | 0.4929 |
| **Df:** | 13.8156 |
| **Mann-Whitney U test:** | 0.6607 |

| Mice excluded by Quality Control |
|---|
| PH06469 (C57BL/6J - ) |

*© Sylics BV*

Long arrest number - light

*Long arrest number - light*: Cummulative number of long arrests during the light phase

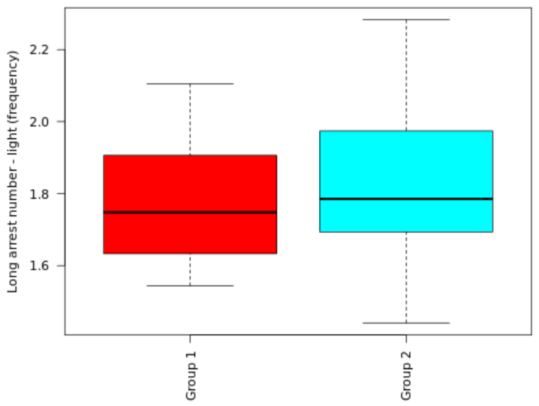

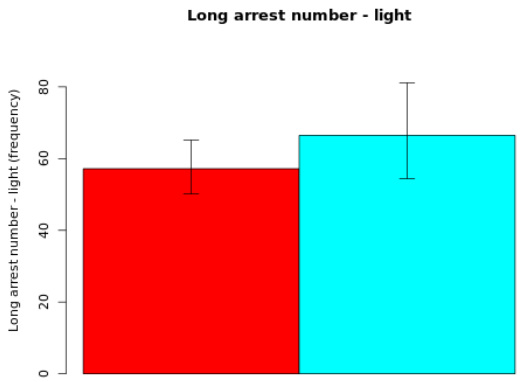

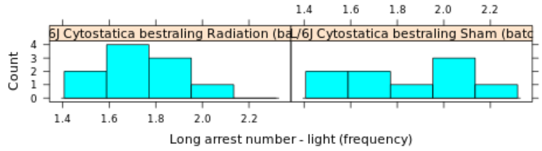

**Secondary plot:**

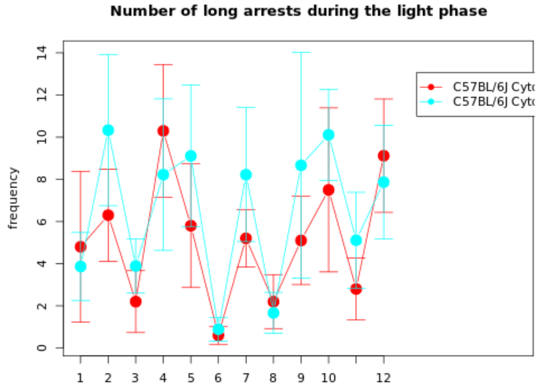

**NOTE: The statistics and boxplot are performed on log10 transformed data**

| Genotype | Mean | Median | SEM | N |
|---|---|---|---|---|
| (Gr 1): C57BL/6J Cytostatica bestraling Radiation (batch 22) | 57.1616 | 0 | Up: 7.9822<br>Down: 7.0189 | 10 |
| (Gr 2): C57BL/6J Cytostatica bestraling Sham (batch 22) | 66.4642 | 0 | Up: 14.657<br>Down: 12.041 | 9 |

| | |
|---|---|
| **T-test:** | -0.6316 |
| **P-value:** | 0.5378 |
| **Df:** | 14.0282 |
| **Mann-Whitney U test:** | 0.549 |

| Mice excluded by Quality Control |
|---|
| PH06469 (C57BL/6J - ) |

 © Sylics BV

Long movement distance - light

*Long movement distance - light*: Cummulative long movement distance during the light phase

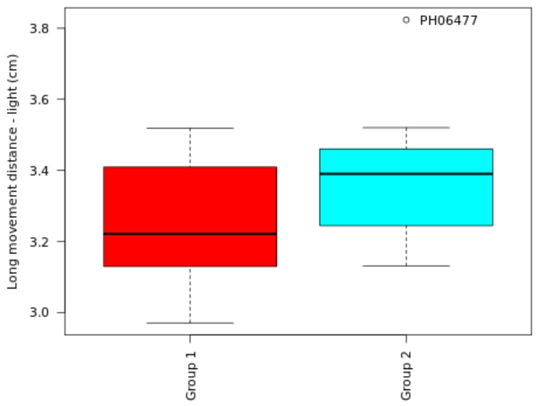

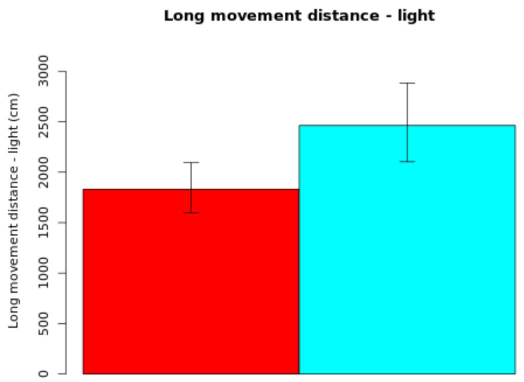

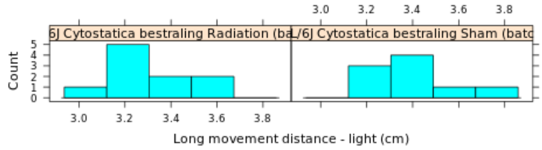

**Secondary plot:**

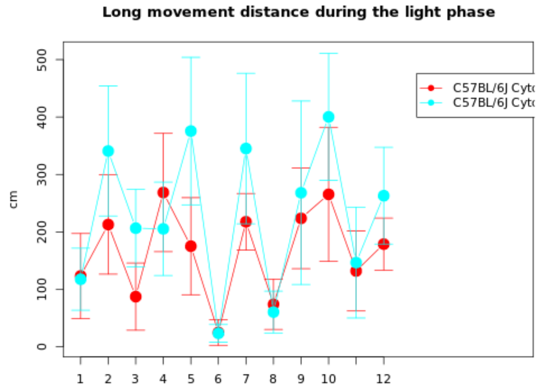

**NOTE: The statistics and boxplot are performed on log10 transformed data**

| Genotype | Mean | Median | SEM | N |
|---|---|---|---|---|
| (Gr 1): C57BL/6J Cytostatica bestraling Radiation (batch 22) | 1830.2733 | 0 | Up: 263.7389<br>Down: 230.5371 | 10 |
| (Gr 2): C57BL/6J Cytostatica bestraling Sham (batch 22) | 2462.4619 | 0 | Up: 420.3746<br>Down: 359.0969 | 9 |

| | |
|---|---|
| **T-test:** | -1.4313 |
| **P-value:** | 0.1713 |
| **Df:** | 16.2439 |
| **Mann-Whitney U test:** | 0.211 |

| Mice excluded by Quality Control |
|---|
| PH06469 (C57BL/6J - ) |

*© Sylics BV*

Mean long movement distance - light

*Mean long movement distance - light*: Mean distance per long movement during the light phase

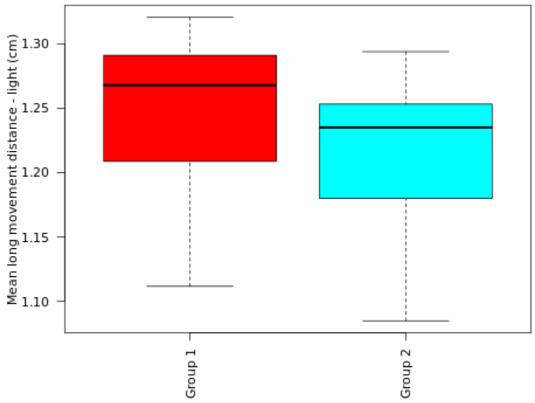

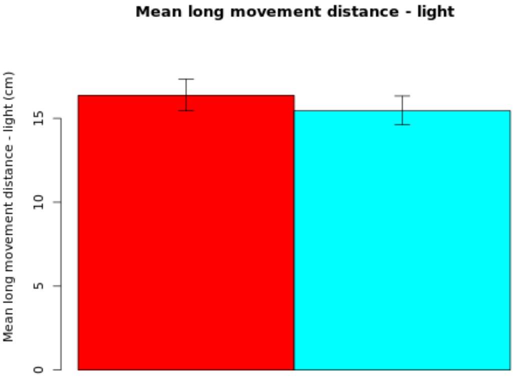

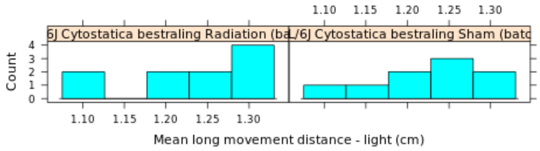

**NOTE: The statistics and boxplot are performed on log10 transformed data**

| Genotype | Mean | Median | SEM | N |
|---|---|---|---|---|
| (Gr 1): C57BL/6J Cytostatica bestraling Radiation (batch 22) | 16.3714 | 0 | Up: 0.9675<br>Down: 0.9165 | 10 |
| (Gr 2): C57BL/6J Cytostatica bestraling Sham (batch 22) | 15.4528 | 0 | Up: 0.876<br>Down: 0.8317 | 9 |

| | |
|---|---|
| **T-test:** | 0.7241 |
| **P-value:** | 0.4788 |
| **Df:** | 16.9962 |
| **Mann-Whitney U test:** | 0.4002 |

| Mice excluded by Quality Control |
|---|
| PH06469 (C57BL/6J - ) |

 © Sylics BV

Long movement number - light

*Long movement number - light*: Cummulative long movement number during the light phase

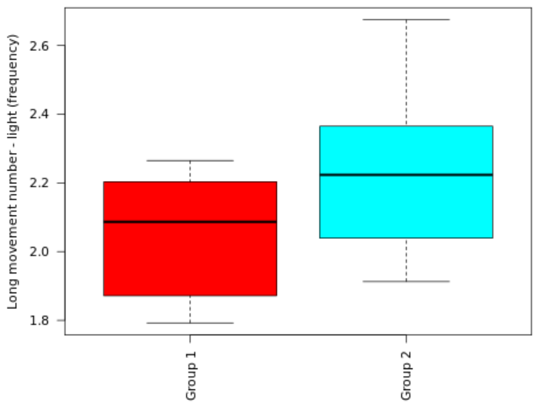

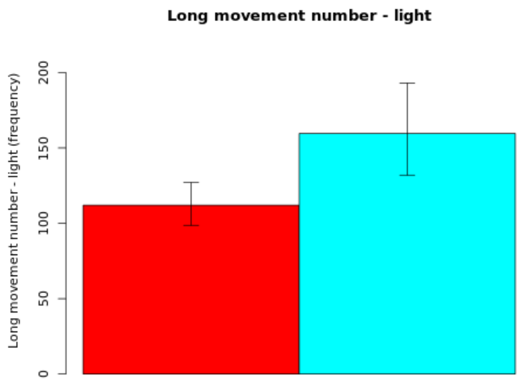

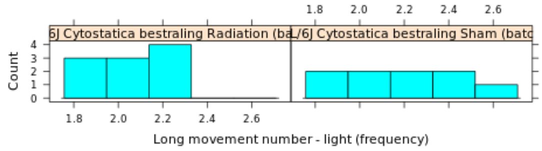

**Secondary plot:**

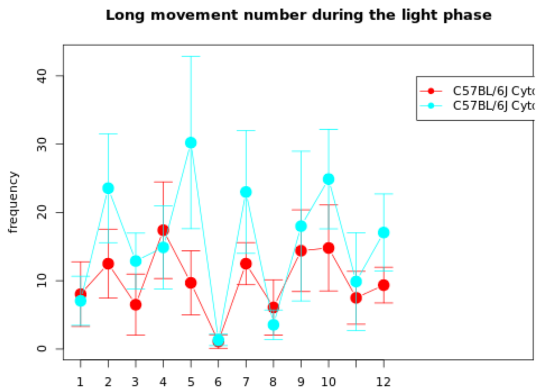

**NOTE: The statistics and boxplot are performed on log10 transformed data**

| Genotype | Mean | Median | SEM | N |
|---|---|---|---|---|
| (Gr 1): C57BL/6J Cytostatica bestraling Radiation (batch 22) | 111.9676 | 0 | Up: 15.0894<br>Down: 13.3113 | 10 |
| (Gr 2): C57BL/6J Cytostatica bestraling Sham (batch 22) | 159.6078 | 0 | Up: 33.4067<br>Down: 27.6545 | 9 |

| | |
|---|---|
| **T-test:** | -1.5516 |
| **P-value:** | 0.1428 |
| **Df:** | 14.1552 |
| **Mann-Whitney U test:** | 0.211 |

| Mice excluded by Quality Control |
|---|
| PH06469 (C57BL/6J - ) |

 © Sylics BV

Short movement distance - light

*Short movement distance - light*: Cummulative short movement distance during the light phase

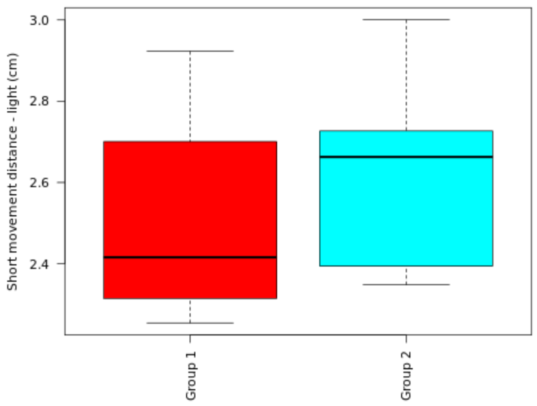

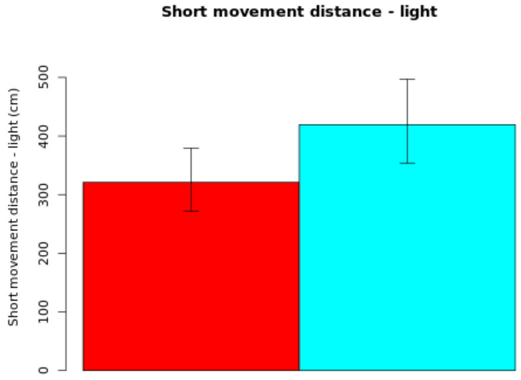

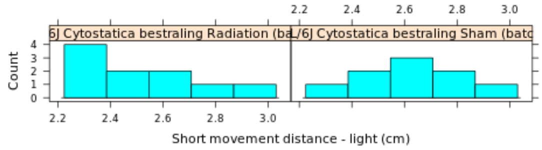

**Secondary plot:**

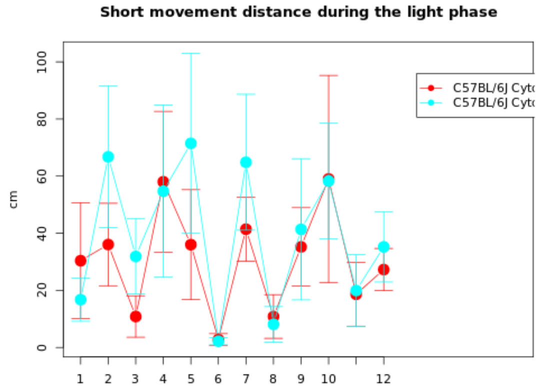

**NOTE: The statistics and boxplot are performed on log10 transformed data**

| Genotype | Mean | Median | SEM | N |
|---|---|---|---|---|
| (Gr 1): C57BL/6J Cytostatica bestraling Radiation (batch 22) | 321.3384 | 0 | Up: 58.1678<br>Down: 49.2758 | 10 |
| (Gr 2): C57BL/6J Cytostatica bestraling Sham (batch 22) | 419.3884 | 0 | Up: 77.6147<br>Down: 65.5183 | 9 |

| | |
|---|---|
| **T-test:** | -1.12 |
| **P-value:** | 0.2784 |
| **Df:** | 16.8919 |
| **Mann-Whitney U test:** | 0.3562 |

| Mice excluded by Quality Control |
|---|
| PH06469 (C57BL/6J - ) |

 © Sylics BV

Mean short movement distance - light

*Mean short movement distance - light*: Mean distance per short movement during the light phase

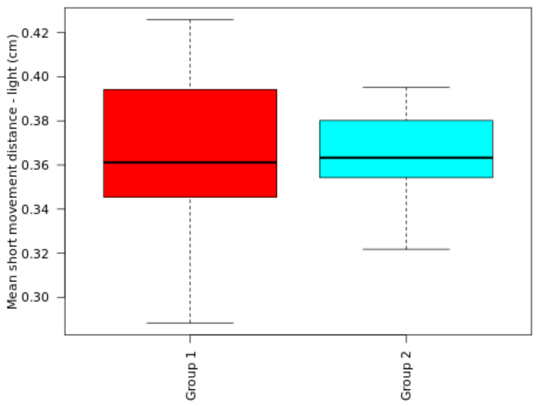

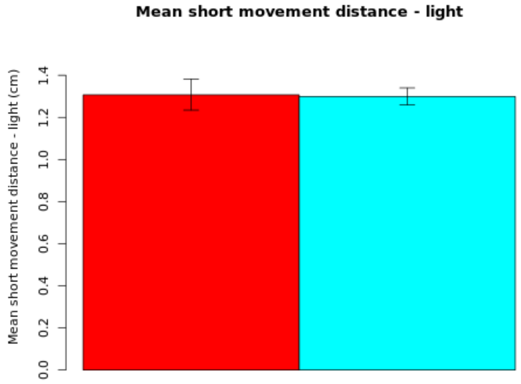

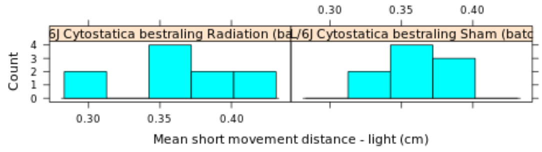

**NOTE: The statistics and boxplot are performed on log10 transformed data**

| Genotype | Mean | Median | SEM | N |
|---|---|---|---|---|
| (Gr 1): C57BL/6J Cytostatica bestraling Radiation (batch 22) | 1.307 | 0 | Up: 0.075<br>Down: 0.0726 | 10 |
| (Gr 2): C57BL/6J Cytostatica bestraling Sham (batch 22) | 1.2992 | 0 | Up: 0.0407<br>Down: 0.04 | 9 |

| | |
|---|---|
| **T-test:** | 0.0934 |
| **P-value:** | 0.9269 |
| **Df:** | 13.8358 |
| **Mann-Whitney U test:** | 0.8421 |

| Mice excluded by Quality Control |
|---|
| PH06469 (C57BL/6J - ) |

*© Sylics BV*

Short movement number - light

*Short movement number - light*: Cummulative short movement number during the light phase

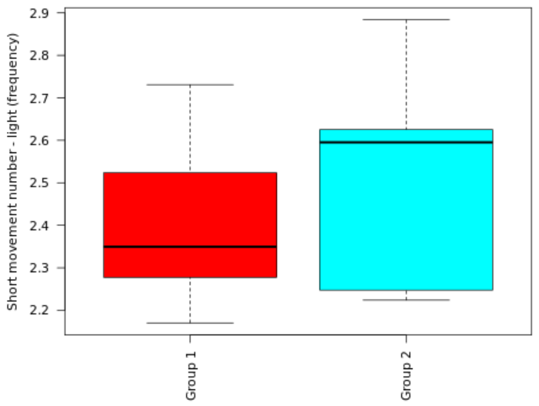

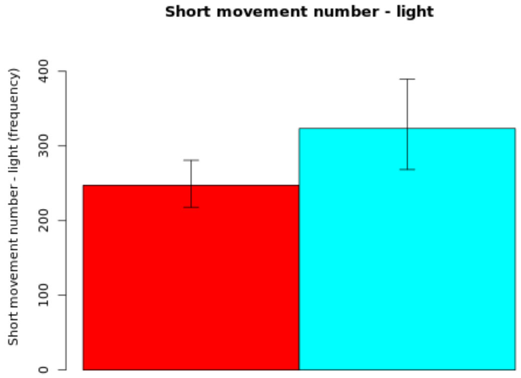

#### Secondary plot:

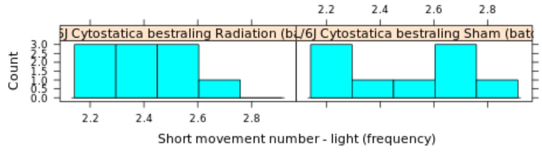

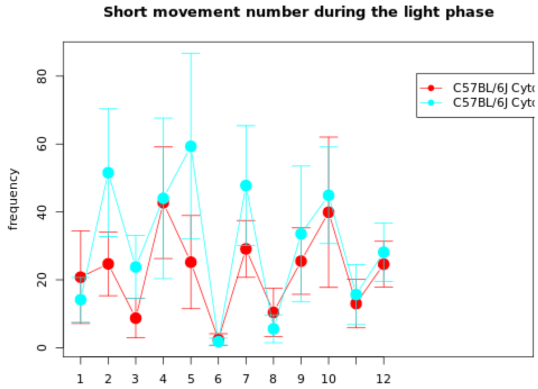

**NOTE: The statistics and boxplot are performed on log10 transformed data**

| Genotype | Mean | Median | SEM | N |
|---|---|---|---|---|
| (Gr 1): C57BL/6J Cytostatica bestraling Radiation (batch 22) | 247.4423 | 0 | Up: 33.4069<br>Down: 29.4473 | 10 |
| (Gr 2): C57BL/6J Cytostatica bestraling Sham (batch 22) | 323.4237 | 0 | Up: 66.179<br>Down: 54.9664 | 9 |

| | |
|---|---|
| **T-test:** | -1.1888 |
| **P-value:** | 0.2538 |
| **Df:** | 14.3713 |
| **Mann-Whitney U test:** | 0.2775 |

| Mice excluded by Quality Control |
|---|
| PH06469 (C57BL/6J - ) |

 © Sylics BV

Long movement threshold

*Long movement threshold*: Cut-off value to separate short and long movements

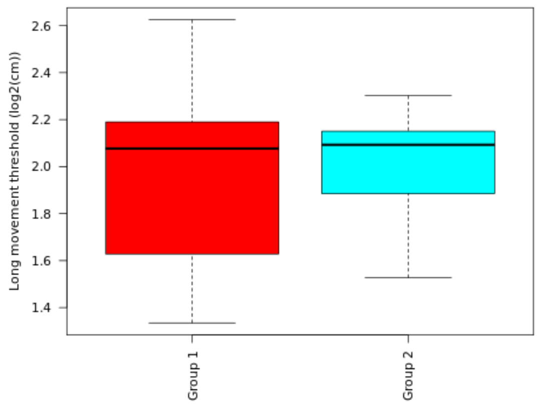

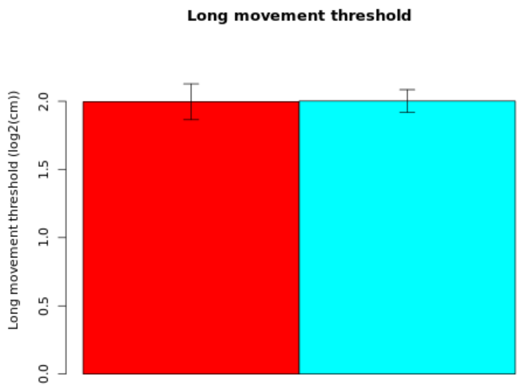

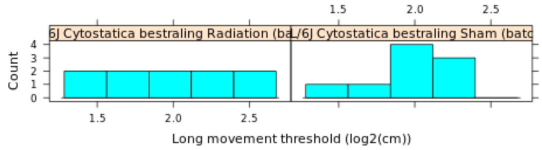

**Secondary plot:**

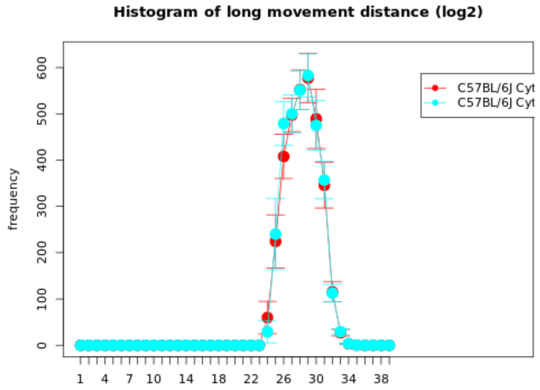

| Genotype | Mean | Median | SEM | N |
|---|---|---|---|---|
| (Gr 1): C57BL/6J Cytostatica bestraling Radiation (batch 22) | 1.9961 | 2.0769 | 0.1314 | 10 |
| (Gr 2): C57BL/6J Cytostatica bestraling Sham (batch 22) | 2.0018 | 2.0927 | 0.0833 | 9 |

| | |
|---|---|
| **T-test:** | -0.0369 |
| **P-value:** | 0.9711 |
| **Df:** | 14.9713 |
| **Mann-Whitney U test:** | 0.9048 |

| Mice excluded by Quality Control |
|---|
| PH06469 (C57BL/6J - ) |

 © Sylics BV

### Spontaneous behavior (Group 2: sheltering)

With respect to spontaneous behaviors in the first three days in the PhenoTyper, 6 groups of behavioral parameters are defined as described below. The first two groups describe specific behavioral elements related to kinematics of mice (description of movement characteristics, group1) and sheltering behavior (group 2). These behavioral parameters were analyzed with respect to temporal aspects, in particular over 4 different time scales, i.e., habituation effects across multiple days (group 3), effects of DarkLight phase across 24h (group 4), differences in the pattern of behavior in the few hours before and after phase shifts (group5), and differences in activity bout properties on the sub-minute time scale (group 6).

**Group 2: Sheltering behavior**
Mice frequently visit the shelter for a few seconds (i.e., passing through) during bouts of activity. In contrast, long shelter visits during which mice appeared to be resting or sleeping, are in the range of hours. Although long shelter visits were infrequent, they appeared as a separate class of events when the frequency distribution of shelter visit durations of the three days in the cage was plotted. Long shelter visits were readily identified by Gaussian mixture model fitting of shelter visit data of individual mice. The 90th percentile of the first fitted Gaussian was used as upper threshold to consistently distinguish short shelter visits.
See for instance the "histogram of shelter visit durations" for C57BL/6J mice at the bottom of this page:
http://mousedata.sylics.com/?page=strainhtp&loaded=true&s=11&htpid=114&e=10&batch=1&t=&texp=&list=batches&tdose=

 © Sylics BV

Short shelter visit duration - dark

*Short shelter visit duration - dark*: Cummulative duration of short shelter visits during the dark phase

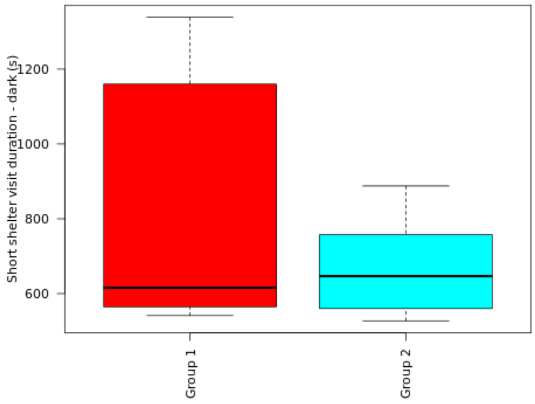

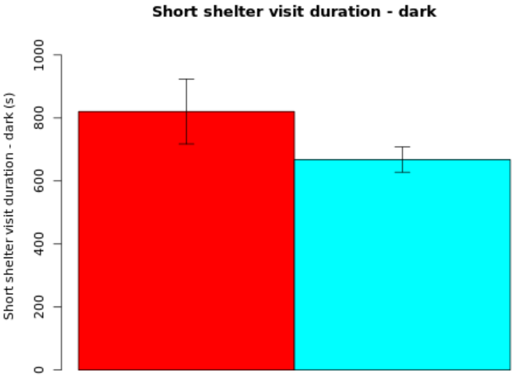

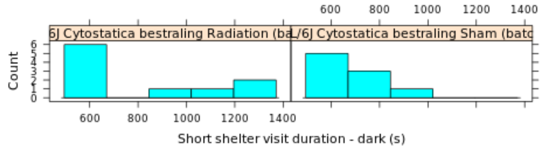

**Secondary plot:**

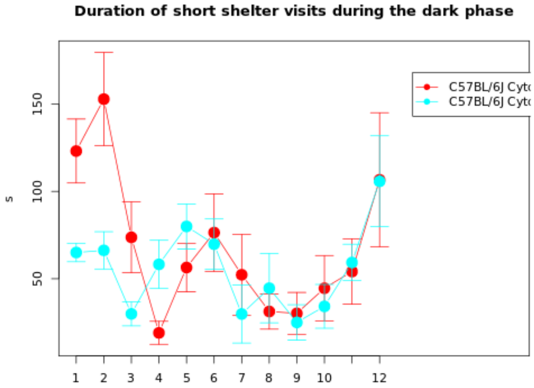

| Genotype | Mean | Median | SEM | N |
|---|---|---|---|---|
| (Gr 1): C57BL/6J Cytostatica bestraling Radiation (batch 22) | 820.6427 | 615.5 | 103.0038 | 10 |
| (Gr 2): C57BL/6J Cytostatica bestraling Sham (batch 22) | 667.9156 | 646.6267 | 39.9173 | 9 |

| | |
|---|---|
| **T-test:** | 1.3825 |
| **P-value:** | 0.1928 |
| **Df:** | 11.6116 |
| **Mann-Whitney U test:** | 0.6038 |

| Mice excluded by Quality Control |
|---|
| PH06469 (C57BL/6J - ) |

 © Sylics BV

Mean short shelter visit duration - dark

*Mean short shelter visit duration - dark*: Mean duration per short shelter visit during the dark phase

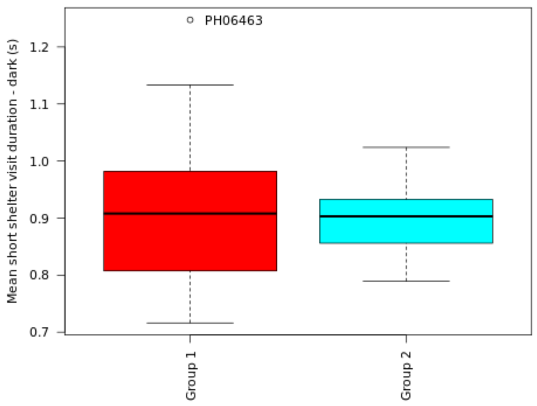

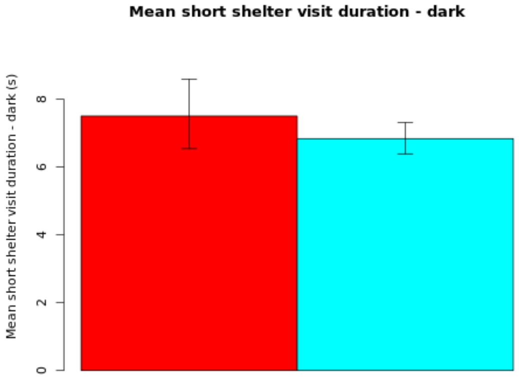

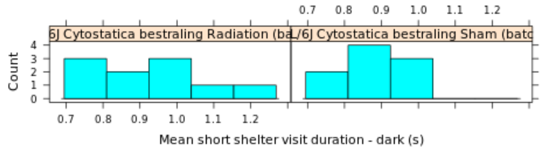

**NOTE: The statistics and boxplot are performed on log10 transformed data**

| Genotype | Mean | Median | SEM | N |
|---|---|---|---|---|
| (Gr 1): C57BL/6J Cytostatica bestraling Radiation (batch 22) | 7.5083 | 0 | Up: 1.0868<br>Down: 0.9637 | 10 |
| (Gr 2): C57BL/6J Cytostatica bestraling Sham (batch 22) | 6.8424 | 0 | Up: 0.4759<br>Down: 0.4487 | 9 |

| | |
|---|---|
| **T-test:** | 0.6087 |
| **P-value:** | 0.5532 |
| **Df:** | 12.999 |
| **Mann-Whitney U test:** | 0.7802 |

| Mice excluded by Quality Control |
|---|
| PH06469 (C57BL/6J - ) |

 © Sylics BV

Short shelter visit number - dark

*Short shelter visit number - dark*: Cummulative number of short shelter visits during the dark phase

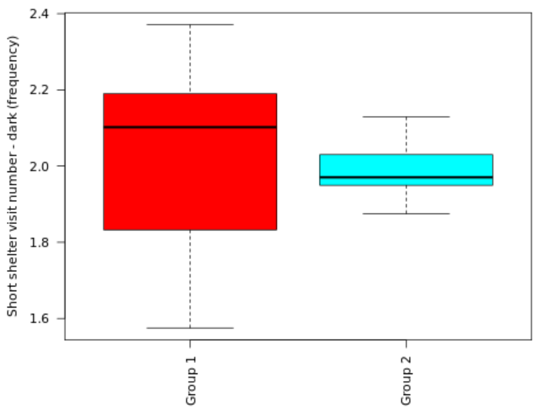

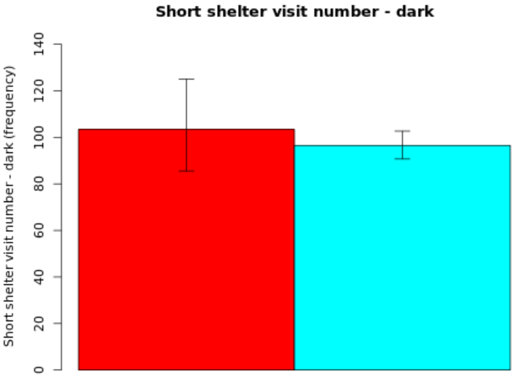

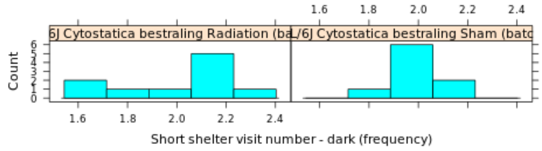

**Secondary plot:**

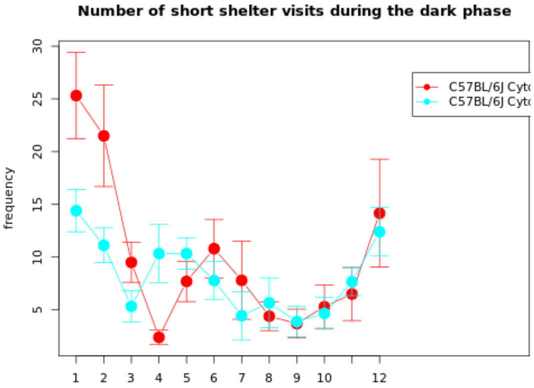

**NOTE: The statistics and boxplot are performed on log10 transformed data**

| Genotype | Mean | Median | SEM | N |
|---|---|---|---|---|
| (Gr 1): C57BL/6J Cytostatica bestraling Radiation (batch 22) | 103.4412 | 0 | Up: 21.5864<br>Down: 17.889 | 10 |
| (Gr 2): C57BL/6J Cytostatica bestraling Sham (batch 22) | 96.5041 | 0 | Up: 6.1705<br>Down: 5.8033 | 9 |

| | |
|---|---|
| **T-test:** | 0.3477 |
| **P-value:** | 0.7347 |
| **Df:** | 10.8833 |
| **Mann-Whitney U test:** | 0.3154 |

| Mice excluded by Quality Control |
|---|
| PH06469 (C57BL/6J - ) |

 *© Sylics BV*

#### Long shelter visit duration - dark

*Long shelter visit duration - dark*: Cummulative duration of long shelter visits during the dark phase

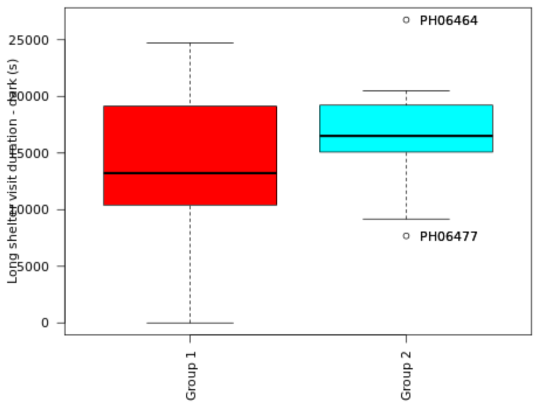

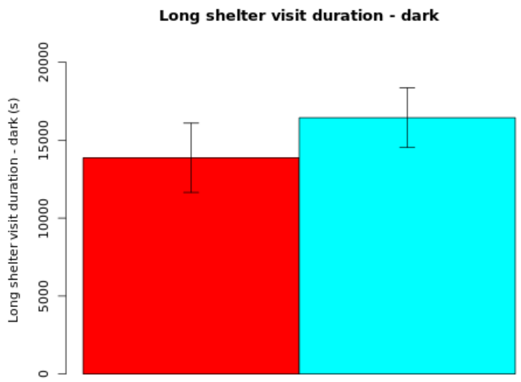

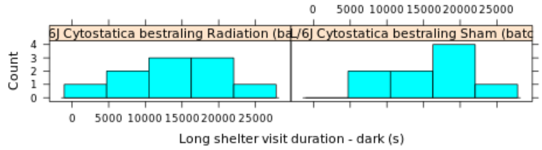

**Secondary plot:**

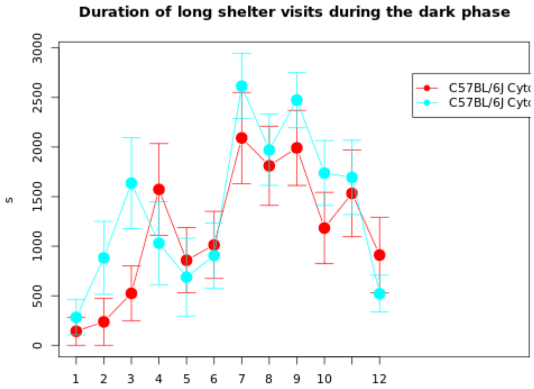

| Genotype | Mean | Median | SEM | N |
|---|---|---|---|---|
| (Gr 1): C57BL/6J Cytostatica bestraling Radiation (batch 22) | 13868.9867 | 13227.8 | 2220.3537 | 10 |
| (Gr 2): C57BL/6J Cytostatica bestraling Sham (batch 22) | 16440.1911 | 16510.3067 | 1915.5607 | 9 |

| | |
|---|---|
| **T-test:** | -0.8768 |
| **P-value:** | 0.3929 |
| **Df:** | 16.8696 |
| **Mann-Whitney U test:** | 0.4967 |

| Mice excluded by Quality Control |
|---|
| PH06469 (C57BL/6J - ) |

 © Sylics BV

Long shelter visit number - dark

*Long shelter visit number - dark*: Cummulative number of long shelter visits during the dark phase

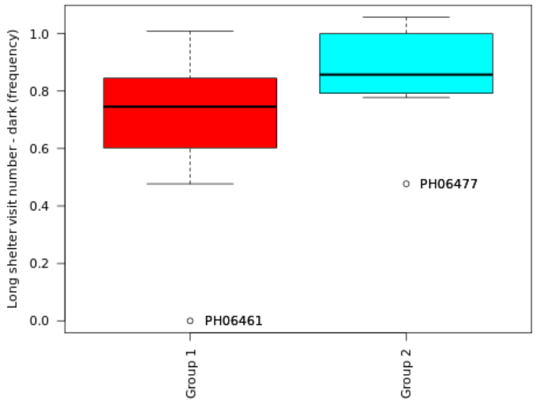

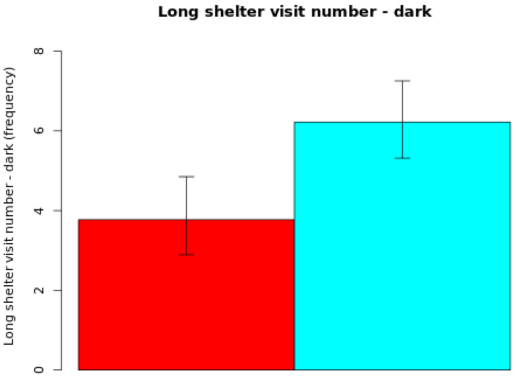

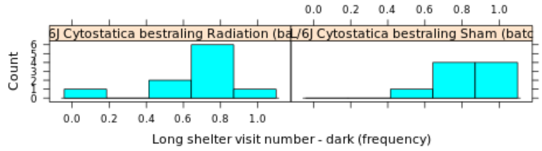

#### Secondary plot:

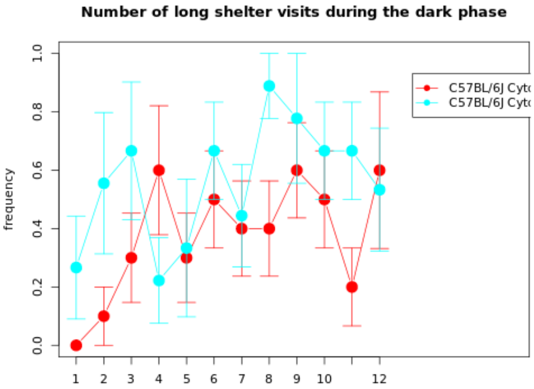

**NOTE: The statistics and boxplot are performed on log10 transformed data**

| Genotype | Mean | Median | SEM | N |
|---|---|---|---|---|
| (Gr 1): C57BL/6J Cytostatica bestraling Radiation (batch 22) | 3.7717 | 0 | Up: 1.0791<br>Down: 0.8801 | 10 |
| (Gr 2): C57BL/6J Cytostatica bestraling Sham (batch 22) | 6.2205 | 0 | Up: 1.0334<br>Down: 0.904 | 9 |

| | |
|---|---|
| **T-test:** | -1.6988 |
| **P-value:** | 0.1097 |
| **Df:** | 15.239 |
| **Mann-Whitney U test:** | 0.0934 |

| Mice excluded by Quality Control |
|---|
| PH06469 (C57BL/6J - ) |

*© Sylics BV*

Mean long shelter visit duration

*Mean long shelter visit duration*: Mean duration per long shelter visit during the first 3 days

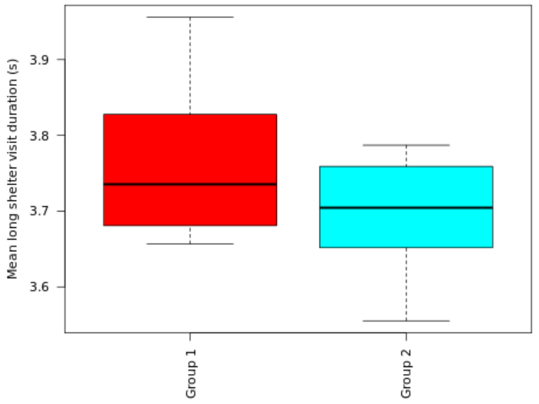

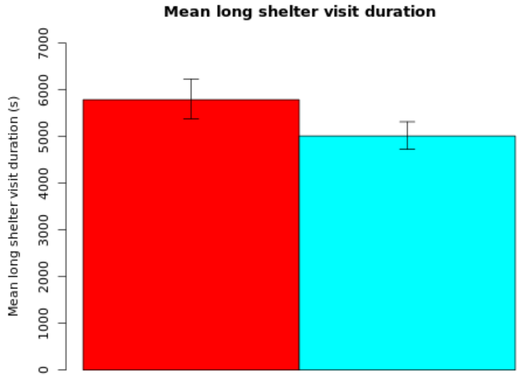

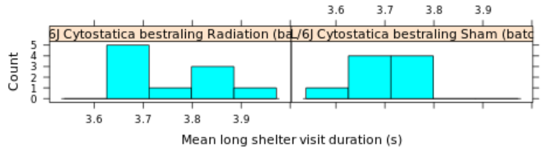

**NOTE: The statistics and boxplot are performed on log10 transformed data**

| Genotype | Mean | Median | SEM | N |
|---|---|---|---|---|
| (Gr 1): C57BL/6J Cytostatica bestraling Radiation (batch 22) | 5790.7478 | 0 | Up: 440.6876<br>Down: 409.5271 | 10 |
| (Gr 2): C57BL/6J Cytostatica bestraling Sham (batch 22) | 5012.934 | 0 | Up: 304.3587<br>Down: 286.9407 | 9 |

| | |
|---|---|
| **T-test:** | 1.5329 |
| **P-value:** | 0.1441 |
| **Df:** | 16.5927 |
| **Mann-Whitney U test:** | 0.2428 |

| Mice excluded by Quality Control |
|---|
| PH06469 (C57BL/6J - ) |

 © Sylics BV

Short shelter visit threshold

*Short shelter visit threshold*: Cut-off value to separate short and intermediate shelter visits

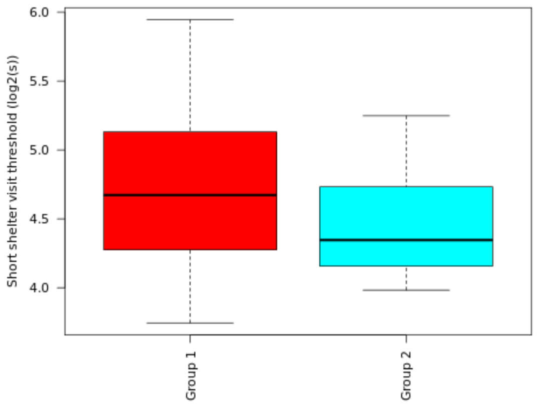

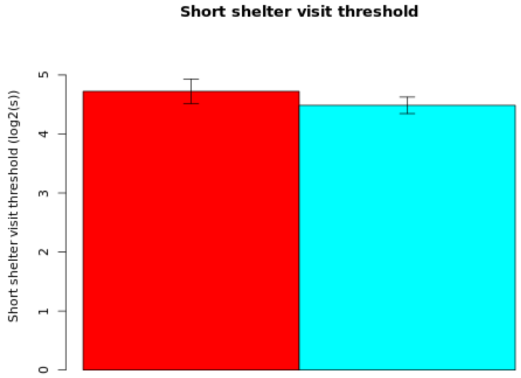

#### Secondary plot:

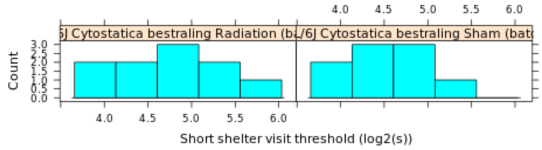

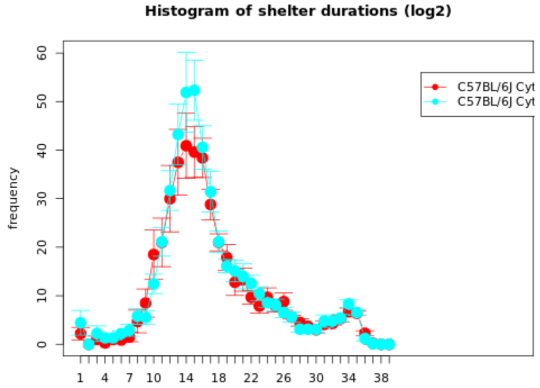

| Genotype | Mean | Median | SEM | N |
| --- | --- | --- | --- | --- |
| (Gr 1): C57BL/6J Cytostatica bestraling Radiation (batch 22) | 4.7226 | 4.6728 | 0.2077 | 10 |
| (Gr 2): C57BL/6J Cytostatica bestraling Sham (batch 22) | 4.4846 | 4.3461 | 0.1397 | 9 |

| | |
| --- | --- |
| **T-test:** | 0.9507 |
| **P-value:** | 0.3564 |
| **Df:** | 15.4301 |
| **Mann-Whitney U test:** | 0.4967 |

| Mice excluded by Quality Control |
| --- |
| PH06469 (C57BL/6J - ) |

 © Sylics BV

Long shelter visit fraction of total visits

*Long shelter visit fraction of total visits*: The fraction of shelter visits with duration longer than long shelter visit threshold

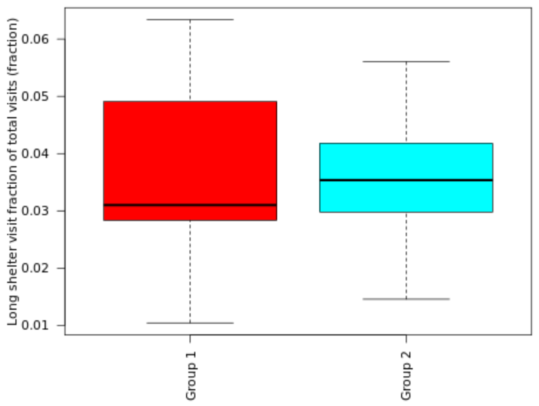

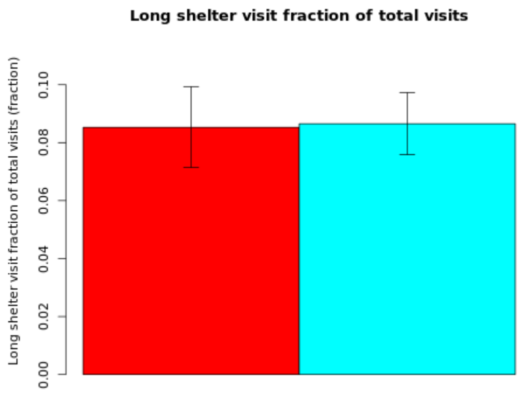

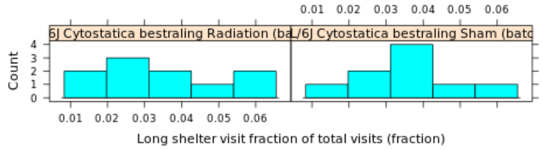

**Secondary plot:**

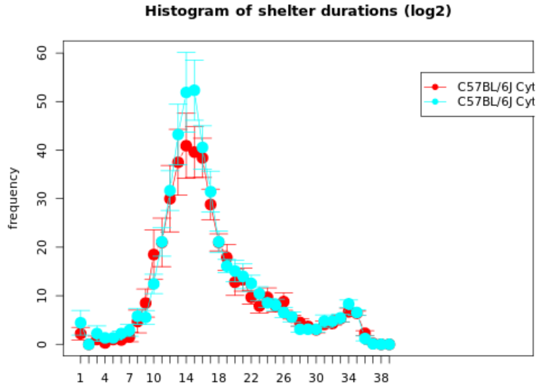

**NOTE: The statistics and boxplot are performed on log10 transformed data**

| Genotype | Mean | Median | SEM | N |
|---|---|---|---|---|
| (Gr 1): C57BL/6J Cytostatica bestraling Radiation (batch 22) | 0.0853 | 0 | Up: 0.014<br>Down: 0.0138 | 10 |
| (Gr 2): C57BL/6J Cytostatica bestraling Sham (batch 22) | 0.0865 | 0 | Up: 0.0107<br>Down: 0.0106 | 9 |

| | |
|---|---|
| **T-test:** | -0.0703 |
| **P-value:** | 0.9448 |
| **Df:** | 16.3443 |
| **Mann-Whitney U test:** | 0.6607 |

| Mice excluded by Quality Control |
|---|
| PH06469 (C57BL/6J - ) |

 © Sylics BV

Sylics

Long shelter visit threshold

*Long shelter visit threshold*: Cut-off value to separate intermediate and long shelter visits

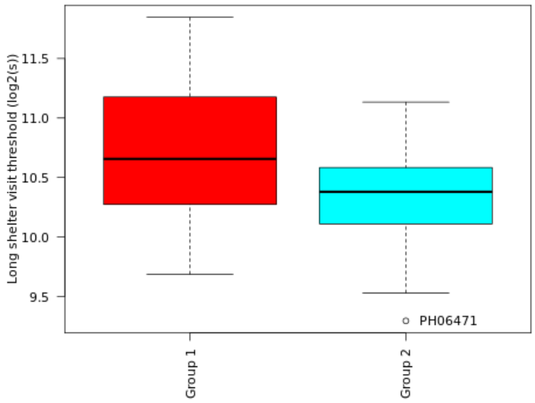

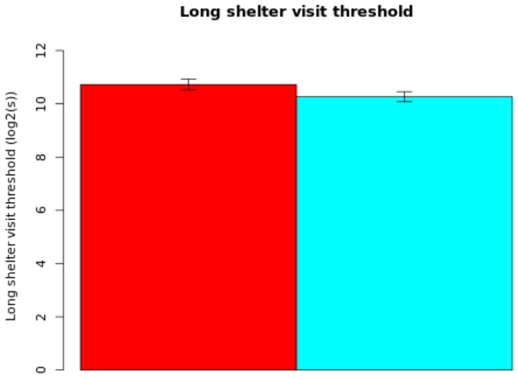

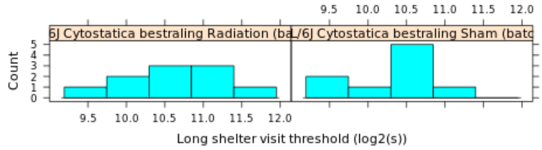

#### Secondary plot:

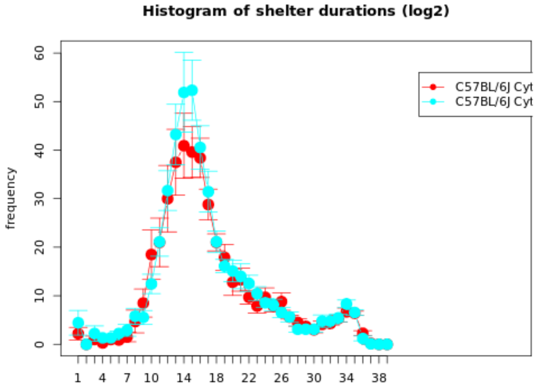

| Genotype | Mean | Median | SEM | N |
|---|---|---|---|---|
| (Gr 1): C57BL/6J Cytostatica bestraling Radiation (batch 22) | 10.7241 | 10.6562 | 0.206 | 10 |
| (Gr 2): C57BL/6J Cytostatica bestraling Sham (batch 22) | 10.2695 | 10.3799 | 0.1877 | 9 |

| | |
|---|---|
| **T-test:** | 1.6312 |
| **P-value:** | 0.1213 |
| **Df:** | 16.9802 |
| **Mann-Whitney U test:** | 0.1823 |

| Mice excluded by Quality Control |
|---|
| PH06469 (C57BL/6J - ) |

© Sylics BV

Short shelter visit duration - light

*Short shelter visit duration - light*: Cummulative duration of short shelter visits during the light phase

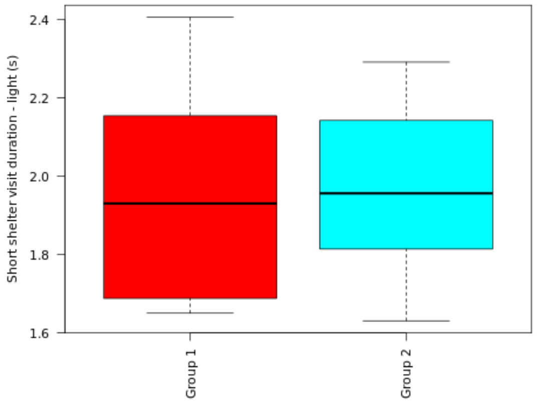

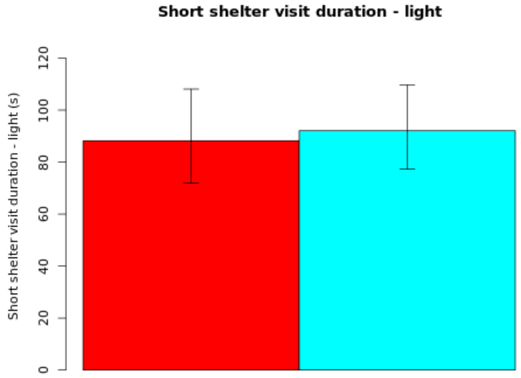

#### Secondary plot:

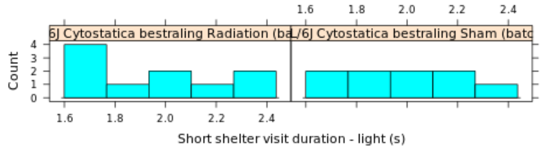

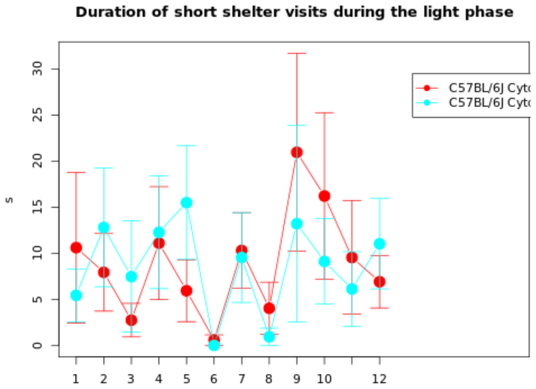

**NOTE: The statistics and boxplot are performed on log10 transformed data**

| Genotype | Mean | Median | SEM | N |
|---|---|---|---|---|
| (Gr 1): C57BL/6J Cytostatica bestraling Radiation (batch 22) | 88.2077 | 0 | Up: 19.862<br>Down: 16.245 | 10 |
| (Gr 2): C57BL/6J Cytostatica bestraling Sham (batch 22) | 92.0637 | 0 | Up: 17.5891<br>Down: 14.7932 | 9 |

| | |
|---|---|
| **T-test:** | -0.1595 |
| **P-value:** | 0.8752 |
| **Df:** | 16.8644 |
| **Mann-Whitney U test:** | 0.9048 |

| Mice excluded by Quality Control |
|---|
| PH06469 (C57BL/6J - ) |

 *© Sylics BV*

Mean short shelter visit duration - light

*Mean short shelter visit duration - light*: Mean duration per short shelter visit during the light phase

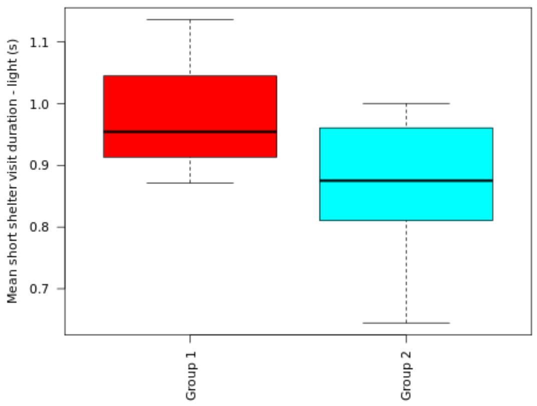

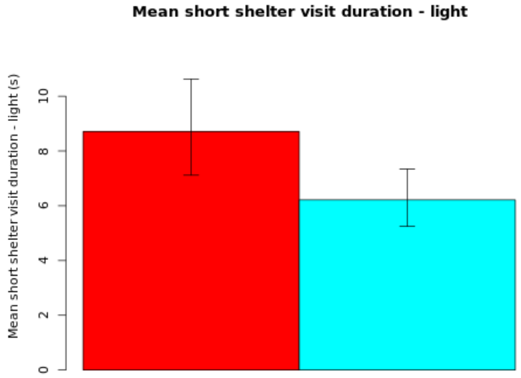

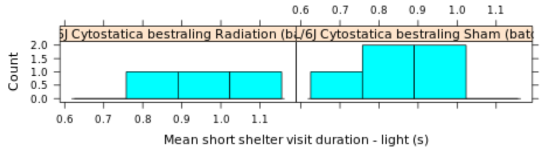

**NOTE: The statistics and boxplot are performed on log10 transformed data**

| Genotype | Mean | Median | SEM | N |
|---|---|---|---|---|
| (Gr 1): C57BL/6J Cytostatica bestraling Radiation (batch 22) | 8.7132 | 0 | Up: 1.913<br>Down: 1.5983 | 3 |
| (Gr 2): C57BL/6J Cytostatica bestraling Sham (batch 22) | 6.2168 | 0 | Up: 1.1234<br>Down: 0.9721 | 5 |

| | |
|---|---|
| **T-test:** | 1.2874 |
| **P-value:** | 0.2603 |
| **Df:** | 4.4881 |
| **Mann-Whitney U test:** | 0.5714 |

| Mice excluded by Quality Control |
|---|
| PH06469 (C57BL/6J - ) |

 © Sylics BV

Short shelter visit number - light

*Short shelter visit number - light*: Cummulative number of short shelter visits during the light phase

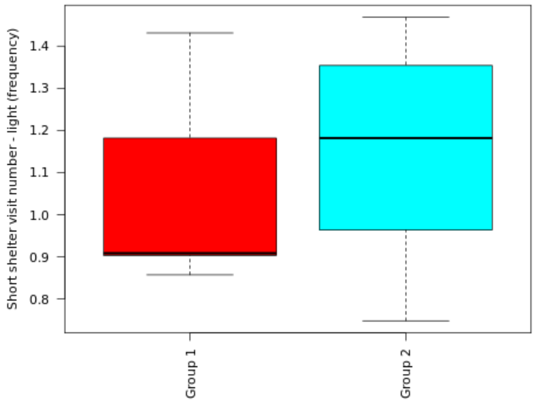

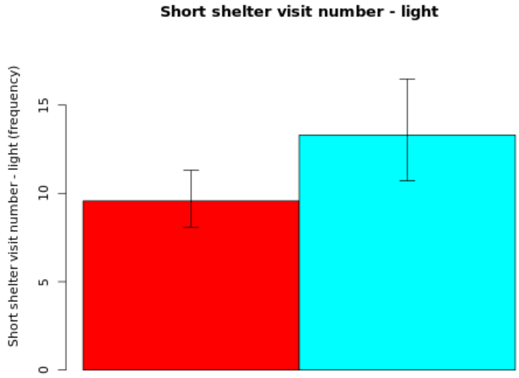

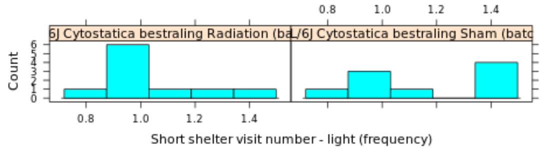

**Secondary plot:**

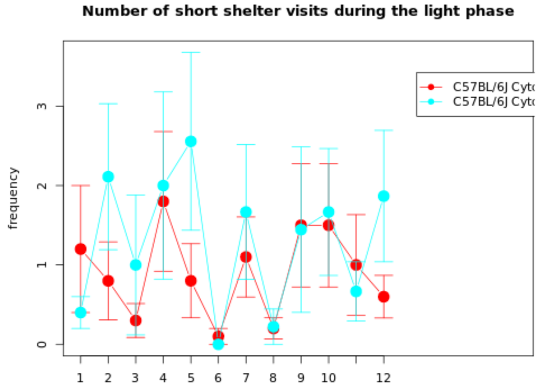

**NOTE: The statistics and boxplot are performed on log10 transformed data**

| Genotype | Mean | Median | SEM | N |
|---|---|---|---|---|
| (Gr 1): C57BL/6J Cytostatica bestraling Radiation (batch 22) | 9.576 | 0 | Up: 1.7265<br>Down: 1.4842 | 10 |
| (Gr 2): C57BL/6J Cytostatica bestraling Sham (batch 22) | 13.3083 | 0 | Up: 3.1668<br>Down: 2.5929 | 9 |

| | |
|---|---|
| **T-test:** | -1.2057 |
| **P-value:** | 0.2462 |
| **Df:** | 15.3152 |
| **Mann-Whitney U test:** | 0.1612 |

| Mice excluded by Quality Control |
|---|
| PH06469 (C57BL/6J - ) |

 *© Sylics BV*

Long shelter visit duration - light

*Long shelter visit duration - light*: Cummulative duration of long shelter visits during the light phase

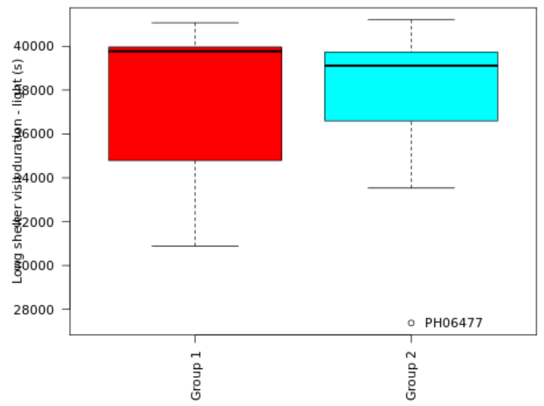

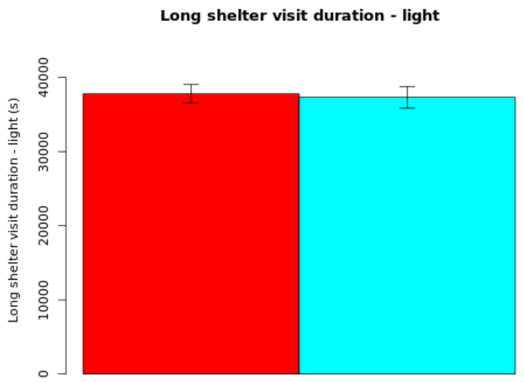

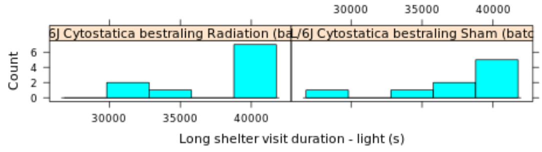

**Secondary plot:**

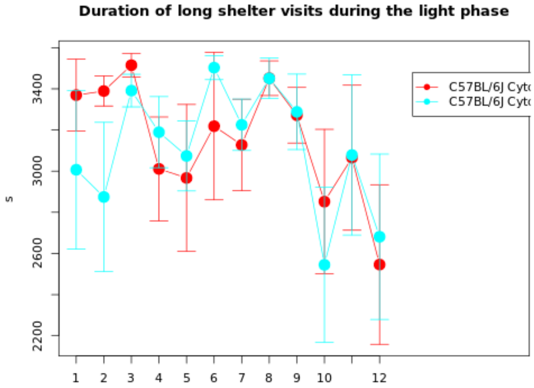

| Genotype | Mean | Median | SEM | N |
|---|---|---|---|---|
| (Gr 1): C57BL/6J Cytostatica bestraling Radiation (batch 22) | 37785.1267 | 39773.82 | 1245.3644 | 10 |
| (Gr 2): C57BL/6J Cytostatica bestraling Sham (batch 22) | 37308.1837 | 39117.0933 | 1452.8056 | 9 |

| | |
|---|---|
| **T-test:** | 0.2492 |
| **P-value:** | 0.8063 |
| **Df:** | 16.2685 |
| **Mann-Whitney U test:** | 0.6038 |

| Mice excluded by Quality Control |
|---|
| PH06469 (C57BL/6J - ) |

 © Sylics BV

Long shelter visit number - light

*Long shelter visit number - light*: Cummulative number of long shelter visits during the light phase

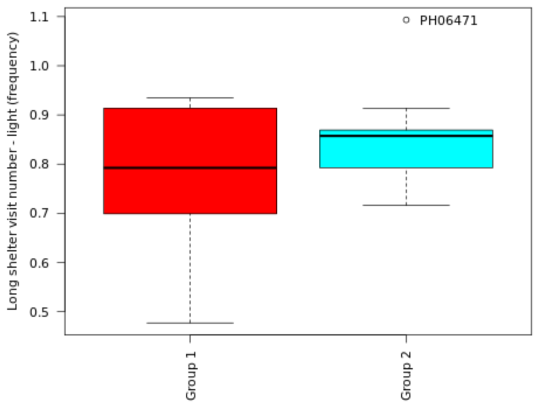

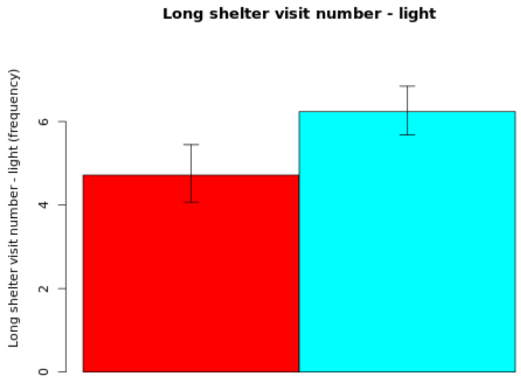

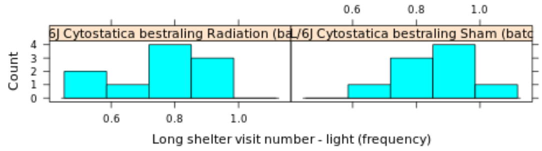

**Secondary plot:**

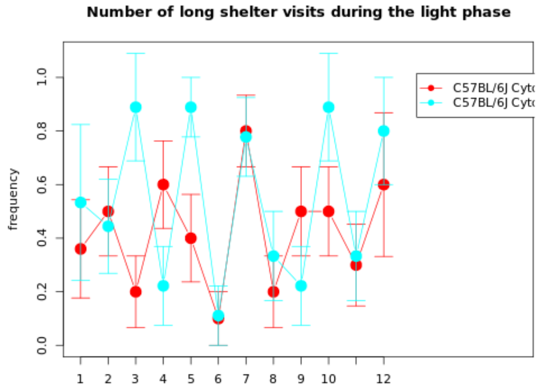

**NOTE: The statistics and boxplot are performed on log10 transformed data**

| Genotype | Mean | Median | SEM | N |
|---|---|---|---|---|
| (Gr 1): C57BL/6J Cytostatica bestraling Radiation (batch 22) | 4.7157 | 0 | Up: 0.7295<br>Down: 0.6469 | 10 |
| (Gr 2): C57BL/6J Cytostatica bestraling Sham (batch 22) | 6.2388 | 0 | Up: 0.6053<br>Down: 0.5586 | 9 |

| | |
|---|---|
| **T-test:** | -1.6351 |
| **P-value:** | 0.1223 |
| **Df:** | 15.3855 |
| **Mann-Whitney U test:** | 0.2311 |

| Mice excluded by Quality Control |
|---|
| PH06469 (C57BL/6J - ) |

 *© Sylics BV*

### Spontaneous behavior (Group 3: habituation)

With respect to spontaneous behaviors in the first three days in the PhenoTyper, 6 groups of behavioral parameters are defined as described below. The first two groups describe specific behavioral elements related to kinematics of mice (description of movement characteristics, group1) and sheltering behavior (group 2). These behavioral parameters were analyzed with respect to temporal aspects, in particular over 4 different time scales, i.e., habituation effects across multiple days (group 3), effects of DarkLight phase across 24h (group 4), differences in the pattern of behavior in the few hours before and after phase shifts (group5), and differences in activity bout properties on the sub-minute time scale (group 6).

**Group 3: habituation**
In general, mice are more active during the first hours of the dark phase on day 1, than during the same period on day 3. To capture these habituation effects, we analyzed the change in activity from day 1 to day 3, i.e., the habituation phase, by taking the ratio of day 3 over day 1.

 *© Sylics BV*

Activity duration - habituation ratio dark

*Activity duration - habituation ratio dark*: Habituation effect: Change in cumulative activity duration during the dark phase of day 3 compared to the dark phase of day 1

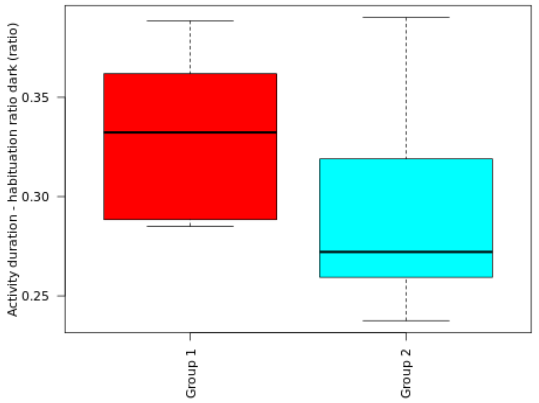

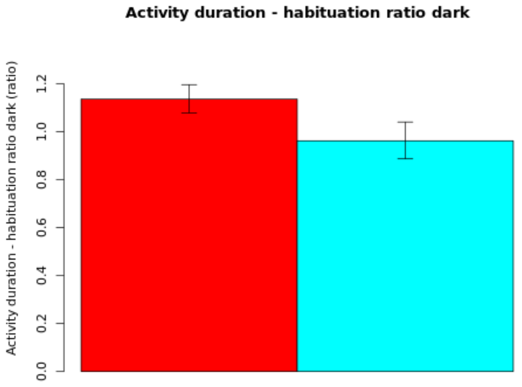

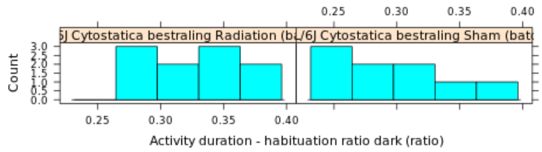

**NOTE: The statistics and boxplot are performed on log10 transformed data**

| Genotype | Mean | Median | SEM | N |
|---|---|---|---|---|
| (Gr 1): C57BL/6J Cytostatica bestraling Radiation (batch 22) | 1.1365 | 0 | Up: 0.0597<br>Down: 0.0581 | 10 |
| (Gr 2): C57BL/6J Cytostatica bestraling Sham (batch 22) | 0.9623 | 0 | Up: 0.0783<br>Down: 0.0753 | 9 |

| | |
|---|---|
| **T-test:** | 1.7778 |
| **P-value:** | 0.0961 |
| **Df:** | 14.6862 |
| **Mann-Whitney U test:** | 0.0947 |

| Mice excluded by Quality Control |
|---|
| PH06469 (C57BL/6J - ) |

 © Sylics BV

Mean activity duration - habituation ratio dark

*Mean activity duration - habituation ratio dark*: Habituation effect: Change in mean activity duration during the dark phase of day 3 compared to the dark phase of day 1

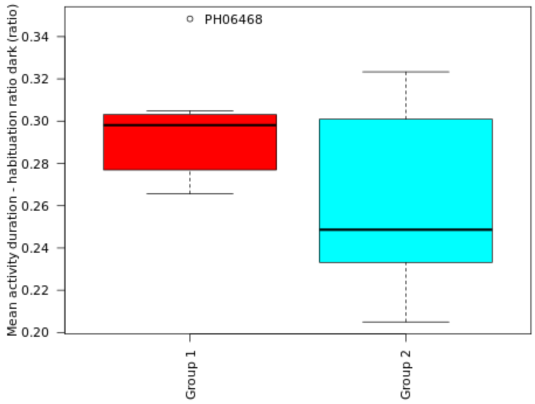

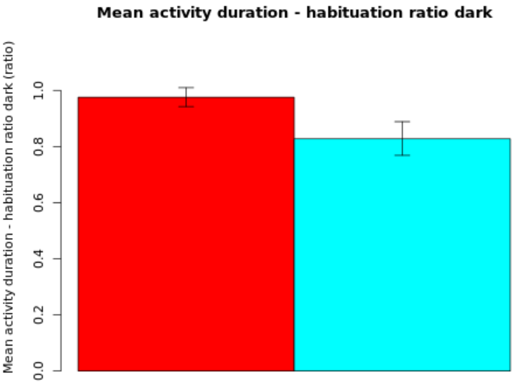

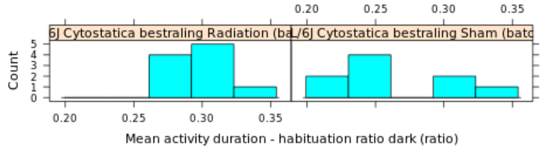

**NOTE: The statistics and boxplot are performed on log10 transformed data**

| Genotype | Mean | Median | SEM | N |
|---|---|---|---|---|
| (Gr 1): C57BL/6J Cytostatica bestraling Radiation (batch 22) | 0.9757 | 0 | Up: 0.034<br>Down: 0.0334 | 10 |
| (Gr 2): C57BL/6J Cytostatica bestraling Sham (batch 22) | 0.8276 | 0 | Up: 0.0611<br>Down: 0.0591 | 9 |

| | |
|---|---|
| **T-test:** | 2.1037 |
| **P-value:** | 0.057 |
| **Df:** | 12.1005 |
| **Mann-Whitney U test:** | 0.0789 |

| Mice excluded by Quality Control |
|---|
| PH06469 (C57BL/6J - ) |

 © Sylics BV

Activity number - habituation ratio dark

*Activity number - habituation ratio dark*: Habituation effect: Change in cummulative activity number during the dark phase of day 3 compared to the dark phase of day 1

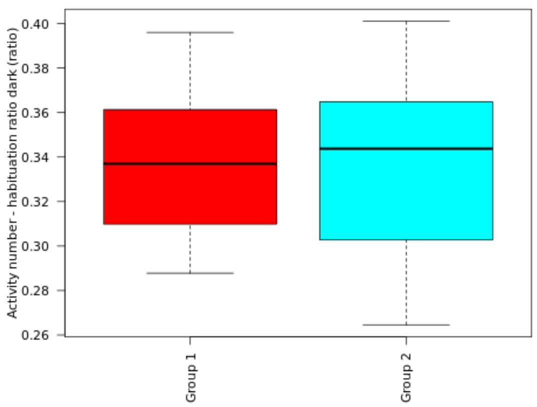

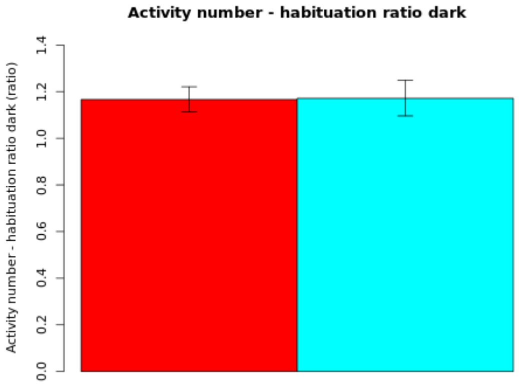

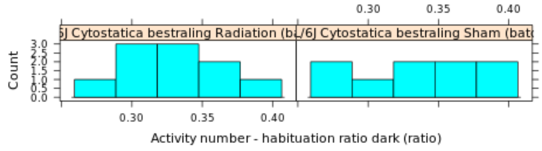

**NOTE: The statistics and boxplot are performed on log10 transformed data**

| Genotype | Mean | Median | SEM | N |
|---|---|---|---|---|
| (Gr 1): C57BL/6J Cytostatica bestraling Radiation (batch 22) | 1.1664 | 0 | Up: 0.0544<br>Down: 0.053 | 10 |
| (Gr 2): C57BL/6J Cytostatica bestraling Sham (batch 22) | 1.1712 | 0 | Up: 0.0779<br>Down: 0.0752 | 9 |

| | |
|---|---|
| **T-test:** | -0.0514 |
| **P-value:** | 0.9597 |
| **Df:** | 14.6788 |
| **Mann-Whitney U test:** | 0.9682 |

| Mice excluded by Quality Control |
|---|
| PH06469 (C57BL/6J - ) |

 © Sylics BV

Mean short arrest duration - habituation ratio dark

*Mean short arrest duration - habituation ratio dark*: Habituation effect: Change in mean short arrest duration during the dark phase of day 3 compared to the dark phase of day 1

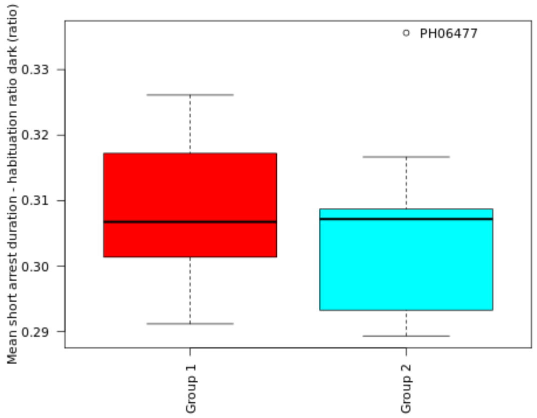

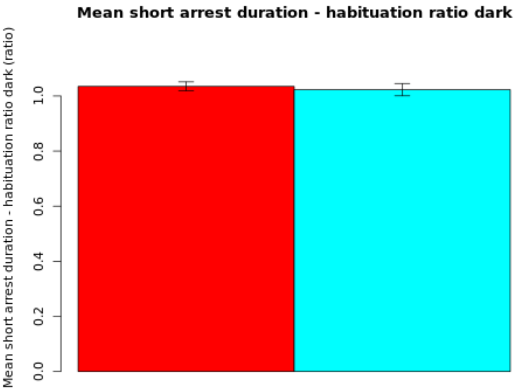

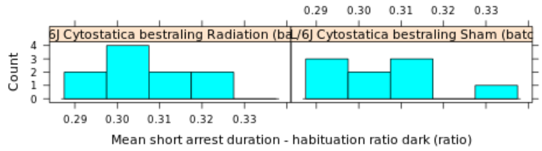

**NOTE: The statistics and boxplot are performed on log10 transformed data**

| Genotype | Mean | Median | SEM | N |
|---|---|---|---|---|
| (Gr 1): C57BL/6J Cytostatica bestraling Radiation (batch 22) | 1.0348 | 0 | Up: 0.0164<br>Down: 0.0162 | 10 |
| (Gr 2): C57BL/6J Cytostatica bestraling Sham (batch 22) | 1.0227 | 0 | Up: 0.0225<br>Down: 0.0222 | 9 |

| | |
|---|---|
| **T-test:** | 0.4353 |
| **P-value:** | 0.6696 |
| **Df:** | 14.9362 |
| **Mann-Whitney U test:** | 0.6607 |

| Mice excluded by Quality Control |
|---|
| PH06469 (C57BL/6J - ) |

 © Sylics BV

Long arrest duration - habituation ratio dark

*Long arrest duration - habituation ratio dark*: Habituation effect: Change in cummulative long arrest long arrest duration during the dark phase of day 3 compared to the dark phase of day 1

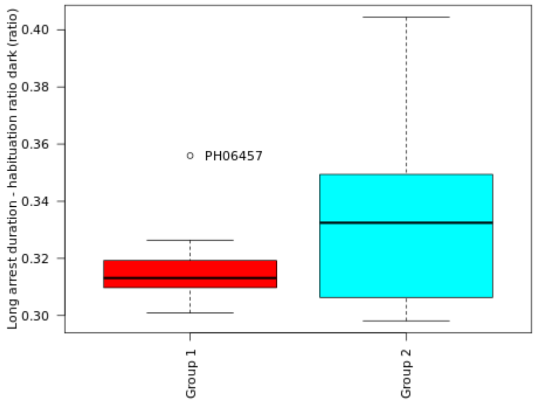

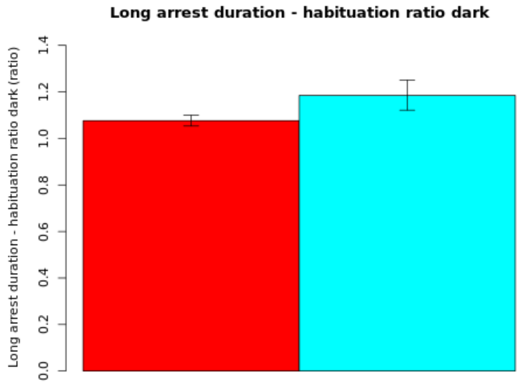

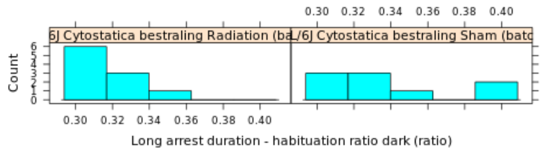

**NOTE: The statistics and boxplot are performed on log10 transformed data**

| Genotype | Mean | Median | SEM | N |
|---|---|---|---|---|
| (Gr 1): C57BL/6J Cytostatica bestraling Radiation (batch 22) | 1.0763 | 0 | Up: 0.0233<br>Down: 0.0231 | 10 |
| (Gr 2): C57BL/6J Cytostatica bestraling Sham (batch 22) | 1.1852 | 0 | Up: 0.0656<br>Down: 0.0637 | 9 |

| | |
|---|---|
| **T-test:** | -1.6164 |
| **P-value:** | 0.1363 |
| **Df:** | 10.2623 |
| **Mann-Whitney U test:** | 0.3154 |

| Mice excluded by Quality Control |
|---|
| PH06469 (C57BL/6J - ) |

Mean long arrest duration - habituation ratio dark

*Mean long arrest duration - habituation ratio dark*: Habituation effect: Change in mean long arrest duration during the dark phase of day 3 compared to the dark phase of day 1

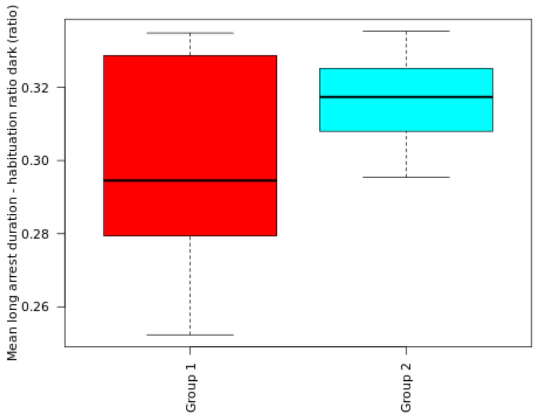

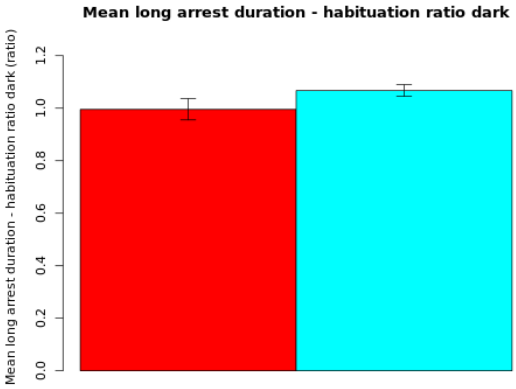

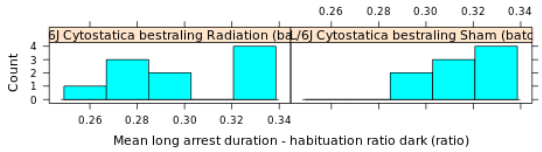

**NOTE: The statistics and boxplot are performed on log10 transformed data**

| Genotype | Mean | Median | SEM | N |
|---|---|---|---|---|
| (Gr 1): C57BL/6J Cytostatica bestraling Radiation (batch 22) | 0.9954 | 0 | Up: 0.0408<br>Down: 0.04 | 10 |
| (Gr 2): C57BL/6J Cytostatica bestraling Sham (batch 22) | 1.0673 | 0 | Up: 0.0222<br>Down: 0.0219 | 9 |

| | |
|---|---|
| **T-test:** | -1.5454 |
| **P-value:** | 0.1453 |
| **Df:** | 13.5085 |
| **Mann-Whitney U test:** | 0.2428 |

| Mice excluded by Quality Control |
|---|
| PH06469 (C57BL/6J - ) |

 © Sylics BV

Sylics

Long arrest number - habituation ratio dark

*Long arrest number - habituation ratio dark*: Habituation effect: Change in cummulative long arrest long arrest number during the dark phase of day 3 compared to the dark phase of day 1

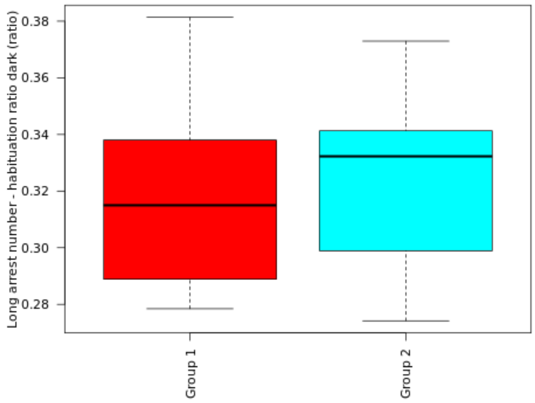

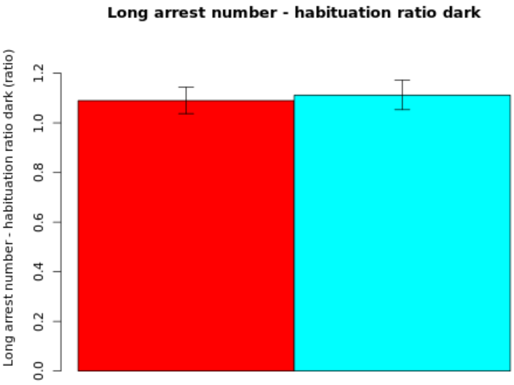

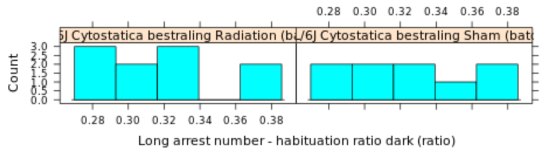

**NOTE: The statistics and boxplot are performed on log10 transformed data**

| Genotype | Mean | Median | SEM | N |
|---|---|---|---|---|
| (Gr 1): C57BL/6J Cytostatica bestraling Radiation (batch 22) | 1.0898 | 0 | Up: 0.0539<br>Down: 0.0525 | 10 |
| (Gr 2): C57BL/6J Cytostatica bestraling Sham (batch 22) | 1.1118 | 0 | Up: 0.0606<br>Down: 0.0589 | 9 |

| | |
|---|---|
| **T-test:** | -0.2761 |
| **P-value:** | 0.7859 |
| **Df:** | 16.5559 |
| **Mann-Whitney U test:** | 0.7802 |

| Mice excluded by Quality Control |
|---|
| PH06469 (C57BL/6J - ) |

 © Sylics BV

Feeding zone duration - habituation ratio dark

*Feeding zone duration - habituation ratio dark*: Habituation effect: Change in cummulative Feeding zone duration during the dark phase of day 3 compared to the dark phase of day 1

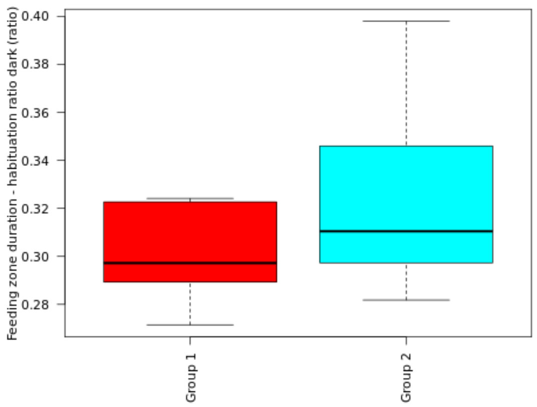

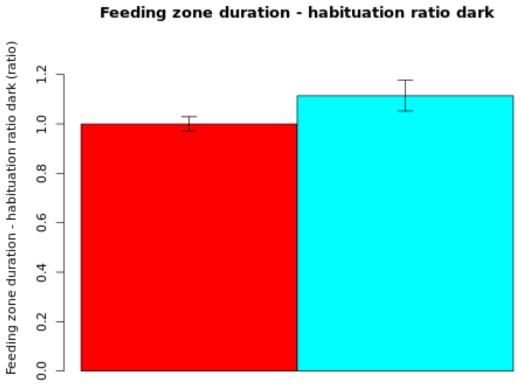

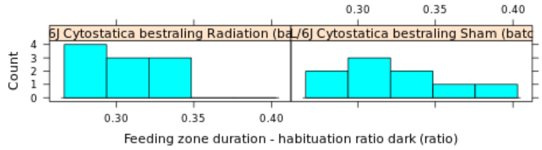

**NOTE: The statistics and boxplot are performed on log10 transformed data**

| Genotype | Mean | Median | SEM | N |
|---|---|---|---|---|
| (Gr 1): C57BL/6J Cytostatica bestraling Radiation (batch 22) | 0.9999 | 0 | Up: 0.0294<br>Down: 0.0289 | 10 |
| (Gr 2): C57BL/6J Cytostatica bestraling Sham (batch 22) | 1.1141 | 0 | Up: 0.0635<br>Down: 0.0617 | 9 |

| | |
|---|---|
| **T-test:** | -1.6833 |
| **P-value:** | 0.1187 |
| **Df:** | 11.734 |
| **Mann-Whitney U test:** | 0.1823 |

| Mice excluded by Quality Control |
|---|
| PH06469 (C57BL/6J - ) |

Mean short shelter visit duration - habituation ratio dark

*Mean short shelter visit duration - habituation ratio dark*: Habituation effect: Change in mean short shelter visit duration during the dark phase of day 3 compared to the dark phase of day 1

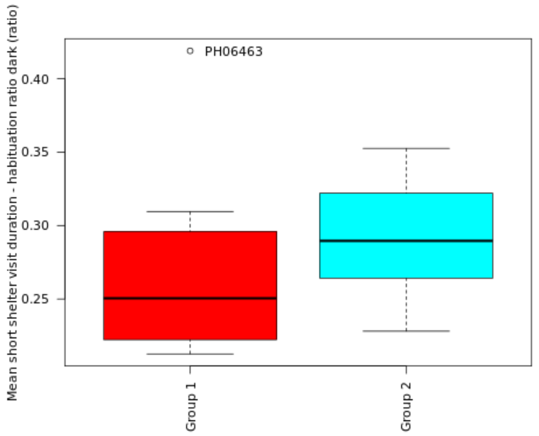

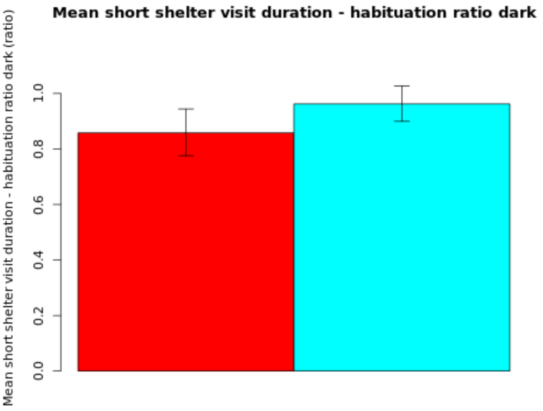

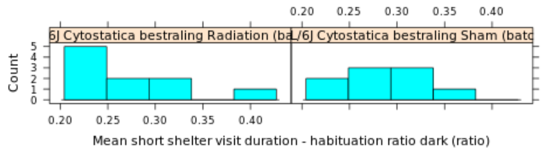

**NOTE: The statistics and boxplot are performed on log10 transformed data**

| Genotype | Mean | Median | SEM | N |
|---|---|---|---|---|
| (Gr 1): C57BL/6J Cytostatica bestraling Radiation (batch 22) | 0.8582 | 0 | Up: 0.086<br>Down: 0.0822 | 10 |
| (Gr 2): C57BL/6J Cytostatica bestraling Sham (batch 22) | 0.9622 | 0 | Up: 0.0649<br>Down: 0.0628 | 9 |

| | |
|---|---|
| **T-test:** | -0.9773 |
| **P-value:** | 0.343 |
| **Df:** | 15.9266 |
| **Mann-Whitney U test:** | 0.1333 |

| Mice excluded by Quality Control |
|---|
| PH06469 (C57BL/6J - ) |

© Sylics BV

Long shelter visit duration - habituation ratio dark

*Long shelter visit duration - habituation ratio dark*: Habituation effect: Change in cummulative long shelter visit duration during the dark phase of day 3 compared to the dark phase of day 1

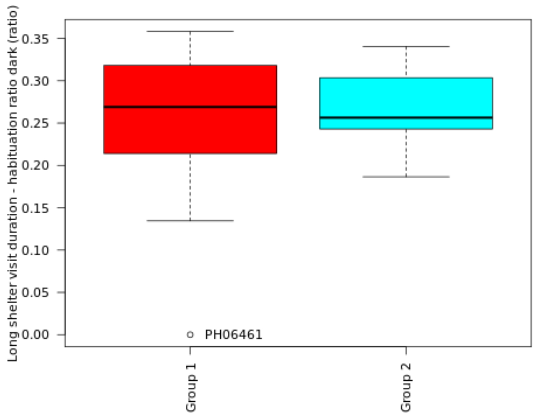

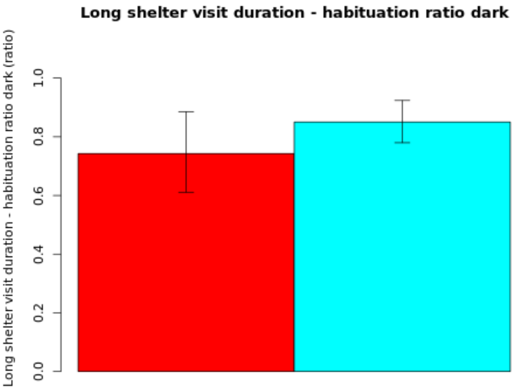

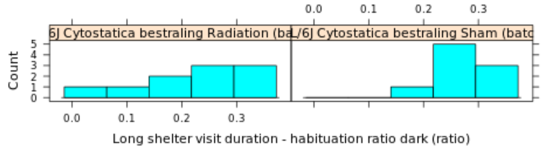

**NOTE: The statistics and boxplot are performed on log10 transformed data**

| Genotype | Mean | Median | SEM | N |
|---|---|---|---|---|
| (Gr 1): C57BL/6J Cytostatica bestraling Radiation (batch 22) | 0.743 | 0 | Up: 0.1426<br>Down: 0.1318 | 10 |
| (Gr 2): C57BL/6J Cytostatica bestraling Sham (batch 22) | 0.8504 | 0 | Up: 0.0733<br>Down: 0.0705 | 9 |

| | |
|---|---|
| **T-test:** | -0.6819 |
| **P-value:** | 0.5072 |
| **Df:** | 13.0554 |
| **Mann-Whitney U test:** | 0.8421 |

| Mice excluded by Quality Control |
|---|
| PH06469 (C57BL/6J - ) |

© Sylics BV

Mean long movement distance - habituation ratio dark

*Mean long movement distance - habituation ratio dark*: Habituation effect: Change in mean long movement distance during the dark phase of day 3 compared to the dark phase of day 1

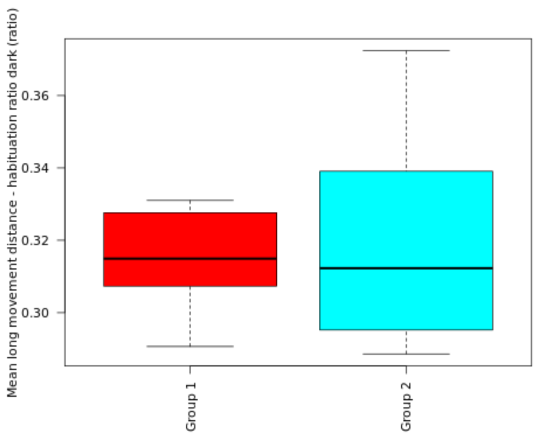

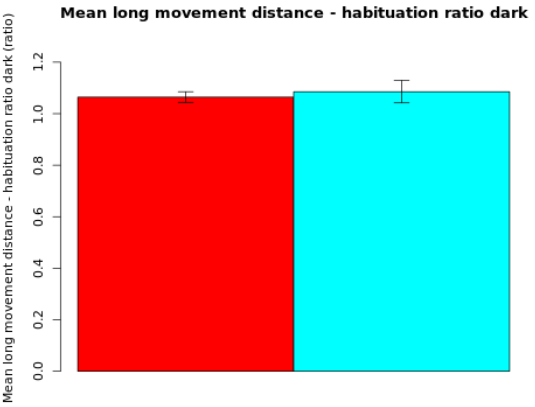

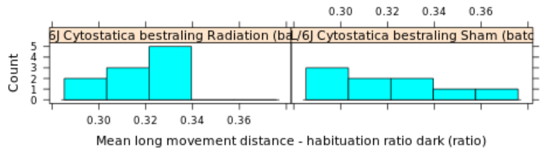

**NOTE: The statistics and boxplot are performed on log10 transformed data**

| Genotype | Mean | Median | SEM | N |
|---|---|---|---|---|
| (Gr 1): C57BL/6J Cytostatica bestraling Radiation (batch 22) | 1.0645 | 0 | Up: 0.0207<br>Down: 0.0205 | 10 |
| (Gr 2): C57BL/6J Cytostatica bestraling Sham (batch 22) | 1.0853 | 0 | Up: 0.0442<br>Down: 0.0433 | 9 |

| | |
|---|---|
| **T-test:** | -0.433 |
| **P-value:** | 0.673 |
| **Df:** | 11.4949 |
| **Mann-Whitney U test:** | 1 |

| Mice excluded by Quality Control |
|---|
| PH06469 (C57BL/6J - ) |

 © Sylics BV

Mean short movement distance - habituation ratio dark

*Mean short movement distance - habituation ratio dark*: Habituation effect: Change in mean short movement distance during the dark phase of day 3 compared to the dark phase of day 1

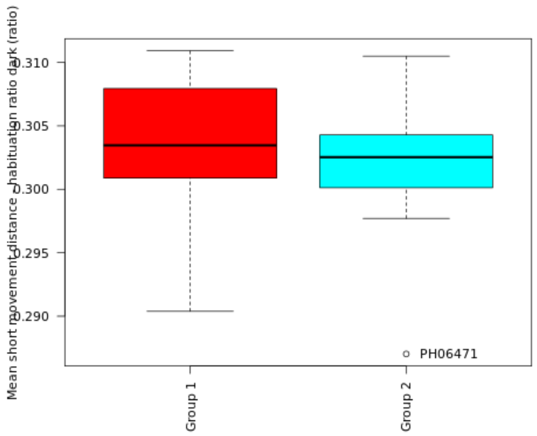

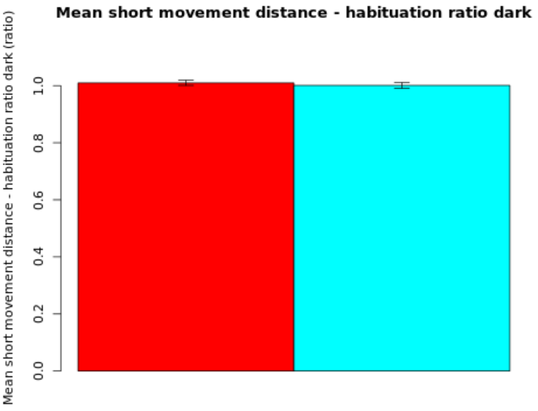

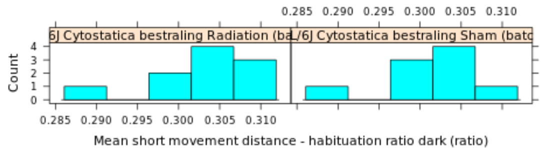

**NOTE: The statistics and boxplot are performed on log10 transformed data**

| Genotype | Mean | Median | SEM | N |
|---|---|---|---|---|
| (Gr 1): C57BL/6J Cytostatica bestraling Radiation (batch 22) | 1.0095 | 0 | Up: 0.0088<br>Down: 0.0088 | 10 |
| (Gr 2): C57BL/6J Cytostatica bestraling Sham (batch 22) | 1.0007 | 0 | Up: 0.0099<br>Down: 0.0099 | 9 |

| | |
|---|---|
| **T-test:** | 0.6664 |
| **P-value:** | 0.5144 |
| **Df:** | 16.4715 |
| **Mann-Whitney U test:** | 0.4967 |

| Mice excluded by Quality Control |
|---|
| PH06469 (C57BL/6J - ) |

Sylics

OnShelter zone duration - habituation ratio dark

*OnShelter zone duration - habituation ratio dark*: Habituation effect: Change in cummulative OnShelter zone duration during the dark phase of day 3 compared to the dark phase of day 1

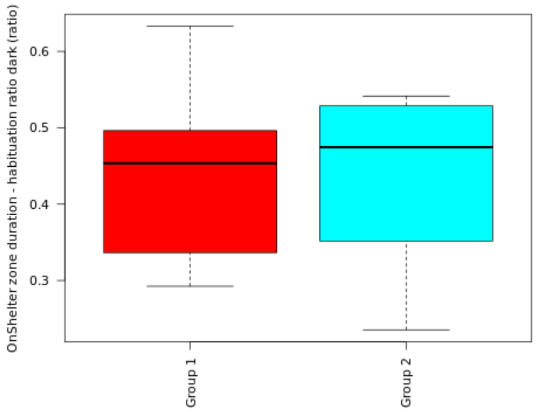

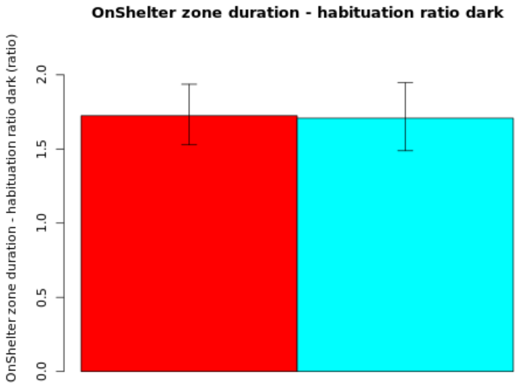

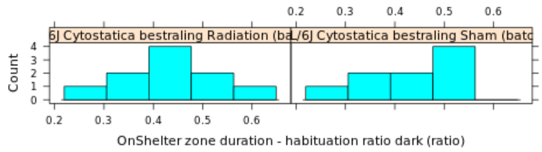

**NOTE: The statistics and boxplot are performed on log10 transformed data**

| Genotype | Mean | Median | SEM | N |
|---|---|---|---|---|
| (Gr 1): C57BL/6J Cytostatica bestraling Radiation (batch 22) | 1.7246 | 0 | Up: 0.2117<br>Down: 0.1964 | 10 |
| (Gr 2): C57BL/6J Cytostatica bestraling Sham (batch 22) | 1.7079 | 0 | Up: 0.2393<br>Down: 0.2199 | 9 |

| | |
|---|---|
| **T-test:** | 0.0544 |
| **P-value:** | 0.9573 |
| **Df:** | 16.453 |
| **Mann-Whitney U test:** | 0.6607 |

| Mice excluded by Quality Control |
|---|
| PH06469 (C57BL/6J - ) |

 *© Sylics BV*

Spout zone duration - habituation ratio dark

*Spout zone duration - habituation ratio dark*: Habituation effect: Change in cummulative Spout zone duration during the dark phase of day 3 compared to the dark phase of day 1

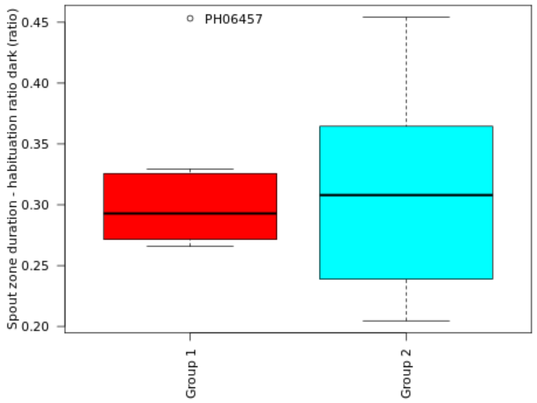

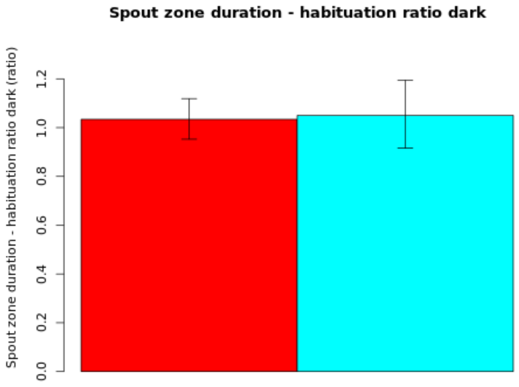

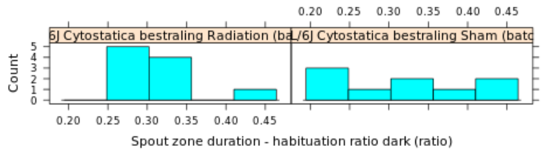

**NOTE: The statistics and boxplot are performed on log10 transformed data**

| Genotype | Mean | Median | SEM | N |
|---|---|---|---|---|
| (Gr 1): C57BL/6J Cytostatica bestraling Radiation (batch 22) | 1.0338 | 0 | Up: 0.0849<br>Down: 0.0815 | 10 |
| (Gr 2): C57BL/6J Cytostatica bestraling Sham (batch 22) | 1.0502 | 0 | Up: 0.1442<br>Down: 0.1347 | 9 |

| | |
|---|---|
| **T-test:** | -0.1009 |
| **P-value:** | 0.9211 |
| **Df:** | 13.2905 |
| **Mann-Whitney U test:** | 0.9682 |

| Mice excluded by Quality Control |
|---|
| PH06469 (C57BL/6J - ) |

 © Sylics BV

Activity duration - habituation ratio light

*Activity duration - habituation ratio light*: Habituation effect: Change in cummulative activity duration during the light phase of day 3 compared to the light phase of day 1

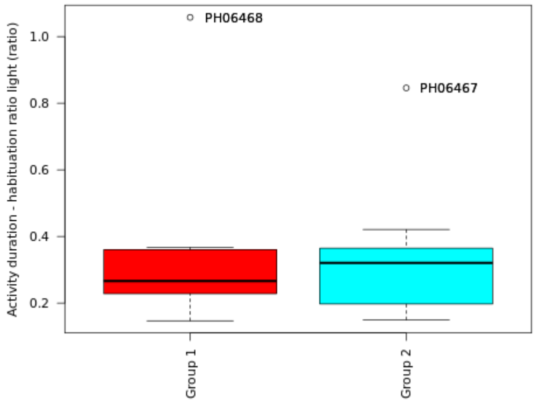

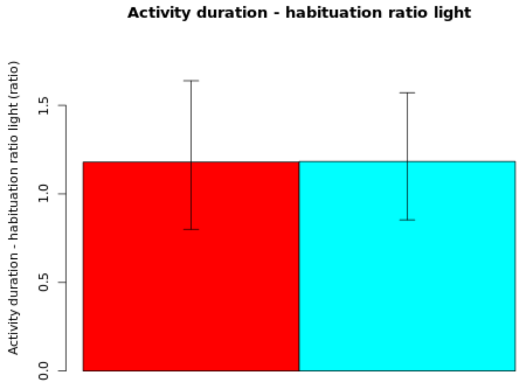

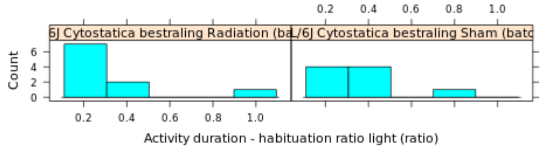

**NOTE: The statistics and boxplot are performed on log10 transformed data**

| Genotype | Mean | Median | SEM | N |
|---|---|---|---|---|
| (Gr 1): C57BL/6J Cytostatica bestraling Radiation (batch 22) | 1.1796 | 0 | Up: 0.4595<br>Down: 0.3795 | 10 |
| (Gr 2): C57BL/6J Cytostatica bestraling Sham (batch 22) | 1.1833 | 0 | Up: 0.3882<br>Down: 0.3296 | 9 |

| | |
|---|---|
| **T-test:** | -0.0068 |
| **P-value:** | 0.9947 |
| **Df:** | 16.8442 |
| **Mann-Whitney U test:** | 0.8421 |

| Mice excluded by Quality Control |
|---|
| PH06469 (C57BL/6J - ) |

 © Sylics BV

Mean activity duration - habituation ratio light

*Mean activity duration - habituation ratio light*: Habituation effect: Change in mean activity duration during the light phase of day 3 compared to the light phase of day 1

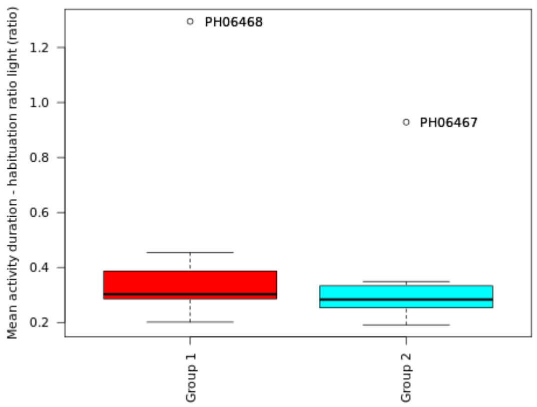

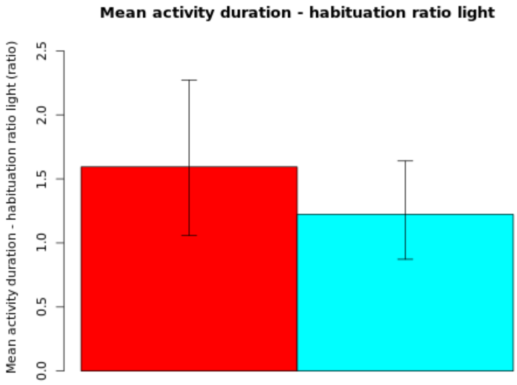

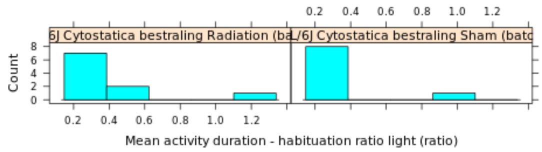

**NOTE: The statistics and boxplot are performed on log10 transformed data**

| Genotype | Mean | Median | SEM | N |
|---|---|---|---|---|
| (Gr 1): C57BL/6J Cytostatica bestraling Radiation (batch 22) | 1.5941 | 0 | Up: 0.6774<br>Down: 0.5372 | 10 |
| (Gr 2): C57BL/6J Cytostatica bestraling Sham (batch 22) | 1.2231 | 0 | Up: 0.4165<br>Down: 0.3508 | 9 |

| | |
|---|---|
| **T-test:** | 0.5347 |
| **P-value:** | 0.6002 |
| **Df:** | 16.1198 |
| **Mann-Whitney U test:** | 0.3154 |

| Mice excluded by Quality Control |
|---|
| PH06469 (C57BL/6J - ) |

Activity number - habituation ratio light

*Activity number - habituation ratio light*: Habituation effect: Change in cummulative activity number during the light phase of day 3 compared to the light phase of day 1

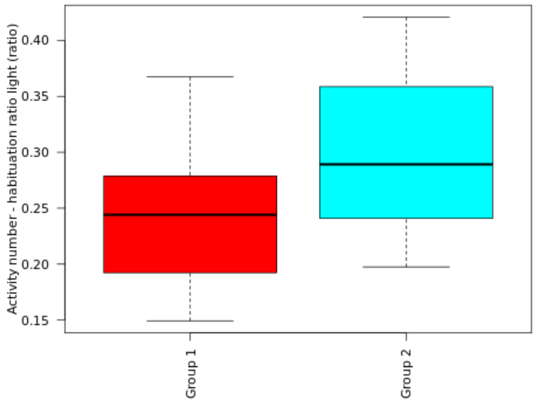

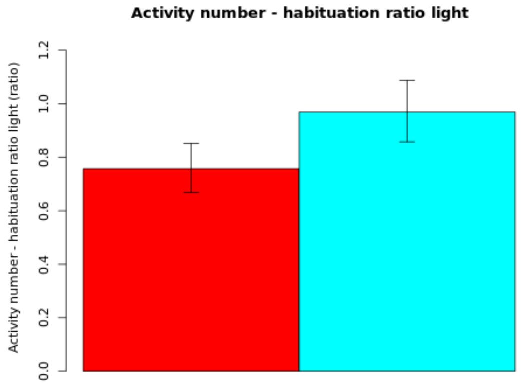

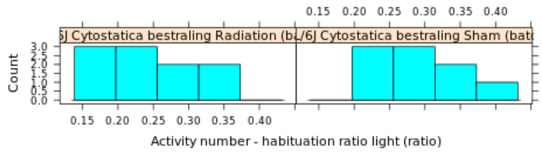

**NOTE: The statistics and boxplot are performed on log10 transformed data**

| Genotype | Mean | Median | SEM | N |
|---|---|---|---|---|
| (Gr 1): C57BL/6J Cytostatica bestraling Radiation (batch 22) | 0.757 | 0 | Up: 0.0944<br>Down: 0.0896 | 10 |
| (Gr 2): C57BL/6J Cytostatica bestraling Sham (batch 22) | 0.969 | 0 | Up: 0.1183<br>Down: 0.1116 | 9 |

| | |
|---|---|
| **T-test:** | -1.4536 |
| **P-value:** | 0.1648 |
| **Df:** | 16.5344 |
| **Mann-Whitney U test:** | 0.2428 |

| Mice excluded by Quality Control |
|---|
| PH06469 (C57BL/6J - ) |

 © Sylics BV

<u>Mean short arrest duration - habituation ratio light</u>

*Mean short arrest duration - habituation ratio light*: Habituation effect: Change in mean short arrest duration during the light phase of day 3 compared to the light phase of day 1

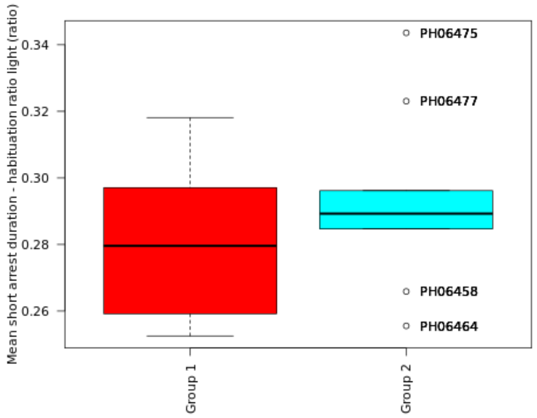

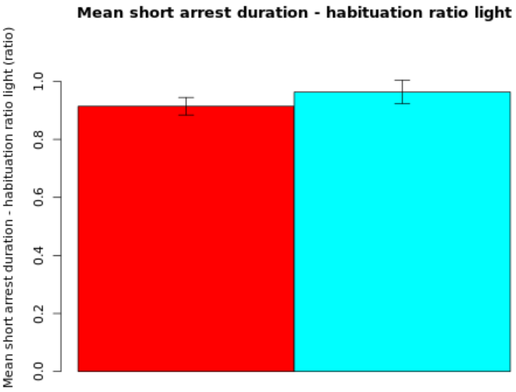

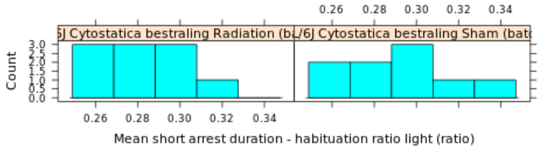

**NOTE: The statistics and boxplot are performed on log10 transformed data**

| Genotype | Mean | Median | SEM | N |
|---|---|---|---|---|
| (Gr 1): C57BL/6J Cytostatica bestraling Radiation (batch 22) | 0.914 | 0 | Up: 0.0305<br>Down: 0.03 | 10 |
| (Gr 2): C57BL/6J Cytostatica bestraling Sham (batch 22) | 0.9632 | 0 | Up: 0.0408<br>Down: 0.0399 | 9 |

| | |
|---|---|
| **T-test:** | -0.9776 |
| **P-value:** | 0.3433 |
| **Df:** | 15.4572 |
| **Mann-Whitney U test:** | 0.447 |

| Mice excluded by Quality Control |
|---|
| PH06469 (C57BL/6J - ) |

 © Sylics BV

Long arrest duration - habituation ratio light

*Long arrest duration - habituation ratio light*: Habituation effect: Change in cummulative long arrest long arrest duration during the light phase of day 3 compared to the light phase of day 1

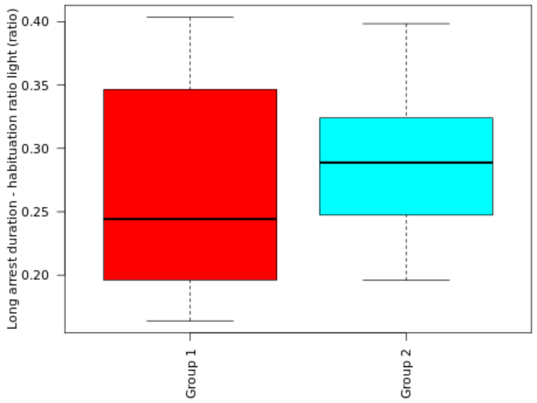

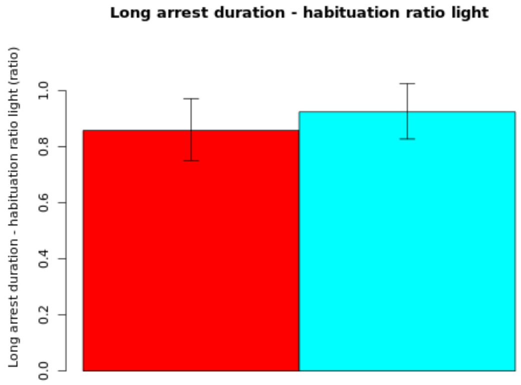

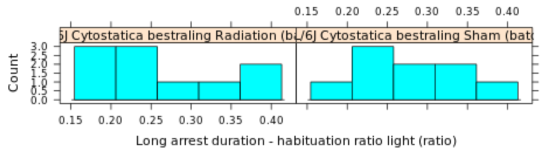

**NOTE: The statistics and boxplot are performed on log10 transformed data**

| Genotype | Mean | Median | SEM | N |
|---|---|---|---|---|
| (Gr 1): C57BL/6J Cytostatica bestraling Radiation (batch 22) | 0.858 | 0 | Up: 0.1138<br>Down: 0.1073 | 10 |
| (Gr 2): C57BL/6J Cytostatica bestraling Sham (batch 22) | 0.9242 | 0 | Up: 0.1015<br>Down: 0.0964 | 9 |

| | |
|---|---|
| **T-test:** | -0.445 |
| **P-value:** | 0.662 |
| **Df:** | 16.8741 |
| **Mann-Whitney U test:** | 0.6607 |

| Mice excluded by Quality Control |
|---|
| PH06469 (C57BL/6J - ) |

 © Sylics BV

Mean long arrest duration - habituation ratio light

*Mean long arrest duration - habituation ratio light*: Habituation effect: Change in mean long arrest duration during the light phase of day 3 compared to the light phase of day 1

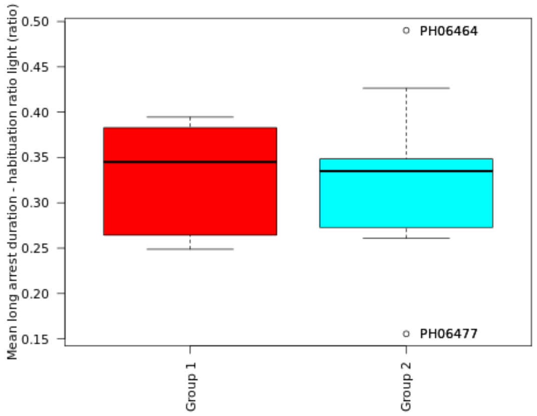

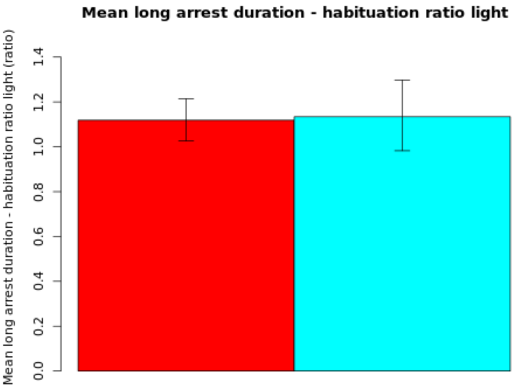

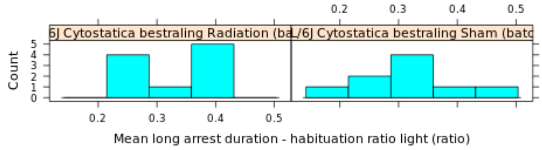

**NOTE: The statistics and boxplot are performed on log10 transformed data**

| Genotype | Mean | Median | SEM | N |
|---|---|---|---|---|
| (Gr 1): C57BL/6J Cytostatica bestraling Radiation (batch 22) | 1.1183 | 0 | Up: 0.0963<br>Down: 0.0921 | 10 |
| (Gr 2): C57BL/6J Cytostatica bestraling Sham (batch 22) | 1.134 | 0 | Up: 0.1634<br>Down: 0.1518 | 9 |

| | |
|---|---|
| **T-test:** | -0.086 |
| **P-value:** | 0.9327 |
| **Df:** | 13.3059 |
| **Mann-Whitney U test:** | 0.9682 |

| Mice excluded by Quality Control |
|---|
| PH06469 (C57BL/6J - ) |

 © Sylics BV

Long arrest number - habituation ratio light

*Long arrest number - habituation ratio light*: Habituation effect: Change in cummulative long arrest long arrest number during the light phase of day 3 compared to the light phase of day 1

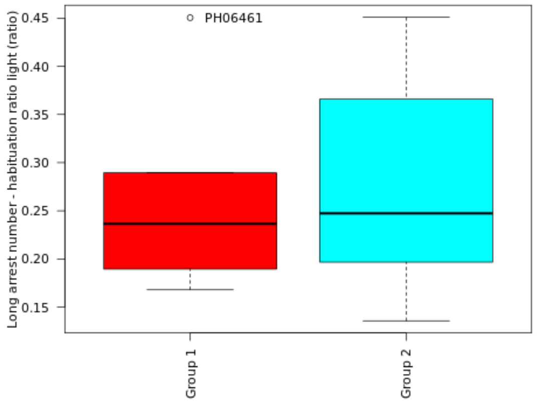

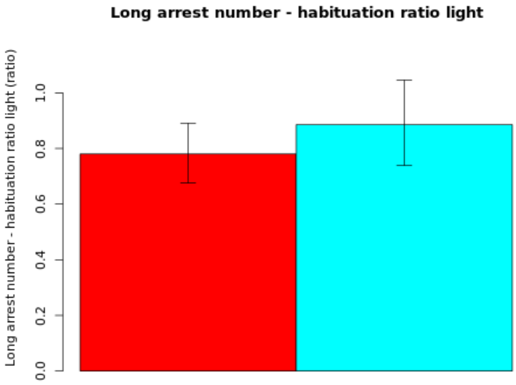

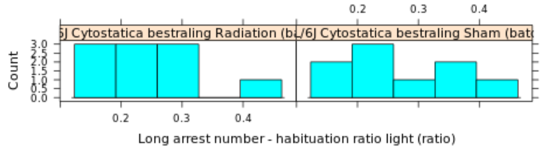

**NOTE: The statistics and boxplot are performed on log10 transformed data**

| Genotype | Mean | Median | SEM | N |
|---|---|---|---|---|
| (Gr 1): C57BL/6J Cytostatica bestraling Radiation (batch 22) | 0.7802 | 0 | Up: 0.1109<br>Down: 0.1044 | 10 |
| (Gr 2): C57BL/6J Cytostatica bestraling Sham (batch 22) | 0.886 | 0 | Up: 0.16<br>Down: 0.1475 | 9 |

| | |
|---|---|
| **T-test:** | -0.5693 |
| **P-value:** | 0.5775 |
| **Df:** | 15.1564 |
| **Mann-Whitney U test:** | 0.549 |

| Mice excluded by Quality Control |
|---|
| PH06469 (C57BL/6J - ) |

 © Sylics BV

Feeding zone duration - habituation ratio light

*Feeding zone duration - habituation ratio light*: Habituation effect: Change in cummulative Feeding zone duration during the light phase of day 3 compared to the light phase of day 1

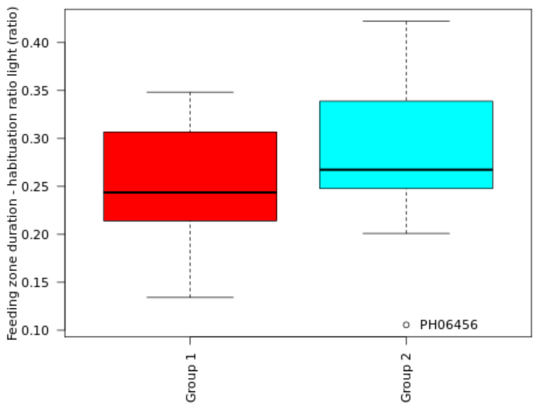

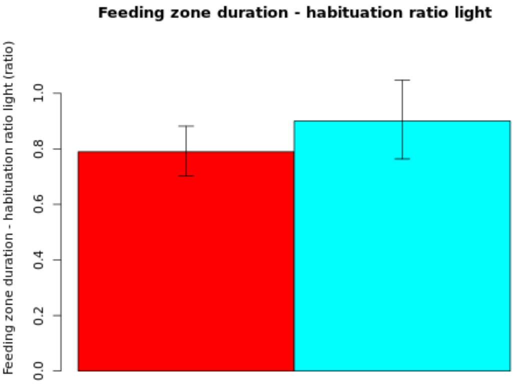

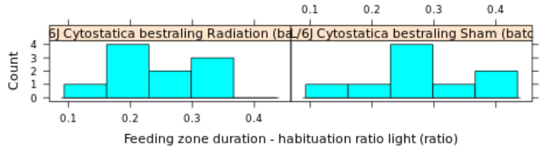

**NOTE: The statistics and boxplot are performed on log10 transformed data**

| Genotype | Mean | Median | SEM | N |
|---|---|---|---|---|
| (Gr 1): C57BL/6J Cytostatica bestraling Radiation (batch 22) | 0.7897 | 0 | Up: 0.092<br>Down: 0.0875 | 10 |
| (Gr 2): C57BL/6J Cytostatica bestraling Sham (batch 22) | 0.9006 | 0 | Up: 0.1473<br>Down: 0.1367 | 9 |

| | |
|---|---|
| **T-test:** | -0.6687 |
| **P-value:** | 0.5144 |
| **Df:** | 14.2685 |
| **Mann-Whitney U test:** | 0.6038 |

| Mice excluded by Quality Control |
|---|
| PH06469 (C57BL/6J - ) |

Mean short shelter visit duration - habituation ratio light

*Mean short shelter visit duration - habituation ratio light*: Habituation effect: Change in mean short shelter visit duration during the light phase of day 3 compared to the light phase of day 1

**NOTE: The statistics and boxplot are performed on log10 transformed data**
**Not enough data to show results**

Long shelter visit duration - habituation ratio light

*Long shelter visit duration - habituation ratio light*: Habituation effect: Change in cummulative long shelter visit duration during the light phase of day 3 compared to the light phase of day 1

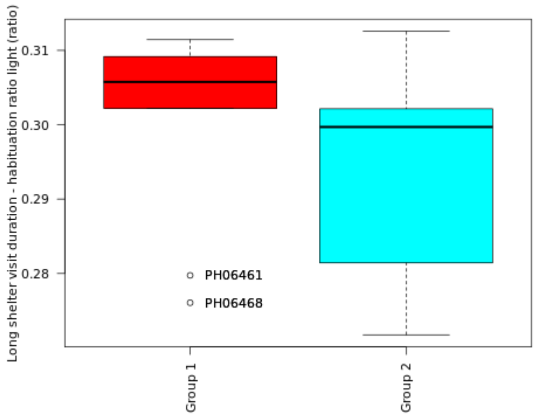

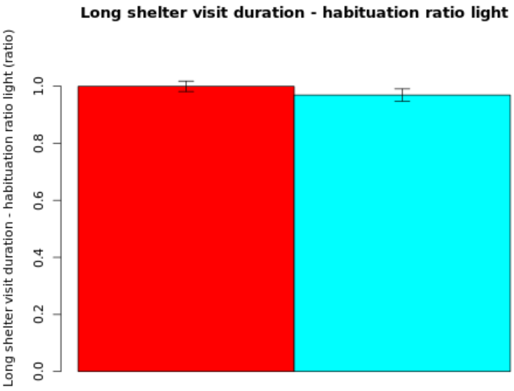

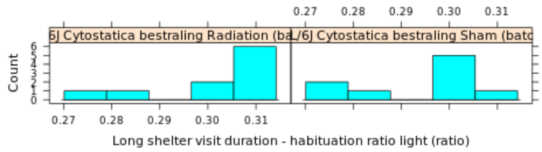

**NOTE: The statistics and boxplot are performed on log10 transformed data**

| Genotype | Mean | Median | SEM | N |
|---|---|---|---|---|
| (Gr 1): C57BL/6J Cytostatica bestraling Radiation (batch 22) | 0.9994 | 0 | Up: 0.0183<br>Down: 0.0181 | 10 |
| (Gr 2): C57BL/6J Cytostatica bestraling Sham (batch 22) | 0.969 | 0 | Up: 0.0216<br>Down: 0.0214 | 9 |

| | |
|---|---|
| **T-test:** | 1.0785 |
| **P-value:** | 0.2967 |
| **Df:** | 16.083 |
| **Mann-Whitney U test:** | 0.0947 |

| Mice excluded by Quality Control |
|---|
| PH06469 (C57BL/6J - ) |

 © Sylics BV

Mean long movement distance - habituation ratio light

*Mean long movement distance - habituation ratio light*: Habituation effect: Change in mean long movement distance during the light phase of day 3 compared to the light phase of day 1

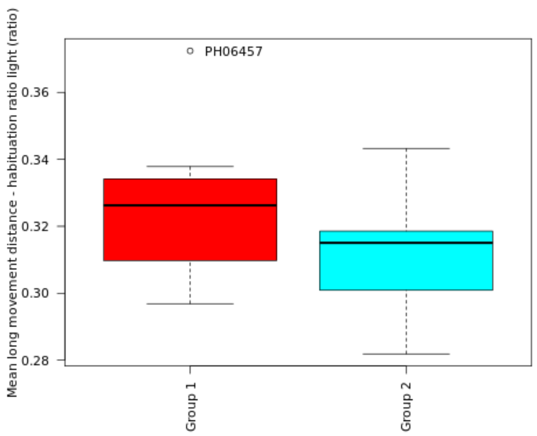

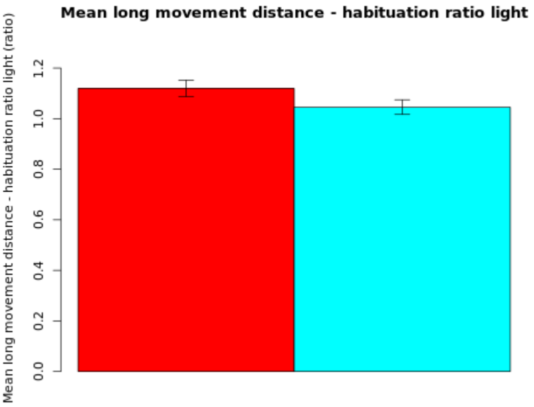

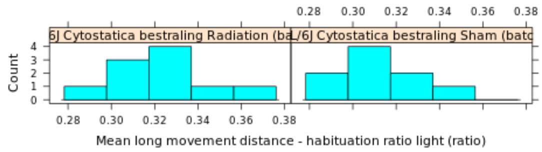

**NOTE: The statistics and boxplot are performed on log10 transformed data**

| Genotype | Mean | Median | SEM | N |
|---|---|---|---|---|
| (Gr 1): C57BL/6J Cytostatica bestraling Radiation (batch 22) | 1.1197 | 0 | Up: 0.0324<br>Down: 0.0319 | 10 |
| (Gr 2): C57BL/6J Cytostatica bestraling Sham (batch 22) | 1.0454 | 0 | Up: 0.0291<br>Down: 0.0287 | 9 |

| | |
|---|---|
| **T-test:** | 1.722 |
| **P-value:** | 0.1032 |
| **Df:** | 16.9973 |
| **Mann-Whitney U test:** | 0.1564 |

| Mice excluded by Quality Control |
|---|
| PH06469 (C57BL/6J - ) |

*© Sylics BV*

Mean short movement distance - habituation ratio light

*Mean short movement distance - habituation ratio light*: Habituation effect: Change in mean short movement distance during the light phase of day 3 compared to the light phase of day 1

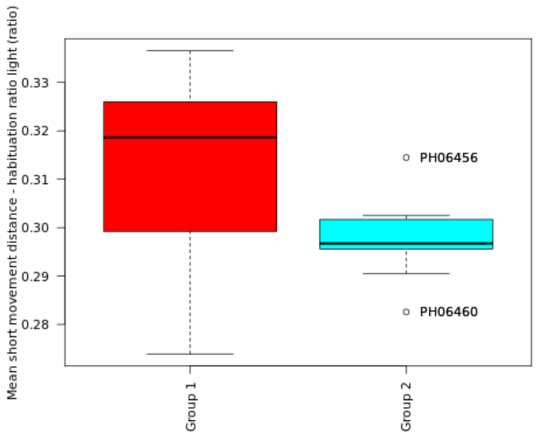

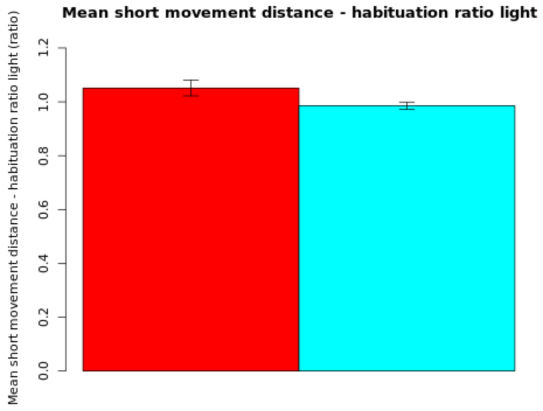

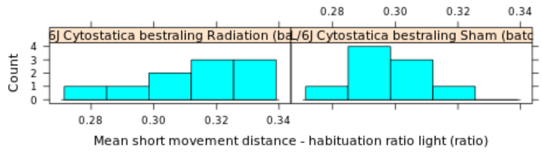

**NOTE: The statistics and boxplot are performed on log10 transformed data**

| Genotype | Mean | Median | SEM | N |
|---|---|---|---|---|
| (Gr 1): C57BL/6J Cytostatica bestraling Radiation (batch 22) | 1.0512 | 0 | Up: 0.0296<br>Down: 0.0292 | 10 |
| (Gr 2): C57BL/6J Cytostatica bestraling Sham (batch 22) | 0.9856 | 0 | Up: 0.0135<br>Down: 0.0134 | 9 |

| | |
|---|---|
| **T-test:** | 2.0509 |
| **P-value:** | 0.0614 |
| **Df:** | 12.7296 |
| **Mann-Whitney U test:** | 0.0653 |

| Mice excluded by Quality Control |
|---|
| PH06469 (C57BL/6J - ) |

 © Sylics BV

OnShelter zone duration - habituation ratio light

*OnShelter zone duration - habituation ratio light*: Habituation effect: Change in cummulative OnShelter zone duration during the light phase of day 3 compared to the light phase of day 1

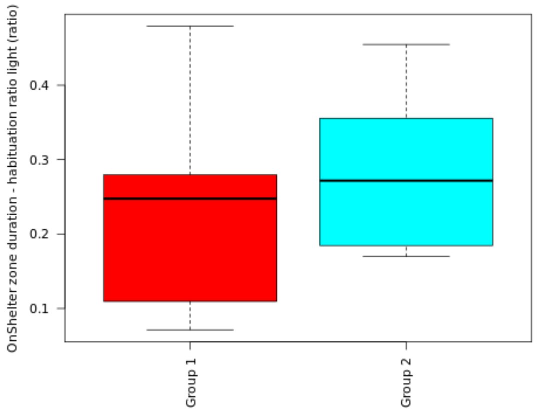

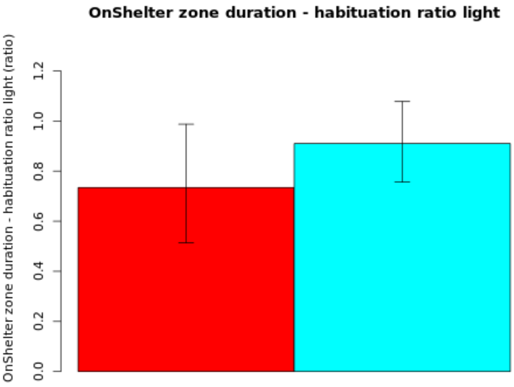

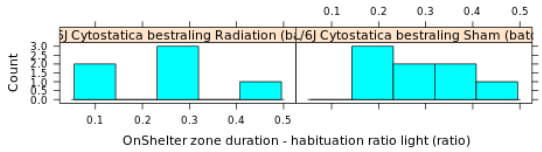

**NOTE: The statistics and boxplot are performed on log10 transformed data**

| Genotype | Mean | Median | SEM | N |
|---|---|---|---|---|
| (Gr 1): C57BL/6J Cytostatica bestraling Radiation (batch 22) | 0.7344 | 0 | Up: 0.2529<br>Down: 0.2207 | 6 |
| (Gr 2): C57BL/6J Cytostatica bestraling Sham (batch 22) | 0.9099 | 0 | Up: 0.1676<br>Down: 0.1541 | 8 |

| | |
|---|---|
| **T-test:** | -0.6024 |
| **P-value:** | 0.5624 |
| **Df:** | 8.6473 |
| **Mann-Whitney U test:** | 0.5728 |

| Mice excluded by Quality Control |
|---|
| PH06469 (C57BL/6J - ) |

Sylics

Spout zone duration - habituation ratio light

*Spout zone duration - habituation ratio light*: Habituation effect: Change in cummulative Spout zone duration during the light phase of day 3 compared to the light phase of day 1

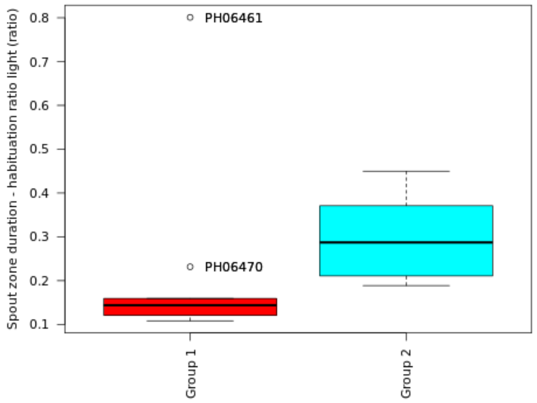

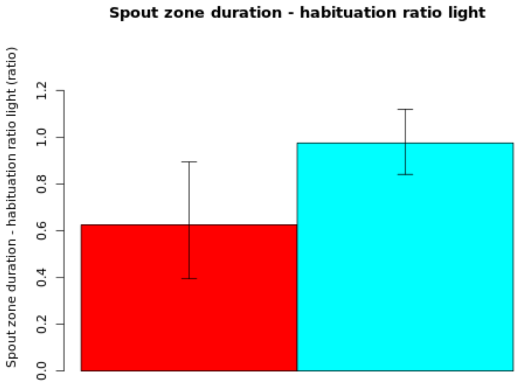

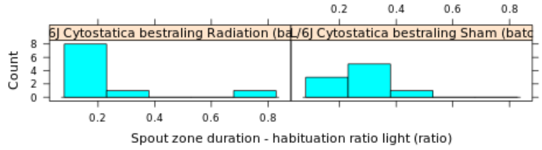

**NOTE: The statistics and boxplot are performed on log10 transformed data**

| Genotype | Mean | Median | SEM | N |
|---|---|---|---|---|
| (Gr 1): C57BL/6J Cytostatica bestraling Radiation (batch 22) | 0.6251 | 0 | Up: 0.2689<br>Down: 0.2307 | 10 |
| (Gr 2): C57BL/6J Cytostatica bestraling Sham (batch 22) | 0.9754 | 0 | Up: 0.1444<br>Down: 0.1345 | 9 |

| | |
|---|---|
| **T-test:** | -1.1577 |
| **P-value:** | 0.2685 |
| **Df:** | 12.5867 |
| **Mann-Whitney U test:** | 0.0057 |

| Mice excluded by Quality Control |
|---|
| PH06469 (C57BL/6J - ) |

 © Sylics BV

### Spontaneous behavior (Group 4: DarkLight index)

With respect to spontaneous behaviors in the first three days in the PhenoTyper, 6 groups of behavioral parameters are defined as described below. The first two groups describe specific behavioral elements related to kinematics of mice (description of movement characteristics, group1) and sheltering behavior (group 2). These behavioral parameters were analyzed with respect to temporal aspects, in particular over 4 different time scales, i.e., habituation effects across multiple days (group 3), effects of DarkLight phase across 24h (group 4), differences in the pattern of behavior in the few hours before and after phase shifts (group5), and differences in activity bout properties on the sub-minute time scale (group 6).

**Group 4: DarkLight index**
The light/dark cycle has a strong impact on behavior during each day of the experiment. To assess whether a given behavior is more prominent during the dark or light phase, we defined a LightDark index (dark value/(dark value + light value)). This LightDark index lies between 0.5 and 1.0 if a specific behavior is more pronounced during the dark phase and between 0.0 and 0.5 if it is more prominent during the light phase.

 *© Sylics BV*

Activity duration - darklight index

*Activity duration - darklight index*: Effect of light regime: Index describing the difference in cummulative activity duration between light and dark phase. If this index is between 0.5 and 1 then parameter values are higher during the dark phase. If this index is between 0 and 0.5 then parameter values are higher during the light phase.

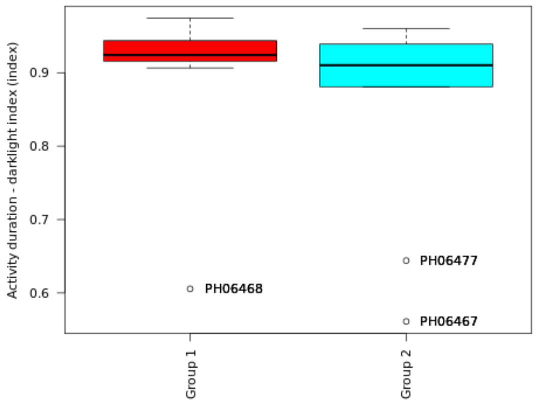

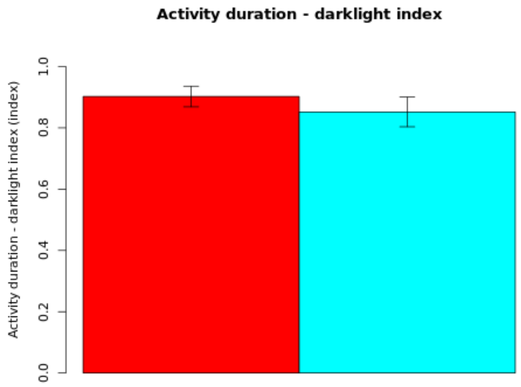

**Secondary plot:**

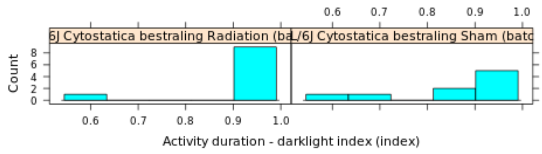

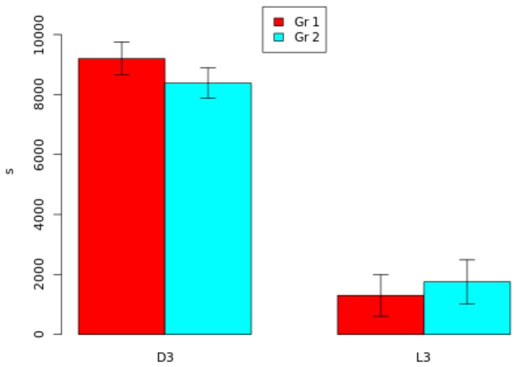

| Genotype | Mean | Median | SEM | N |
|---|---|---|---|---|
| (Gr 1): C57BL/6J Cytostatica bestraling Radiation (batch 22) | 0.9024 | 0.9244 | 0.0337 | 10 |
| (Gr 2): C57BL/6J Cytostatica bestraling Sham (batch 22) | 0.8518 | 0.9103 | 0.0483 | 9 |

| | |
|---|---|
| **T-test:** | 0.8588 |
| **P-value:** | 0.4043 |
| **Df:** | 14.6083 |
| **Mann-Whitney U test:** | 0.4002 |

| Mice excluded by Quality Control |
|---|
| PH06469 (C57BL/6J - ) |

 © Sylics BV

Mean activity duration - darklight index

*Mean activity duration - darklight index*: Effect of light regime: Index describing the difference in mean activity duration between light and dark phase. If this index is between 0.5 and 1 then parameter values are higher during the dark phase. If this index is between 0 and 0.5 then parameter values are higher during the light phase.

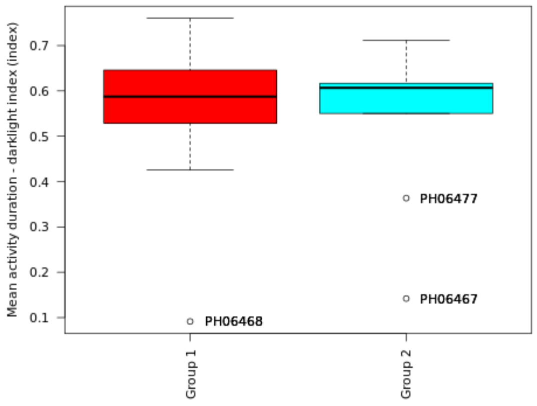

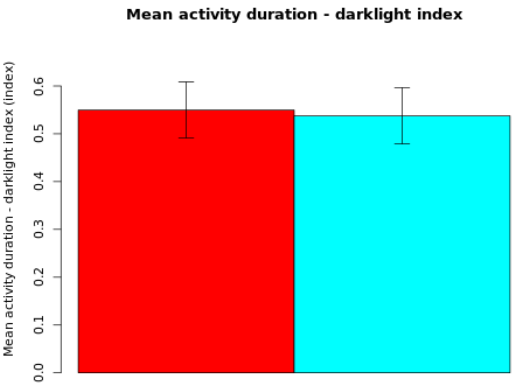

**Secondary plot:**

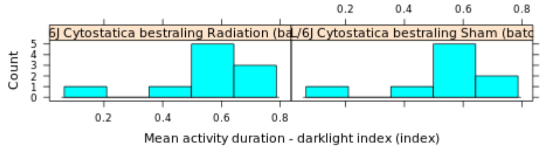

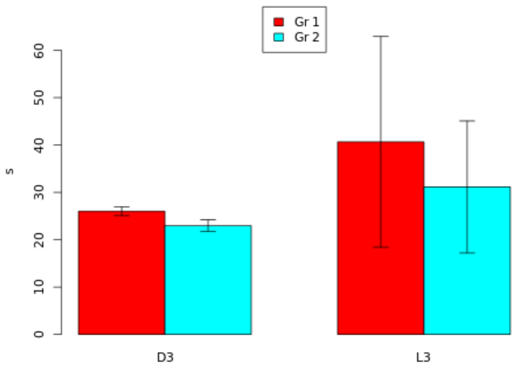

| Genotype | Mean | Median | SEM | N |
|---|---|---|---|---|
| (Gr 1): C57BL/6J Cytostatica bestraling Radiation (batch 22) | 0.5498 | 0.5873 | 0.0586 | 10 |
| (Gr 2): C57BL/6J Cytostatica bestraling Sham (batch 22) | 0.5376 | 0.6068 | 0.0589 | 9 |

| | |
|---|---|
| **T-test:** | 0.1467 |
| **P-value:** | 0.8851 |
| **Df:** | 16.929 |
| **Mann-Whitney U test:** | 1 |

| Mice excluded by Quality Control |
|---|
| PH06469 (C57BL/6J - ) |

 © Sylics BV

Activity number - darklight index

*Activity number - darklight index*: Effect of light regime: Index describing the difference in cummulative activity number between light and dark phase. If this index is between 0.5 and 1 then parameter values are higher during the dark phase. If this index is between 0 and 0.5 then parameter values are higher during the light phase.

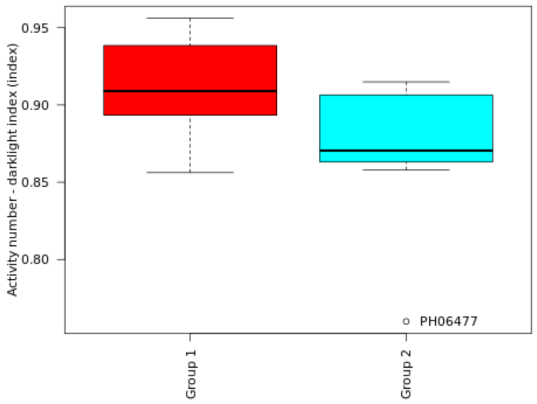

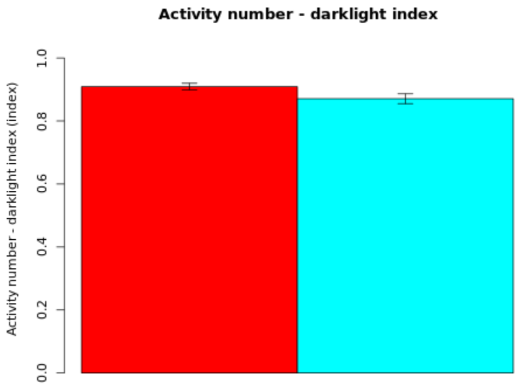

**Secondary plot:**

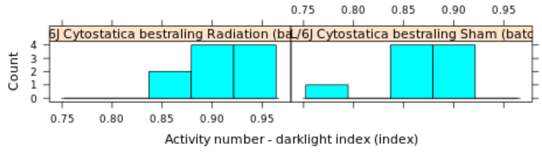

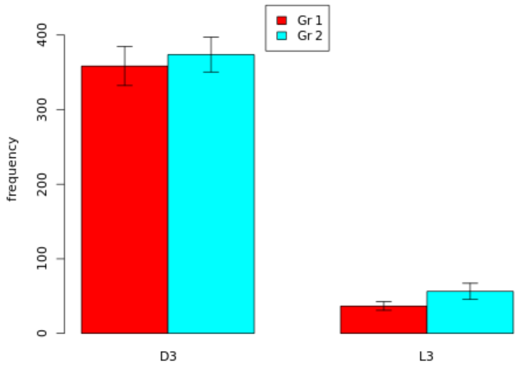

| Genotype | Mean | Median | SEM | N |
|---|---|---|---|---|
| (Gr 1): C57BL/6J Cytostatica bestraling Radiation (batch 22) | 0.9091 | 0.909 | 0.0103 | 10 |
| (Gr 2): C57BL/6J Cytostatica bestraling Sham (batch 22) | 0.8705 | 0.8705 | 0.0155 | 9 |

| | |
|---|---|
| **T-test:** | 2.0699 |
| **P-value:** | 0.0572 |
| **Df:** | 14.1532 |
| **Mann-Whitney U test:** | 0.0789 |

| Mice excluded by Quality Control |
|---|
| PH06469 (C57BL/6J - ) |

 *© Sylics BV*

Mean short arrest duration - darklight index

*Mean short arrest duration - darklight index*: Effect of light regime: Index describing the difference in mean short arrest duration between light and dark phase. If this index is between 0.5 and 1 then parameter values are higher during the dark phase. If this index is between 0 and 0.5 then parameter values are higher during the light phase.

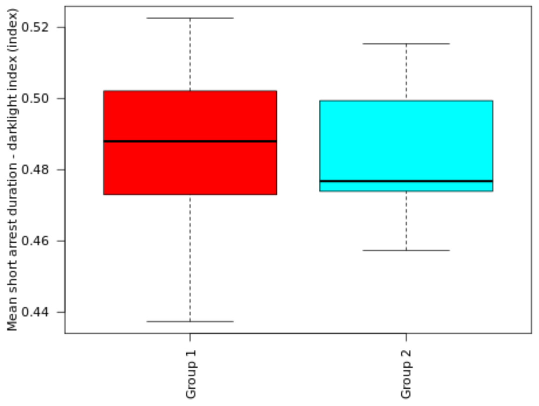

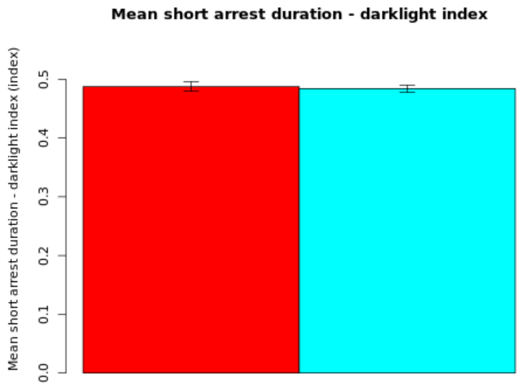

**Secondary plot:**

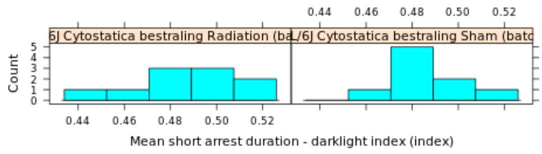

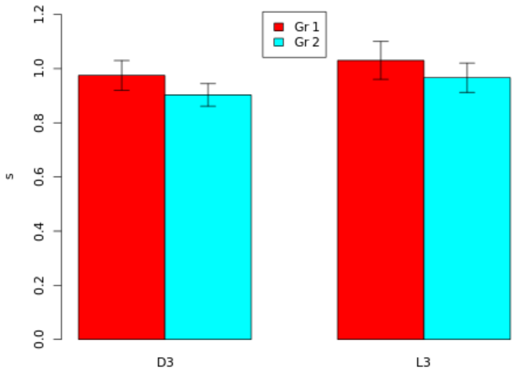

| Genotype | Mean | Median | SEM | N |
|---|---|---|---|---|
| (Gr 1): C57BL/6J Cytostatica bestraling Radiation (batch 22) | 0.4876 | 0.488 | 0.0081 | 10 |
| (Gr 2): C57BL/6J Cytostatica bestraling Sham (batch 22) | 0.484 | 0.4768 | 0.006 | 9 |

| | |
|---|---|
| **T-test:** | 0.3622 |
| **P-value:** | 0.7219 |
| **Df:** | 16.1265 |
| **Mann-Whitney U test:** | 0.6607 |

| Mice excluded by Quality Control |
|---|
| PH06469 (C57BL/6J - ) |

 © Sylics BV

Long arrest duration - darklight index

*Long arrest duration - darklight index*: Effect of light regime: Index describing the difference in cummulative long arrest long arrest duration between light and dark phase. If this index is between 0.5 and 1 then parameter values are higher during the dark phase. If this index is between 0 and 0.5 then parameter values are higher during the light phase.

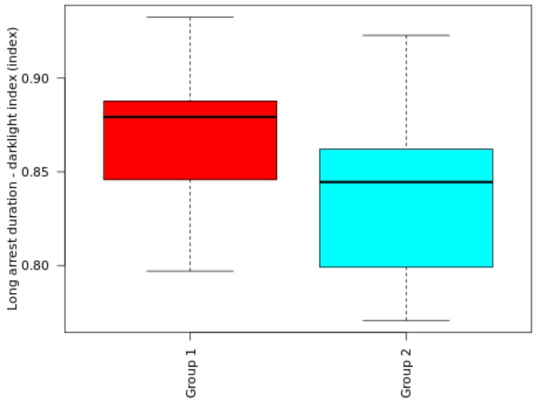

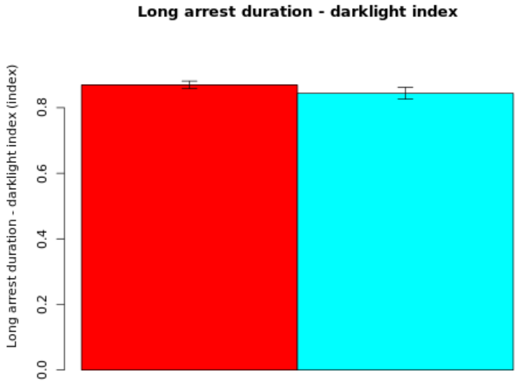

**Secondary plot:**

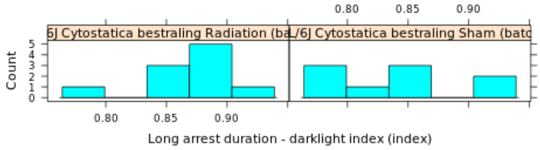

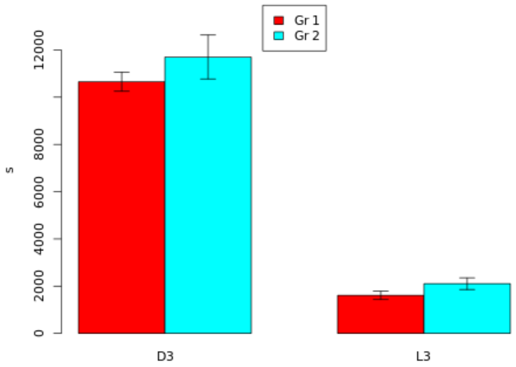

| Genotype | Mean | Median | SEM | N |
|---|---|---|---|---|
| (Gr 1): C57BL/6J Cytostatica bestraling Radiation (batch 22) | 0.8696 | 0.8793 | 0.0114 | 10 |
| (Gr 2): C57BL/6J Cytostatica bestraling Sham (batch 22) | 0.8441 | 0.8445 | 0.0176 | 9 |

| | |
|---|---|
| **T-test:** | 1.2105 |
| **P-value:** | 0.2462 |
| **Df:** | 13.9628 |
| **Mann-Whitney U test:** | 0.211 |

| Mice excluded by Quality Control |
|---|
| PH06469 (C57BL/6J - ) |

 © Sylics BV

Mean long arrest duration - darklight index

*Mean long arrest duration - darklight index*: Effect of light regime: Index describing the difference in mean long arrest duration between light and dark phase. If this index is between 0.5 and 1 then parameter values are higher during the dark phase. If this index is between 0 and 0.5 then parameter values are higher during the light phase.

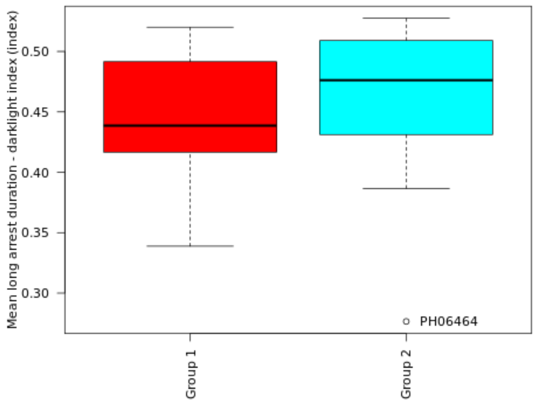

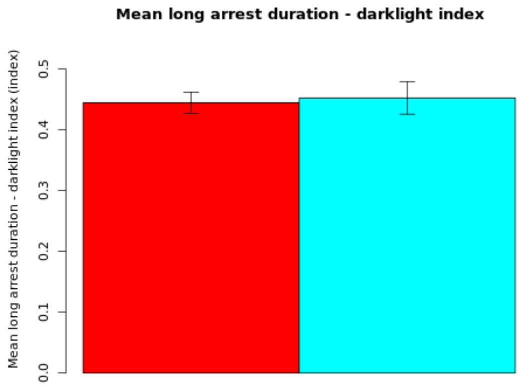

**Secondary plot:**

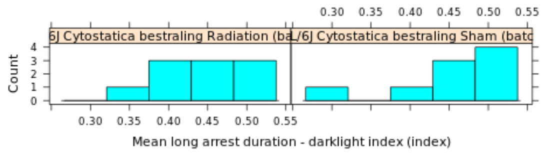

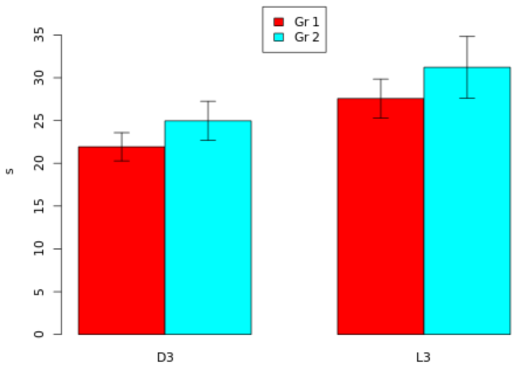

| Genotype | Mean | Median | SEM | N |
|---|---|---|---|---|
| (Gr 1): C57BL/6J Cytostatica bestraling Radiation (batch 22) | 0.4446 | 0.4386 | 0.0174 | 10 |
| (Gr 2): C57BL/6J Cytostatica bestraling Sham (batch 22) | 0.4525 | 0.4762 | 0.0267 | 9 |

| | |
|---|---|
| **T-test:** | -0.2466 |
| **P-value:** | 0.8088 |
| **Df:** | 14.0221 |
| **Mann-Whitney U test:** | 0.549 |

| Mice excluded by Quality Control |
|---|
| PH06469 (C57BL/6J - ) |

 © Sylics BV

Long arrest number - darklight index

*Long arrest number - darklight index*: Effect of light regime: Index describing the difference in cummulative long arrest long arrest number between light and dark phase. If this index is between 0.5 and 1 then parameter values are higher during the dark phase. If this index is between 0 and 0.5 then parameter values are higher during the light phase.

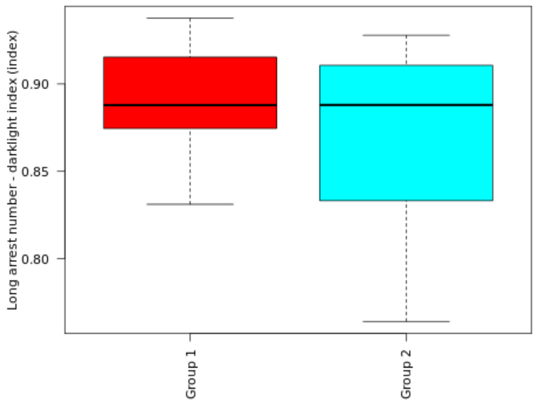

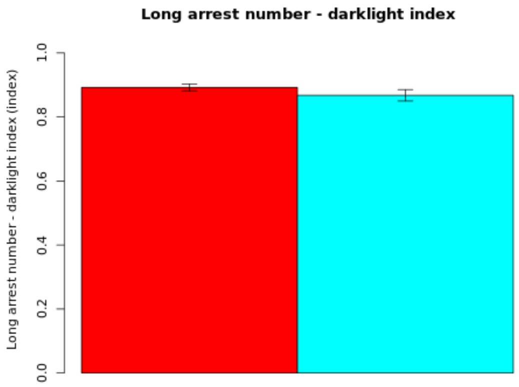

**Secondary plot:**

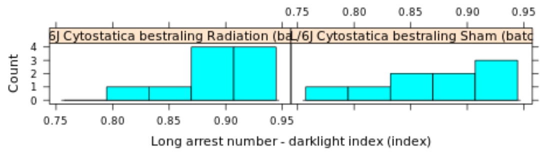

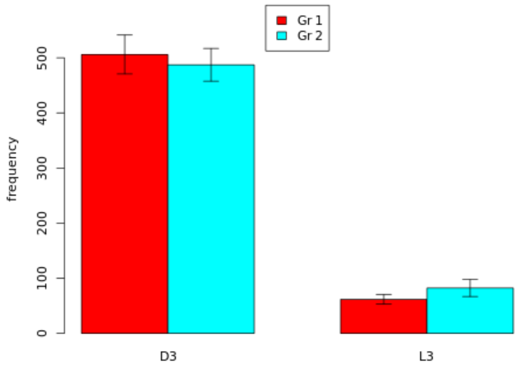

| Genotype | Mean | Median | SEM | N |
|---|---|---|---|---|
| (Gr 1): C57BL/6J Cytostatica bestraling Radiation (batch 22) | 0.8924 | 0.8879 | 0.0103 | 10 |
| (Gr 2): C57BL/6J Cytostatica bestraling Sham (batch 22) | 0.8677 | 0.888 | 0.0178 | 9 |

| | |
|---|---|
| **T-test:** | 1.2046 |
| **P-value:** | 0.2498 |
| **Df:** | 12.9905 |
| **Mann-Whitney U test:** | 0.447 |

| Mice excluded by Quality Control |
|---|
| PH06469 (C57BL/6J - ) |

 © Sylics BV

Feeding zone duration - darklight index

*Feeding zone duration - darklight index*: Effect of light regime: Index describing the difference in cummulative Feeding zone duration between light and dark phase. If this index is between 0.5 and 1 then parameter values are higher during the dark phase. If this index is between 0 and 0.5 then parameter values are higher during the light phase.

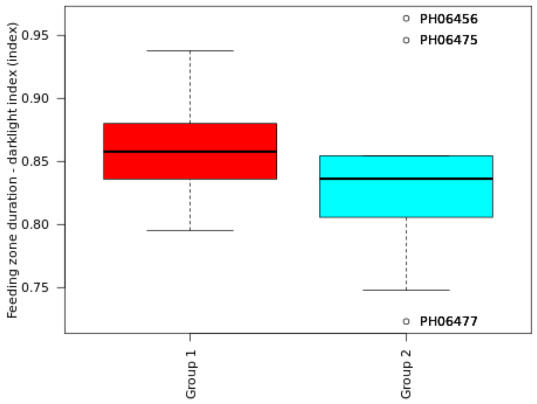

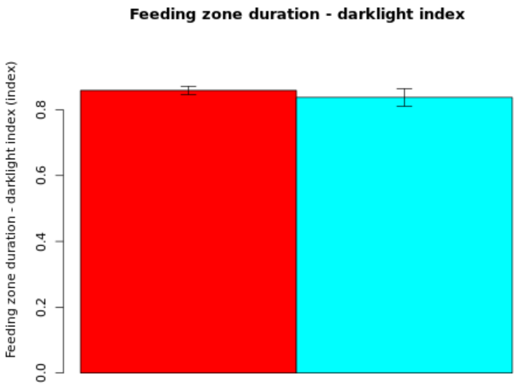

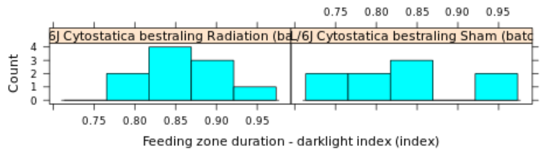

**Secondary plot:**

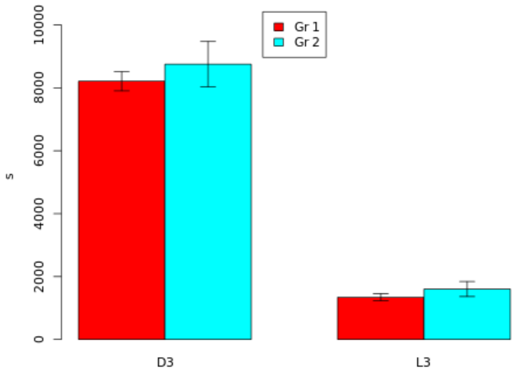

| Genotype | Mean | Median | SEM | N |
|---|---|---|---|---|
| (Gr 1): C57BL/6J Cytostatica bestraling Radiation (batch 22) | 0.8588 | 0.8578 | 0.0125 | 10 |
| (Gr 2): C57BL/6J Cytostatica bestraling Sham (batch 22) | 0.8376 | 0.8363 | 0.0266 | 9 |

| | |
|---|---|
| **T-test:** | 0.7226 |
| **P-value:** | 0.4845 |
| **Df:** | 11.4246 |
| **Mann-Whitney U test:** | 0.3562 |

| Mice excluded by Quality Control |
|---|
| PH06469 (C57BL/6J - ) |

Mean short shelter visit duration - darklight index

*Mean short shelter visit duration - darklight index*: Effect of light regime: Index describing the difference in mean short shelter visit duration between light and dark phase. If this index is between 0.5 and 1 then parameter values are higher during the dark phase. If this index is between 0 and 0.5 then parameter values are higher during the light phase.

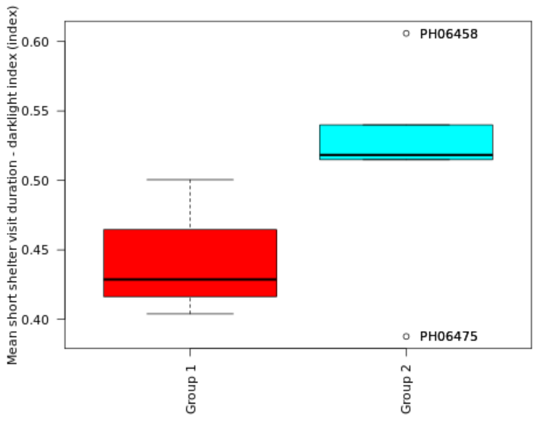

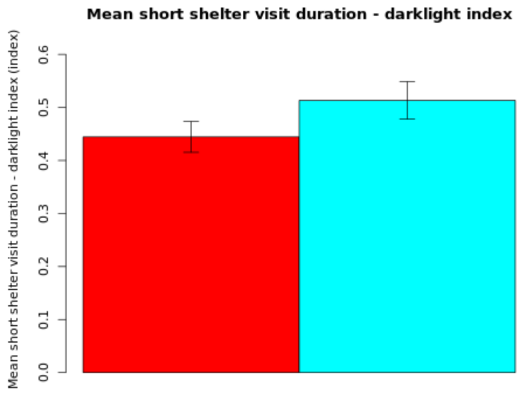

**Secondary plot:**

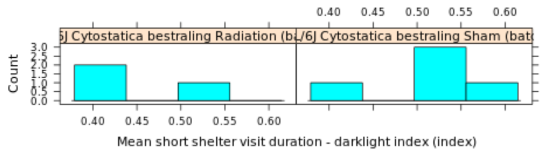

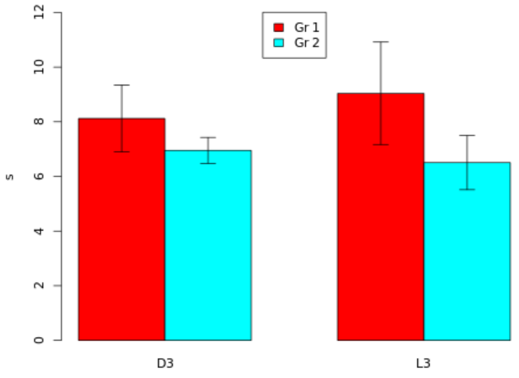

| Genotype | Mean | Median | SEM | N |
|---|---|---|---|---|
| (Gr 1): C57BL/6J Cytostatica bestraling Radiation (batch 22) | 0.4444 | 0.4287 | 0.029 | 3 |
| (Gr 2): C57BL/6J Cytostatica bestraling Sham (batch 22) | 0.5134 | 0.5183 | 0.0354 | 5 |

| | |
|---|---|
| **T-test:** | -1.5073 |
| **P-value:** | 0.1835 |
| **Df:** | 5.879 |
| **Mann-Whitney U test:** | 0.25 |

| Mice excluded by Quality Control |
|---|
| PH06469 (C57BL/6J - ) |

 © Sylics BV

Long shelter visit duration - darklight index

*Long shelter visit duration - darklight index*: Effect of light regime: Index describing the difference in cummulative long shelter visit duration between light and dark phase. If this index is between 0.5 and 1 then parameter values are higher during the dark phase. If this index is between 0 and 0.5 then parameter values are higher during the light phase.

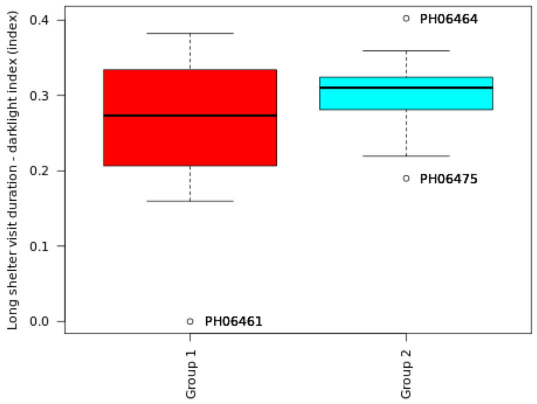

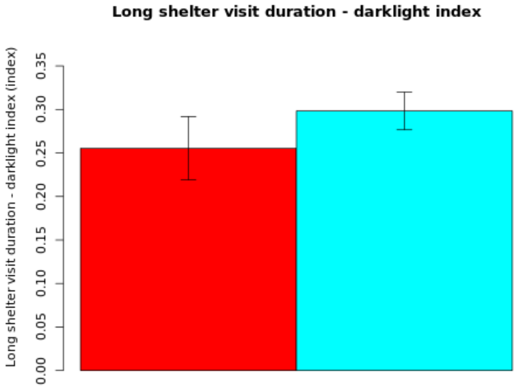

**Secondary plot:**

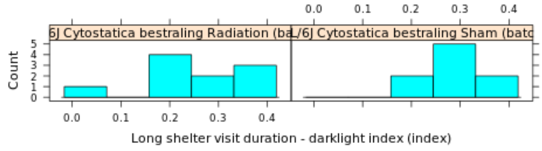

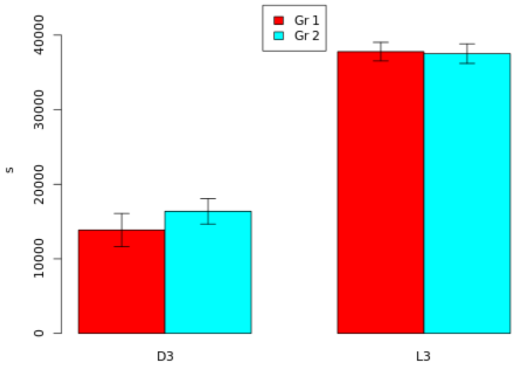

| Genotype | Mean | Median | SEM | N |
|---|---|---|---|---|
| (Gr 1): C57BL/6J Cytostatica bestraling Radiation (batch 22) | 0.2554 | 0.2734 | 0.0364 | 10 |
| (Gr 2): C57BL/6J Cytostatica bestraling Sham (batch 22) | 0.2983 | 0.3103 | 0.0217 | 9 |

| | |
|---|---|
| **T-test:** | -1.0115 |
| **P-value:** | 0.3284 |
| **Df:** | 14.4776 |
| **Mann-Whitney U test:** | 0.6038 |

| Mice excluded by Quality Control |
|---|
| PH06469 (C57BL/6J - ) |

 © Sylics BV

Long shelter visit number - darklight index

*Long shelter visit number - darklight index*: Effect of light regime: Index describing the difference in cummulative long shelter visit number between light and dark phase. If this index is between 0.5 and 1 then parameter values are higher during the dark phase. If this index is between 0 and 0.5 then parameter values are higher during the light phase.

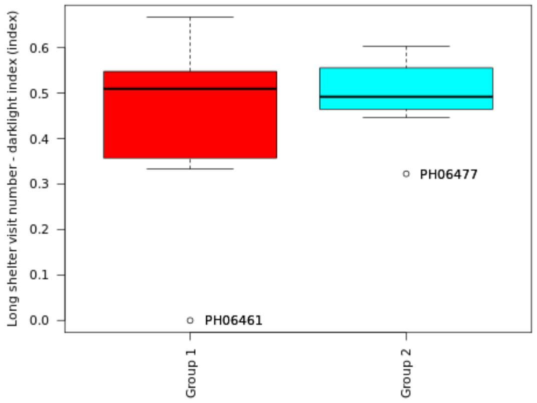

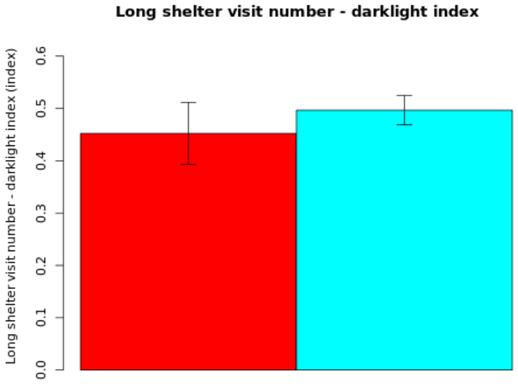

**Secondary plot:**

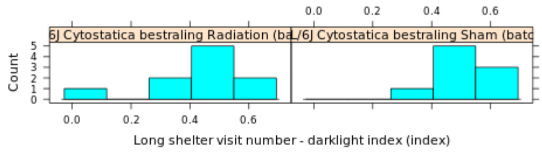

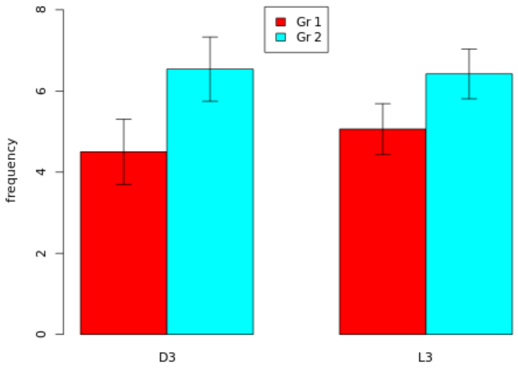

| Genotype | Mean | Median | SEM | N |
|---|---|---|---|---|
| (Gr 1): C57BL/6J Cytostatica bestraling Radiation (batch 22) | 0.4522 | 0.5096 | 0.0595 | 10 |
| (Gr 2): C57BL/6J Cytostatica bestraling Sham (batch 22) | 0.4967 | 0.4921 | 0.028 | 9 |

| | |
|---|---|
| **T-test:** | -0.6758 |
| **P-value:** | 0.5113 |
| **Df:** | 12.7264 |
| **Mann-Whitney U test:** | 0.8702 |

| Mice excluded by Quality Control |
|---|
| PH06469 (C57BL/6J - ) |

Mean long movement distance - darklight index

*Mean long movement distance - darklight index*: Effect of light regime: Index describing the difference in mean long movement distance between light and dark phase. If this index is between 0.5 and 1 then parameter values are higher during the dark phase. If this index is between 0 and 0.5 then parameter values are higher during the light phase.

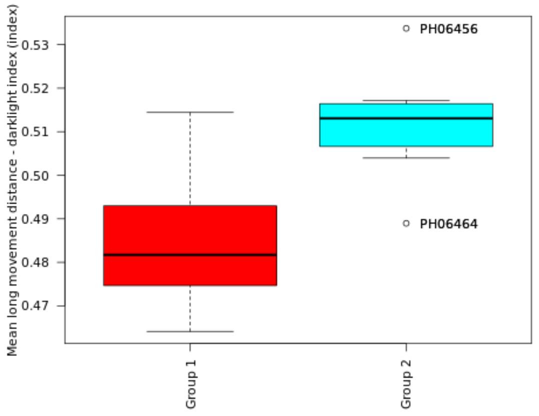

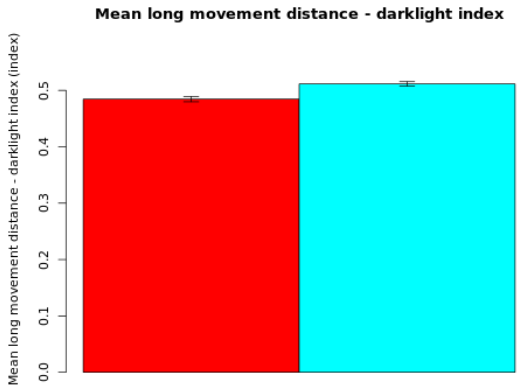

**Secondary plot:**

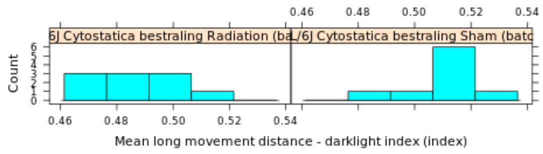

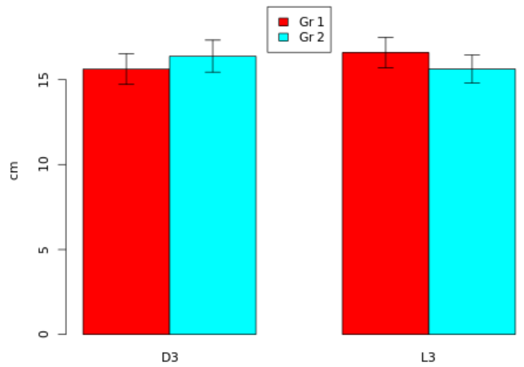

| Genotype | Mean | Median | SEM | N |
|---|---|---|---|---|
| (Gr 1): C57BL/6J Cytostatica bestraling Radiation (batch 22) | 0.4845 | 0.4817 | 0.0047 | 10 |
| (Gr 2): C57BL/6J Cytostatica bestraling Sham (batch 22) | 0.5114 | 0.5131 | 0.004 | 9 |

| | |
|---|---|
| **T-test:** | -4.368 |
| **P-value:** | 0.0004 |
| **Df:** | 16.8747 |
| **Mann-Whitney U test:** | 0.0015 |

| Mice excluded by Quality Control |
|---|
| PH06469 (C57BL/6J - ) |

 © Sylics BV

Mean short movement distance - darklight index

*Mean short movement distance - darklight index*: Effect of light regime: Index describing the difference in mean short movement distance between light and dark phase. If this index is between 0.5 and 1 then parameter values are higher during the dark phase. If this index is between 0 and 0.5 then parameter values are higher during the light phase.

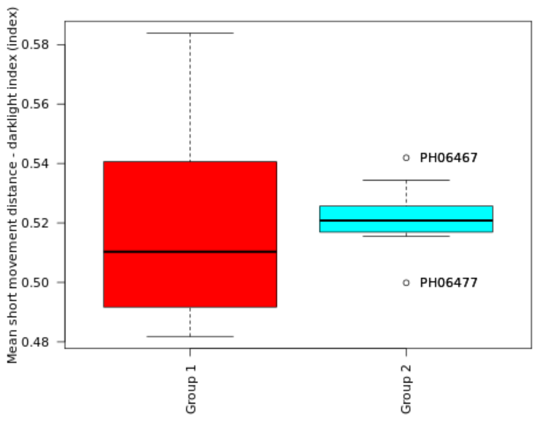

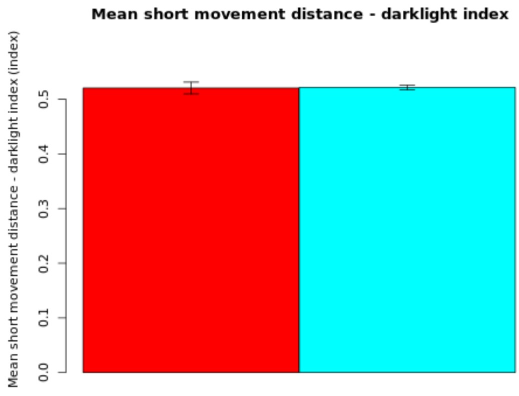

**Secondary plot:**

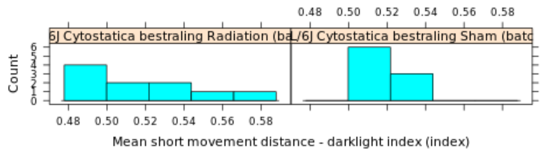

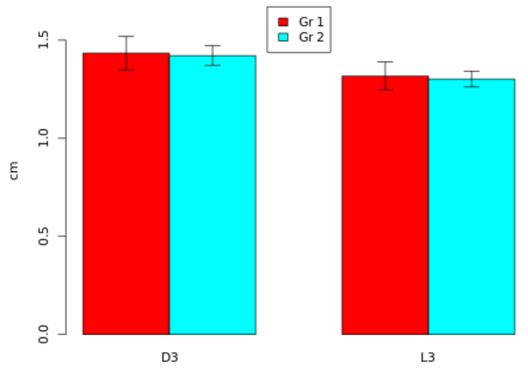

| Genotype | Mean | Median | SEM | N |
|---|---|---|---|---|
| (Gr 1): C57BL/6J Cytostatica bestraling Radiation (batch 22) | 0.5206 | 0.5103 | 0.0108 | 10 |
| (Gr 2): C57BL/6J Cytostatica bestraling Sham (batch 22) | 0.5216 | 0.5208 | 0.004 | 9 |

| | |
|---|---|
| **T-test:** | -0.0892 |
| **P-value:** | 0.9304 |
| **Df:** | 11.3907 |
| **Mann-Whitney U test:** | 0.6607 |

| Mice excluded by Quality Control |
|---|
| PH06469 (C57BL/6J - ) |

 © Sylics BV

OnShelter zone duration - darklight index

*OnShelter zone duration - darklight index*: Effect of light regime: Index describing the difference in cummulative OnShelter zone duration between light and dark phase. If this index is between 0.5 and 1 then parameter values are higher during the dark phase. If this index is between 0 and 0.5 then parameter values are higher during the light phase.

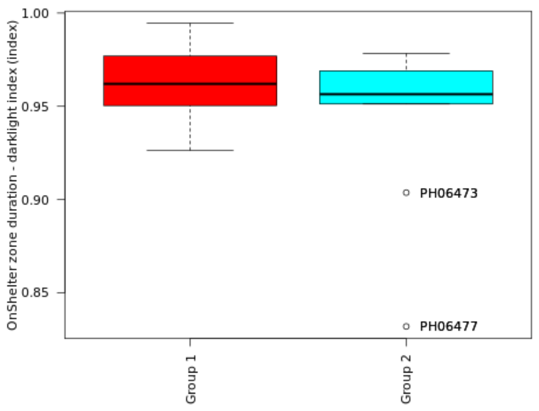

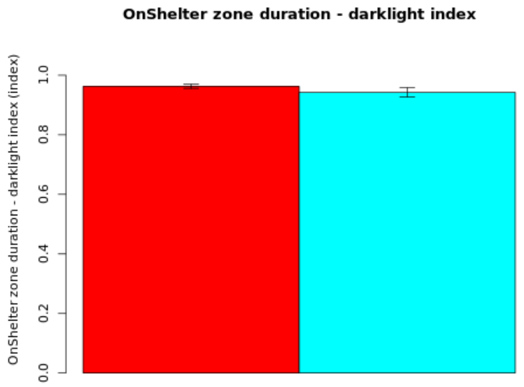

**Secondary plot:**

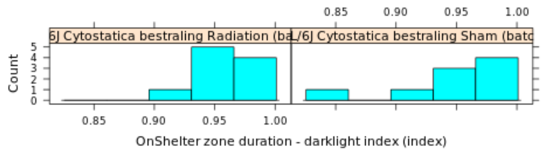

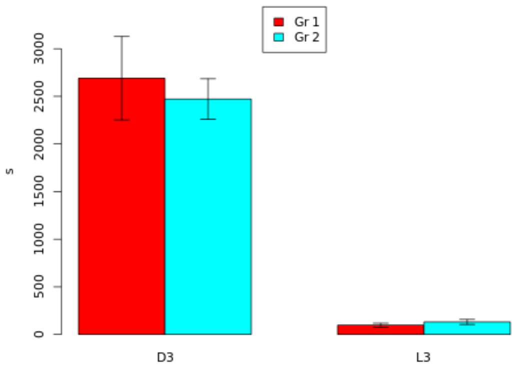

| Genotype | Mean | Median | SEM | N |
|---|---|---|---|---|
| (Gr 1): C57BL/6J Cytostatica bestraling Radiation (batch 22) | 0.9628 | 0.962 | 0.0067 | 10 |
| (Gr 2): C57BL/6J Cytostatica bestraling Sham (batch 22) | 0.9427 | 0.9565 | 0.0157 | 9 |

| | |
|---|---|
| **T-test:** | 1.1767 |
| **P-value:** | 0.2644 |
| **Df:** | 10.8919 |
| **Mann-Whitney U test:** | 0.6038 |

| Mice excluded by Quality Control |
|---|
| PH06469 (C57BL/6J - ) |

 © Sylics BV

Spout zone duration - darklight index

*Spout zone duration - darklight index*: Effect of light regime: Index describing the difference in cummulative Spout zone duration between light and dark phase. If this index is between 0.5 and 1 then parameter values are higher during the dark phase. If this index is between 0 and 0.5 then parameter values are higher during the light phase.

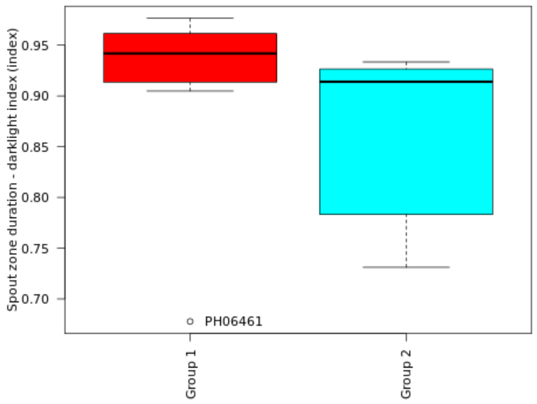

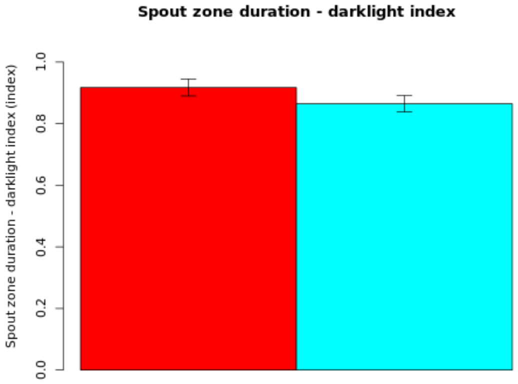

**Secondary plot:**

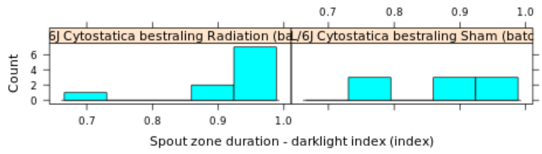

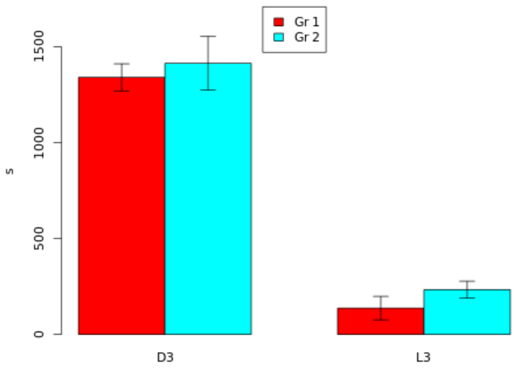

| Genotype | Mean | Median | SEM | N |
|---|---|---|---|---|
| (Gr 1): C57BL/6J Cytostatica bestraling Radiation (batch 22) | 0.9173 | 0.9419 | 0.0275 | 10 |
| (Gr 2): C57BL/6J Cytostatica bestraling Sham (batch 22) | 0.8644 | 0.914 | 0.0267 | 9 |

| | |
|---|---|
| **T-test:** | 1.3791 |
| **P-value:** | 0.1858 |
| **Df:** | 16.9874 |
| **Mann-Whitney U test:** | 0.035 |

| Mice excluded by Quality Control |
|---|
| PH06469 (C57BL/6J - ) |

 © Sylics BV

### Spontaneous behavior (Group 5: activity pattern)

With respect to spontaneous behaviors in the first three days in the PhenoTyper, 6 groups of behavioral parameters are defined as described below. The first two groups describe specific behavioral elements related to kinematics of mice (description of movement characteristics, group1) and sheltering behavior (group 2). These behavioral parameters were analyzed with respect to temporal aspects, in particular over 4 different time scales, i.e., habituation effects across multiple days (group 3), effects of DarkLight phase across 24h (group 4), differences in the pattern of behavior in the few hours before and after phase shifts (group5), and differences in activity bout properties on the sub-minute time scale (group 6).

**Group 5: Activity pattern**
The activity of mice follows a complex circadian pattern, with prominent changes during periods surrounding light/dark phase transition. To capture strain-specific circadian patterns, we studied the behavior of mice by quantifying the anticipation and response to the onset of both light and dark phases.
change in anticipation of dark (fraction of total time) : Change in activity during the last 5 hours of the light phase in anticipation of the upcoming dark phase
change in anticipation of light (fraction of total time) : Change in activity during the last 5 hours of the dark phase in anticipation of the upcoming light phase
change in response to to dark (fraction of total time) : Change in activity during the first 2 hours of the dark phase in response to the onset of the dark phase
change in response to to light (fraction of total time) : Change in activity during the first 2 hours of the light phase in response to the onset of the light phase

Activity change in anticipation of dark

*Activity change in anticipation of dark*: Change in activity during the last 5 hours of the light phase in anticipation of the upcoming dark phase

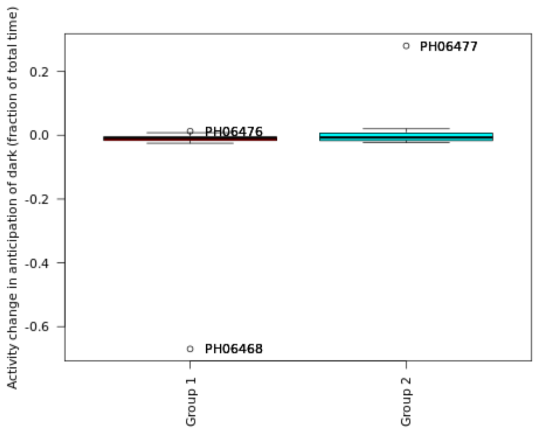

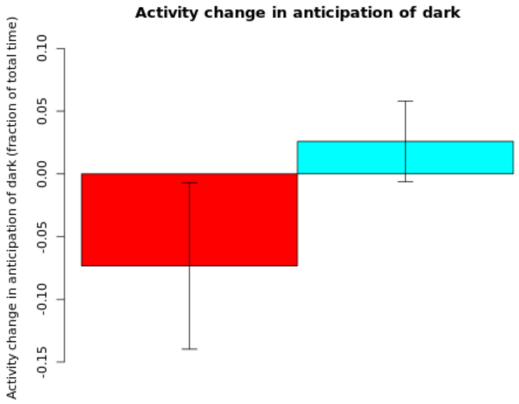

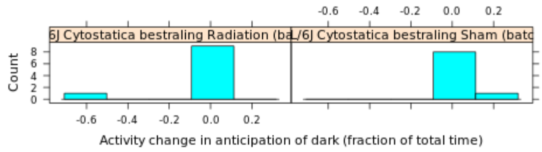

#### Secondary plot:

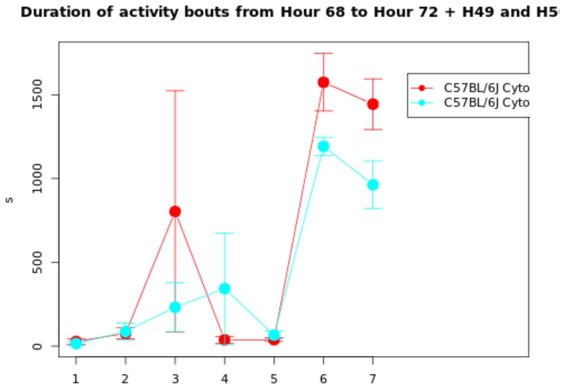

| Genotype | Mean | Median | SEM | N |
|---|---|---|---|---|
| (Gr 1): C57BL/6J Cytostatica bestraling Radiation (batch 22) | -0.0734 | -0.0091 | 0.0664 | 10 |
| (Gr 2): C57BL/6J Cytostatica bestraling Sham (batch 22) | 0.0259 | -0.007 | 0.0321 | 9 |

| | |
|---|---|
| **T-test:** | -1.3456 |
| **P-value:** | 0.2016 |
| **Df:** | 12.914 |
| **Mann-Whitney U test:** | 0.6038 |

| Mice excluded by Quality Control |
|---|
| PH06469 (C57BL/6J - ) |

*© Sylics BV*

Activity change in anticipation of light

*Activity change in anticipation of light*: Change in activity during the last 5 hours of the dark phase in anticipation of the upcoming light phase

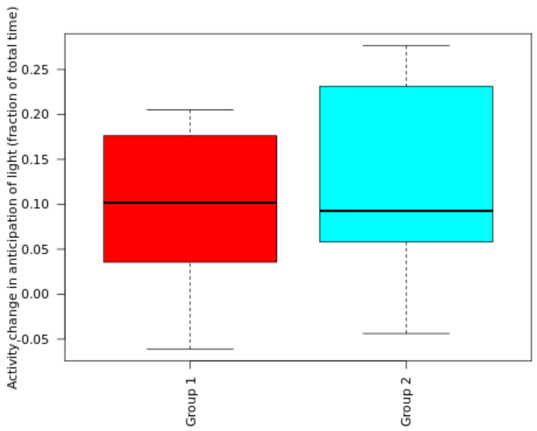

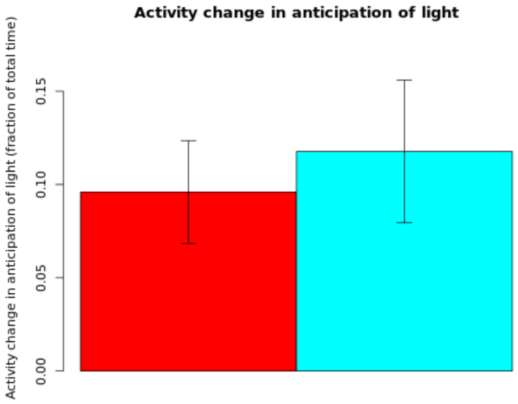

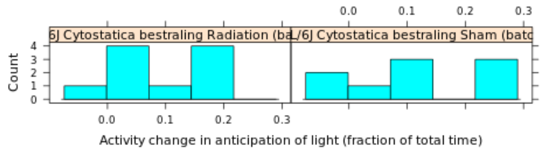

#### Secondary plot:

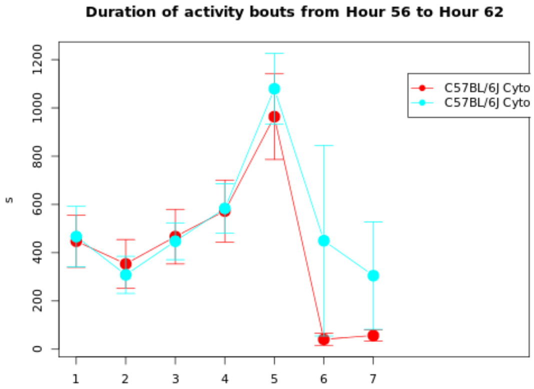

| Genotype | Mean | Median | SEM | N |
|---|---|---|---|---|
| (Gr 1): C57BL/6J Cytostatica bestraling Radiation (batch 22) | 0.096 | 0.102 | 0.0276 | 10 |
| (Gr 2): C57BL/6J Cytostatica bestraling Sham (batch 22) | 0.1177 | 0.0927 | 0.0383 | 9 |

| | |
|---|---|
| **T-test:** | -0.4613 |
| **P-value:** | 0.6512 |
| **Df:** | 14.8963 |
| **Mann-Whitney U test:** | 0.6607 |

| Mice excluded by Quality Control |
|---|
| PH06469 (C57BL/6J - ) |

 © Sylics BV

Activity change in response to to dark

*Activity change in response to to dark*: Change in activity during the first 2 hours of the dark phase in response to the onset of the dark phase

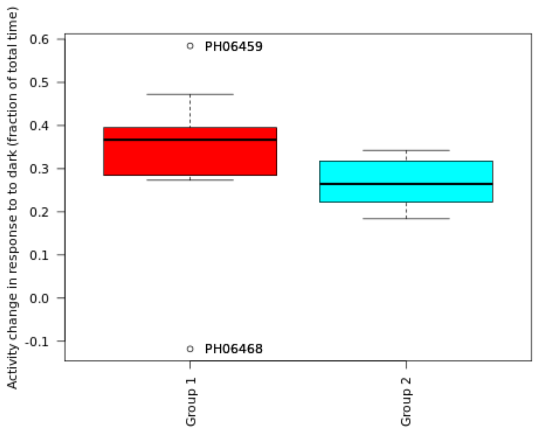

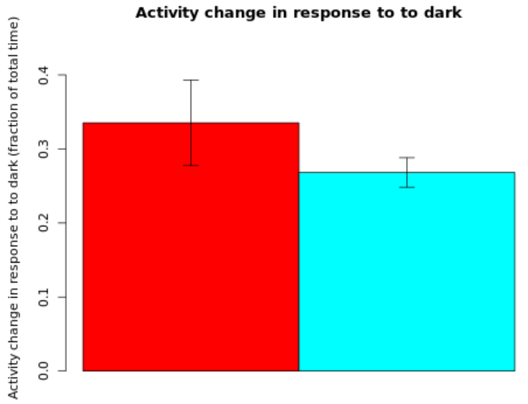

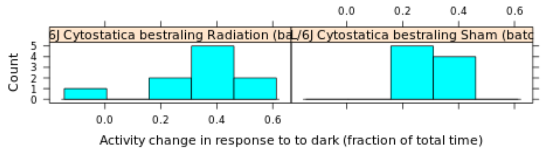

#### Secondary plot:

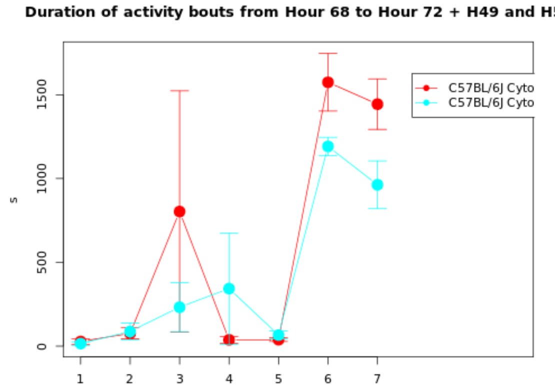

| Genotype | Mean | Median | SEM | N |
|---|---|---|---|---|
| (Gr 1): C57BL/6J Cytostatica bestraling Radiation (batch 22) | 0.3352 | 0.367 | 0.0578 | 10 |
| (Gr 2): C57BL/6J Cytostatica bestraling Sham (batch 22) | 0.2682 | 0.2646 | 0.0199 | 9 |

| | |
|---|---|
| **T-test:** | 1.0958 |
| **P-value:** | 0.2964 |
| **Df:** | 11.0886 |
| **Mann-Whitney U test:** | 0.022 |

| Mice excluded by Quality Control |
|---|
| PH06469 (C57BL/6J - ) |

 © Sylics BV

Activity change in response to to light

*Activity change in response to to light*: Change in activity during the first 2 hours of the light phase in response to the onset of the light phase

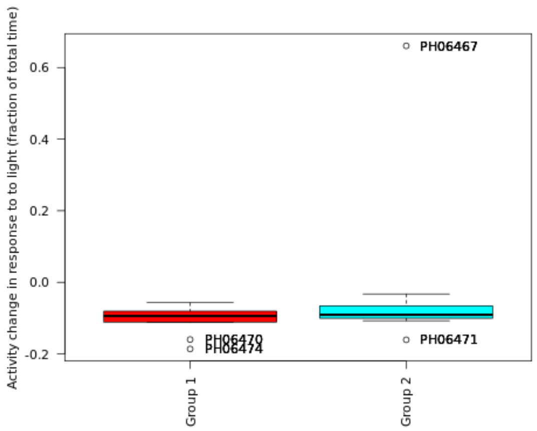

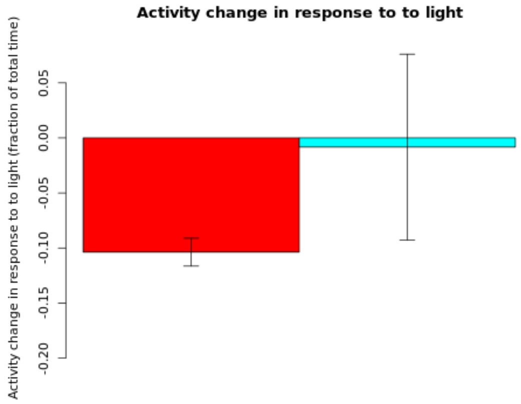

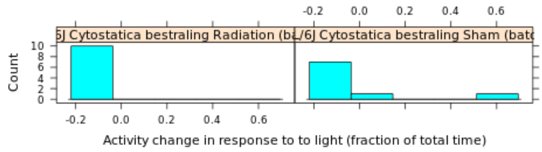

**Secondary plot:**

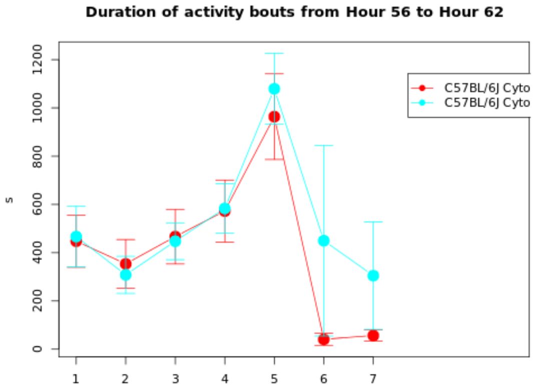

| Genotype | Mean | Median | SEM | N |
|---|---|---|---|---|
| (Gr 1): C57BL/6J Cytostatica bestraling Radiation (batch 22) | -0.1038 | -0.094 | 0.0126 | 10 |
| (Gr 2): C57BL/6J Cytostatica bestraling Sham (batch 22) | -0.0084 | -0.0906 | 0.0844 | 9 |

| | |
|---|---|
| **T-test:** | -1.1176 |
| **P-value:** | 0.2948 |
| **Df:** | 8.3593 |
| **Mann-Whitney U test:** | 0.549 |

| Mice excluded by Quality Control |
|---|
| PH06469 (C57BL/6J - ) |

 *© Sylics BV*

Feeding zone change in anticipation dark

*Feeding zone change in anticipation dark*: Change in Feeding zone during the last 5 hours of the light phase in anticipation of the upcoming dark phase

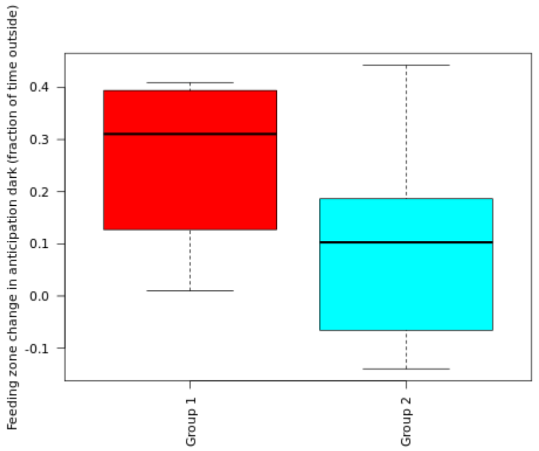

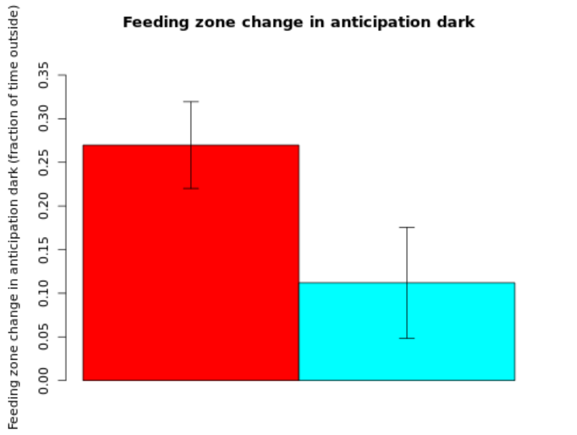

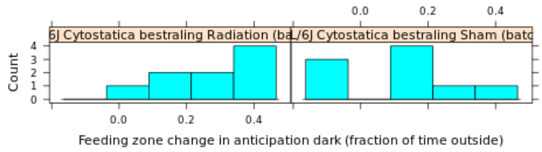

**Secondary plot:**

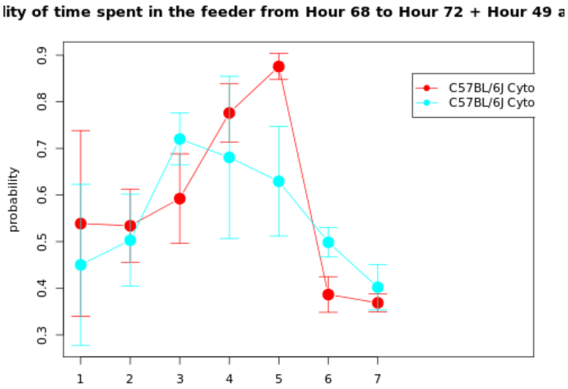

| Genotype | Mean | Median | SEM | N |
|---|---|---|---|---|
| (Gr 1): C57BL/6J Cytostatica bestraling Radiation (batch 22) | 0.2698 | 0.3106 | 0.0496 | 9 |
| (Gr 2): C57BL/6J Cytostatica bestraling Sham (batch 22) | 0.112 | 0.1029 | 0.0636 | 9 |

| | |
|---|---|
| **T-test:** | 1.9551 |
| **P-value:** | 0.0693 |
| **Df:** | 15.1082 |
| **Mann-Whitney U test:** | 0.0939 |

| Mice excluded by Quality Control |
|---|
| PH06469 (C57BL/6J - ) |

 © Sylics BV

Feeding zone change in response to dark

*Feeding zone change in response to dark*: Change in Feeding zone during the last 5 hours of the dark phase in anticipation of the upcoming light phase

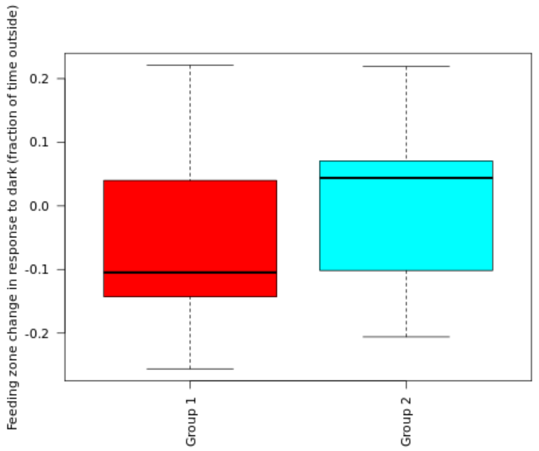

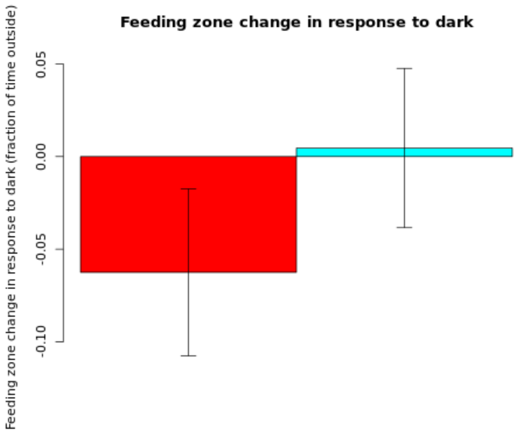

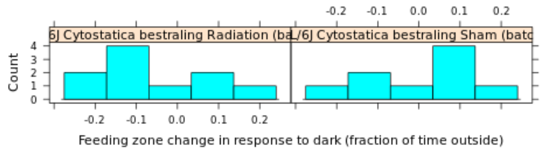

**Secondary plot:**

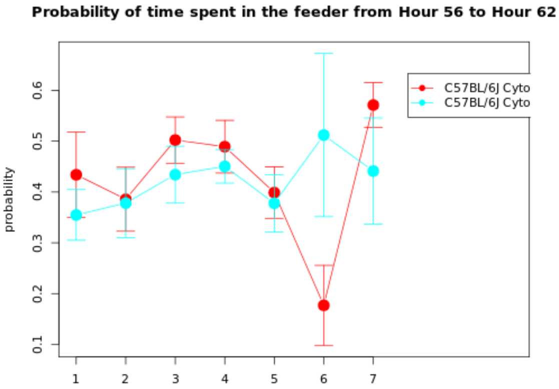

| Genotype | Mean | Median | SEM | N |
|---|---|---|---|---|
| (Gr 1): C57BL/6J Cytostatica bestraling Radiation (batch 22) | -0.0625 | -0.1047 | 0.045 | 10 |
| (Gr 2): C57BL/6J Cytostatica bestraling Sham (batch 22) | 0.0046 | 0.0439 | 0.043 | 9 |

| | |
|---|---|
| **T-test:** | -1.0784 |
| **P-value:** | 0.2959 |
| **Df:** | 16.9968 |
| **Mann-Whitney U test:** | 0.2428 |

| Mice excluded by Quality Control |
|---|
| PH06469 (C57BL/6J - ) |

 *© Sylics BV*

Feeding zone change in anticipation light

*Feeding zone change in anticipation light*: Change in Feeding zone during the first 2 hours of the dark phase in response to the onset of the dark phase

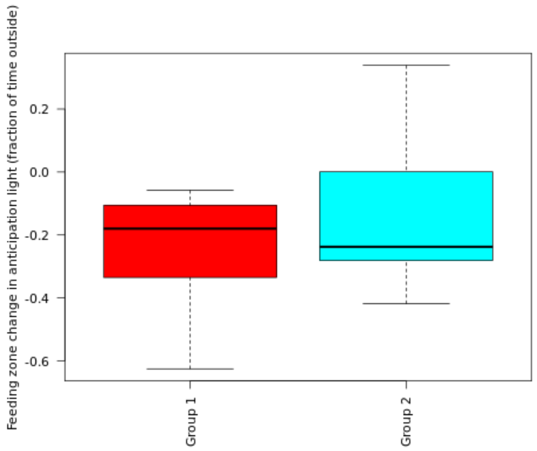

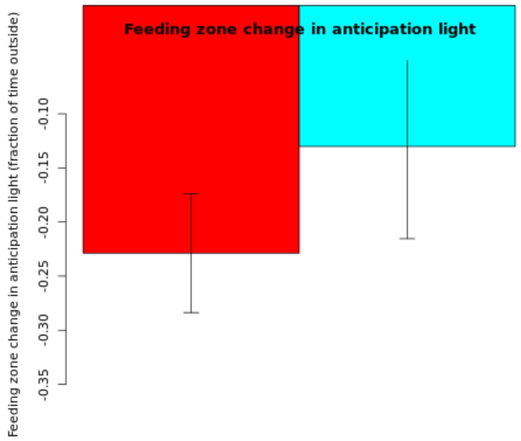

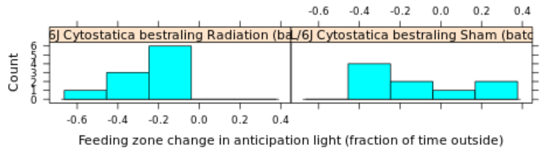

**Secondary plot:**

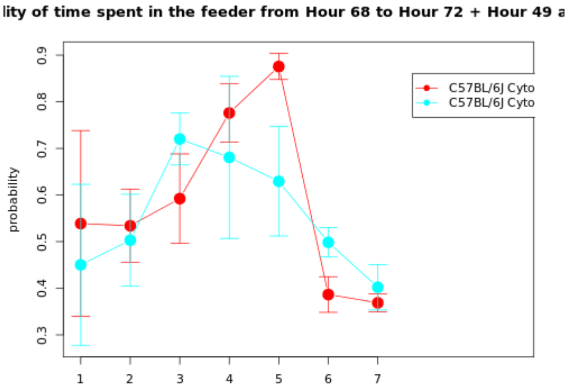

| Genotype | Mean | Median | SEM | N |
|---|---|---|---|---|
| (Gr 1): C57BL/6J Cytostatica bestraling Radiation (batch 22) | -0.2289 | -0.1798 | 0.0548 | 10 |
| (Gr 2): C57BL/6J Cytostatica bestraling Sham (batch 22) | -0.1302 | -0.2376 | 0.0852 | 9 |

| | |
|---|---|
| **T-test:** | -0.9738 |
| **P-value:** | 0.3468 |
| **Df:** | 13.8808 |
| **Mann-Whitney U test:** | 0.7802 |

| Mice excluded by Quality Control |
|---|
| PH06469 (C57BL/6J - ) |

 *© Sylics BV*

Feeding zone change in response to light

*Feeding zone change in response to light*: Change in Feeding zone during the first 2 hours of the light phase in response to the onset of the light phase

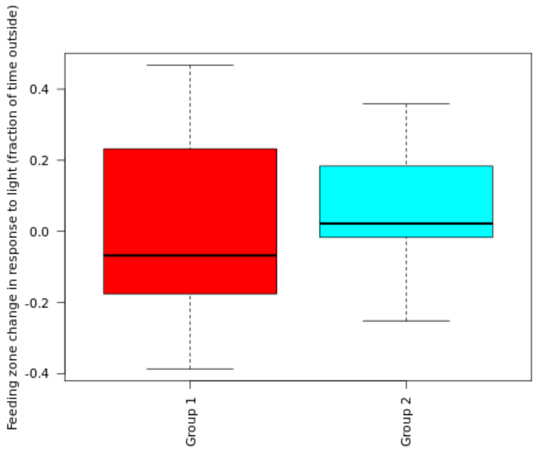

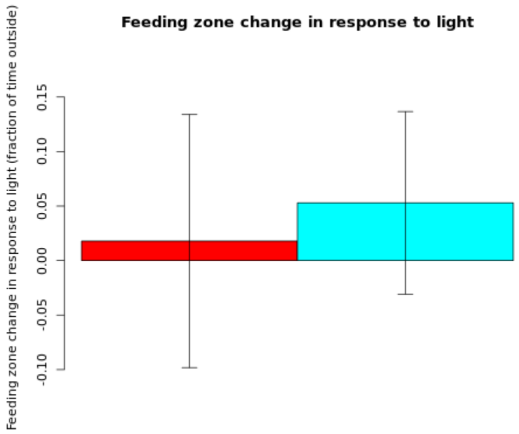

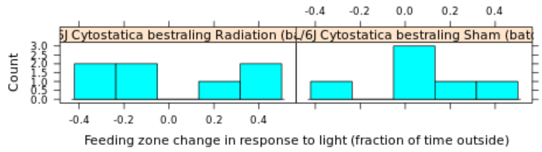

**Secondary plot:**

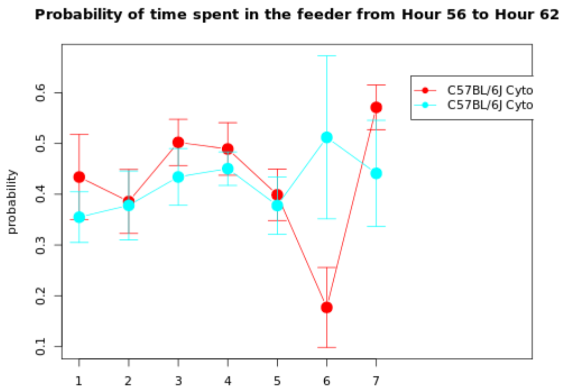

| Genotype | Mean | Median | SEM | N |
|---|---|---|---|---|
| (Gr 1): C57BL/6J Cytostatica bestraling Radiation (batch 22) | 0.0179 | -0.0678 | 0.1161 | 7 |
| (Gr 2): C57BL/6J Cytostatica bestraling Sham (batch 22) | 0.053 | 0.0221 | 0.0837 | 6 |

| | |
|---|---|
| **T-test:** | -0.2453 |
| **P-value:** | 0.811 |
| **Df:** | 10.4672 |
| **Mann-Whitney U test:** | 0.7308 |

| Mice excluded by Quality Control |
|---|
| PH06469 (C57BL/6J - ) |

OnShelter zone change in anticipation dark

*OnShelter zone change in anticipation dark*: Change in OnShelter zone during the last 5 hours of the light phase in anticipation of the upcoming dark phase

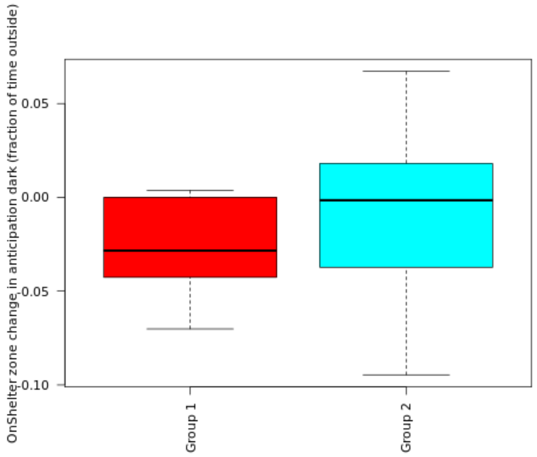

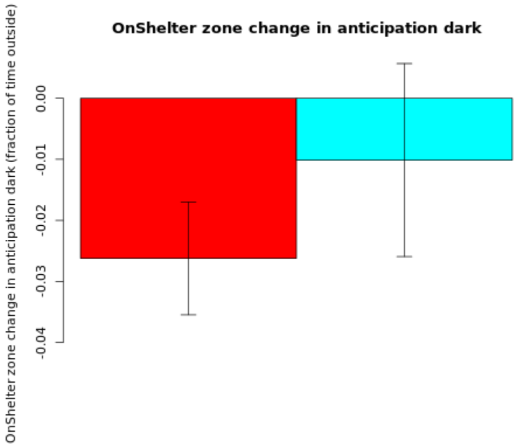

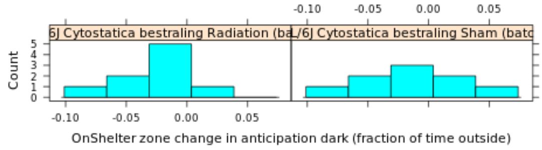

**Secondary plot:**

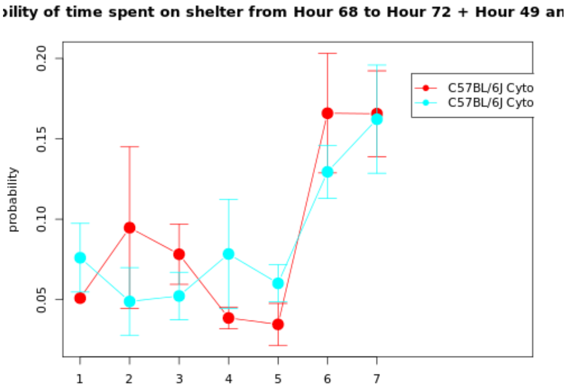

| Genotype | Mean | Median | SEM | N |
|---|---|---|---|---|
| (Gr 1): C57BL/6J Cytostatica bestraling Radiation (batch 22) | -0.0262 | -0.0284 | 0.0092 | 9 |
| (Gr 2): C57BL/6J Cytostatica bestraling Sham (batch 22) | -0.0101 | -0.0015 | 0.0158 | 9 |

| | |
|---|---|
| **T-test:** | -0.8797 |
| **P-value:** | 0.3951 |
| **Df:** | 12.8994 |
| **Mann-Whitney U test:** | 0.4258 |

| Mice excluded by Quality Control |
|---|
| PH06469 (C57BL/6J - ) |

 *© Sylics BV*

Sylics

OnShelter zone change in response to dark

*OnShelter zone change in response to dark*: Change in OnShelter zone during the last 5 hours of the dark phase in anticipation of the upcoming light phase

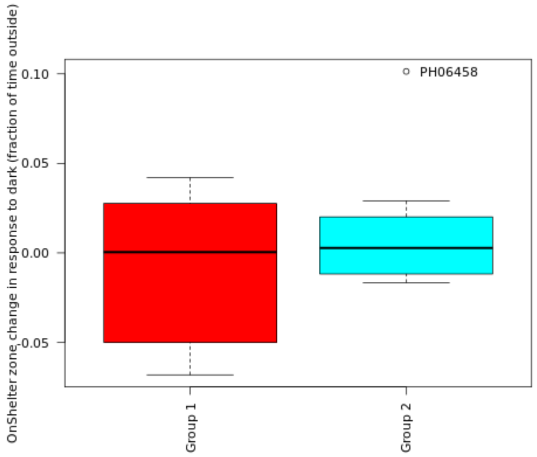

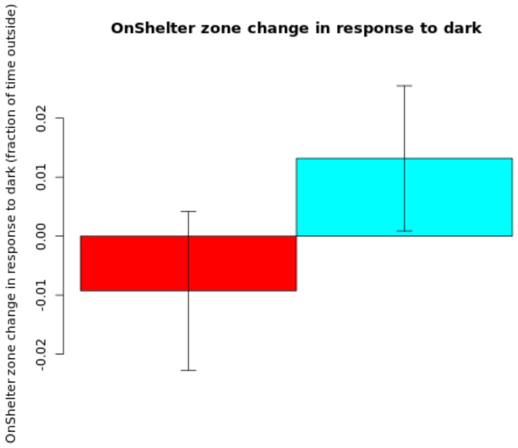

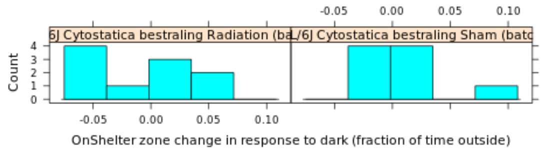

**Secondary plot:**

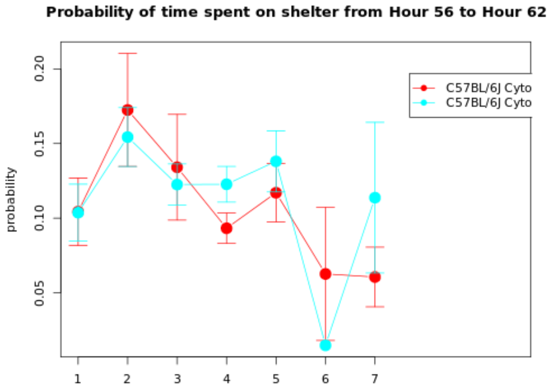

| Genotype | Mean | Median | SEM | N |
|---|---|---|---|---|
| (Gr 1): C57BL/6J Cytostatica bestraling Radiation (batch 22) | -0.0093 | 0.0004 | 0.0135 | 10 |
| (Gr 2): C57BL/6J Cytostatica bestraling Sham (batch 22) | 0.0132 | 0.0028 | 0.0123 | 9 |

| | |
|---|---|
| **T-test:** | -1.2299 |
| **P-value:** | 0.2355 |
| **Df:** | 16.9822 |
| **Mann-Whitney U test:** | 0.549 |

| Mice excluded by Quality Control |
|---|
| PH06469 (C57BL/6J - ) |

 © Sylics BV

OnShelter zone change in anticipation light

*OnShelter zone change in anticipation light*: Change in OnShelter zone during the first 2 hours of the dark phase in response to the onset of the dark phase

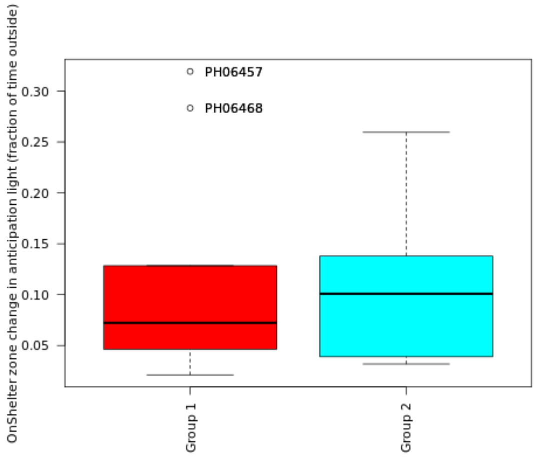

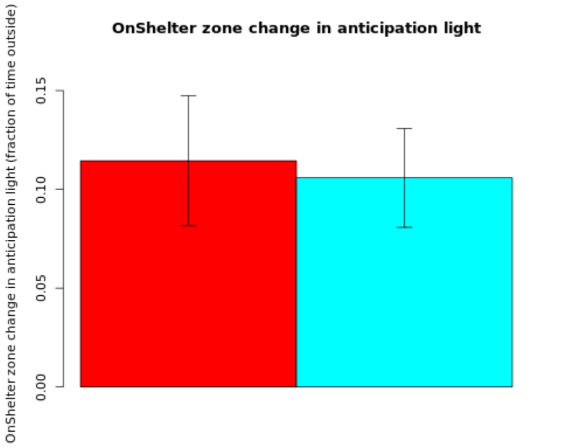

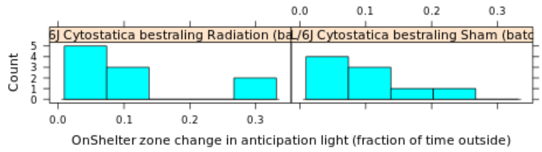

**Secondary plot:**

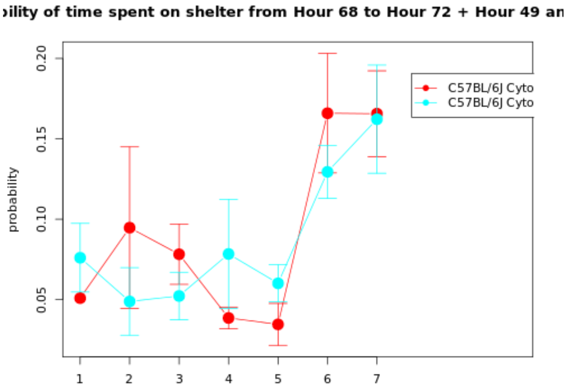

| Genotype | Mean | Median | SEM | N |
|---|---|---|---|---|
| (Gr 1): C57BL/6J Cytostatica bestraling Radiation (batch 22) | 0.1145 | 0.0723 | 0.033 | 10 |
| (Gr 2): C57BL/6J Cytostatica bestraling Sham (batch 22) | 0.1059 | 0.1009 | 0.0251 | 9 |

| | |
|---|---|
| **T-test:** | 0.2081 |
| **P-value:** | 0.8378 |
| **Df:** | 16.2889 |
| **Mann-Whitney U test:** | 0.9682 |

| Mice excluded by Quality Control |
|---|
| PH06469 (C57BL/6J - ) |

 *© Sylics BV*

OnShelter zone change in response to light

*OnShelter zone change in response to light*: Change in OnShelter zone during the first 2 hours of the light phase in response to the onset of the light phase

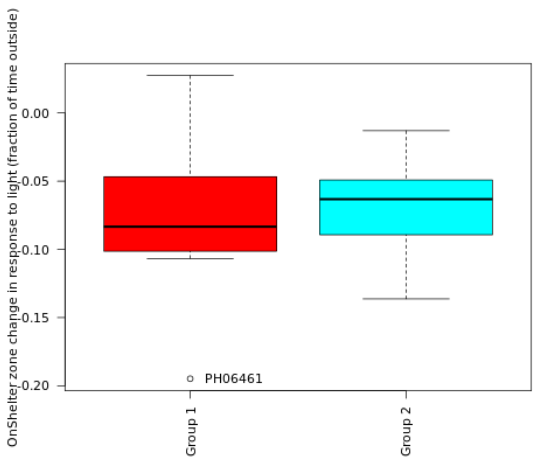

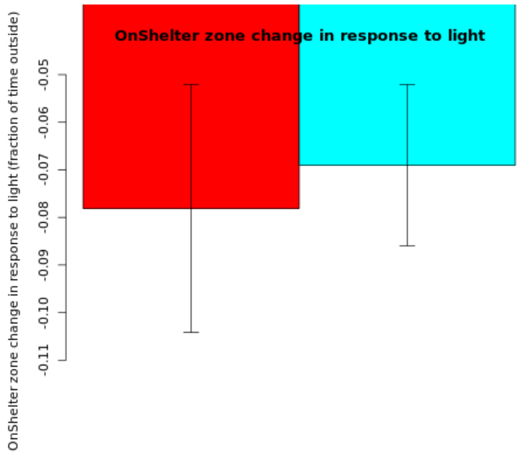

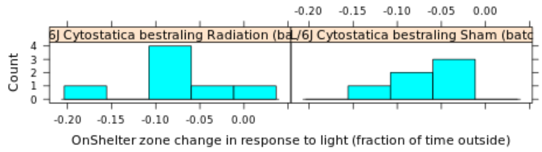

**Secondary plot:**

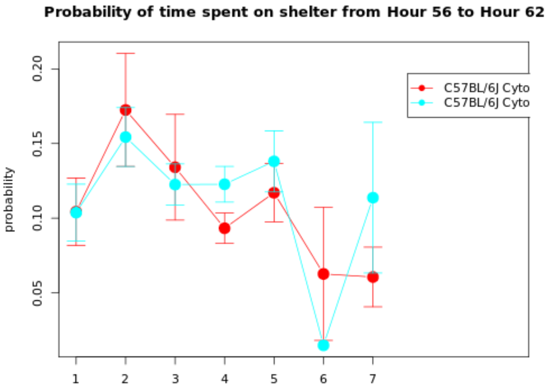

| Genotype | Mean | Median | SEM | N |
|---|---|---|---|---|
| (Gr 1): C57BL/6J Cytostatica bestraling Radiation (batch 22) | -0.0781 | -0.0834 | 0.026 | 7 |
| (Gr 2): C57BL/6J Cytostatica bestraling Sham (batch 22) | -0.069 | -0.0632 | 0.0169 | 6 |

| | |
|---|---|
| **T-test:** | -0.2932 |
| **P-value:** | 0.7754 |
| **Df:** | 9.9918 |
| **Mann-Whitney U test:** | 0.7308 |

| Mice excluded by Quality Control |
|---|
| PH06469 (C57BL/6J - ) |

 © Sylics BV

Sylics

Spout zone change in anticipation dark

*Spout zone change in anticipation dark*: Change in Spout zone during the last 5 hours of the light phase in anticipation of the upcoming dark phase

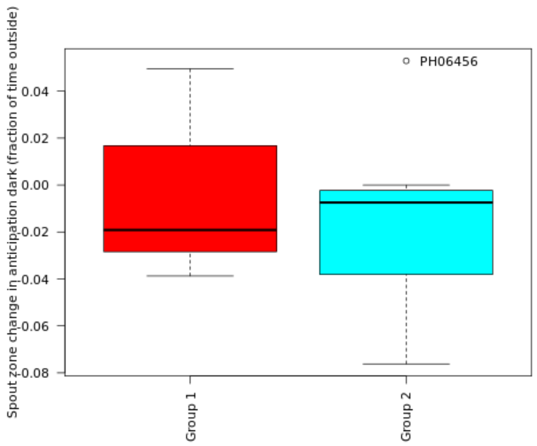

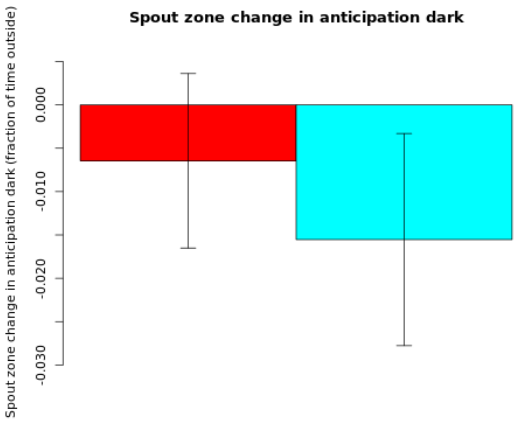

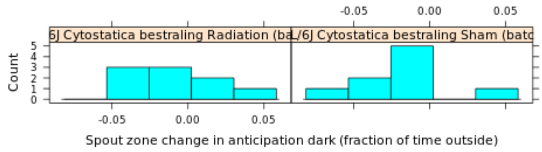

#### Secondary plot:

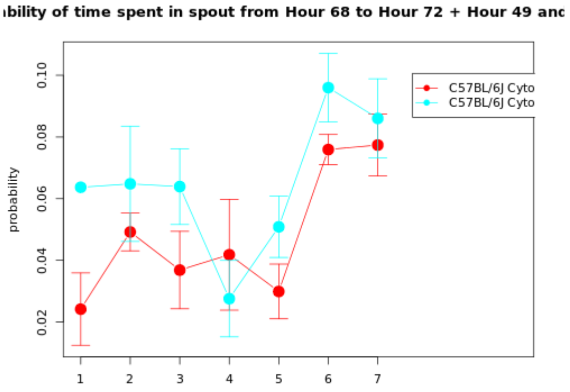

| Genotype | Mean | Median | SEM | N |
|---|---|---|---|---|
| (Gr 1): C57BL/6J Cytostatica bestraling Radiation (batch 22) | -0.0065 | -0.0191 | 0.0101 | 9 |
| (Gr 2): C57BL/6J Cytostatica bestraling Sham (batch 22) | -0.0155 | -0.0074 | 0.0122 | 9 |

| | |
|---|---|
| **T-test:** | 0.5736 |
| **P-value:** | 0.5745 |
| **Df:** | 15.4228 |
| **Mann-Whitney U test:** | 0.6665 |

| Mice excluded by Quality Control |
|---|
| PH06469 (C57BL/6J - ) |

 © Sylics BV

Spout zone change in response to dark

*Spout zone change in response to dark*: Change in Spout zone during the last 5 hours of the dark phase in anticipation of the upcoming light phase

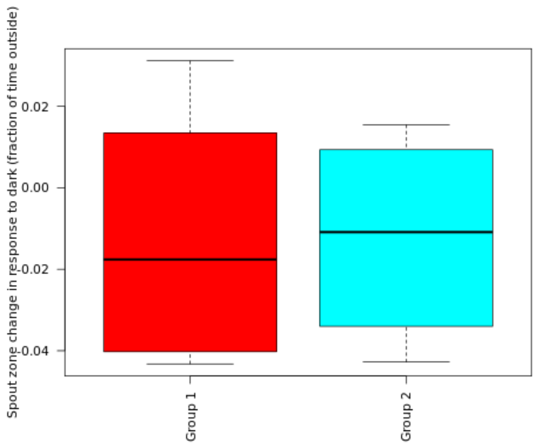

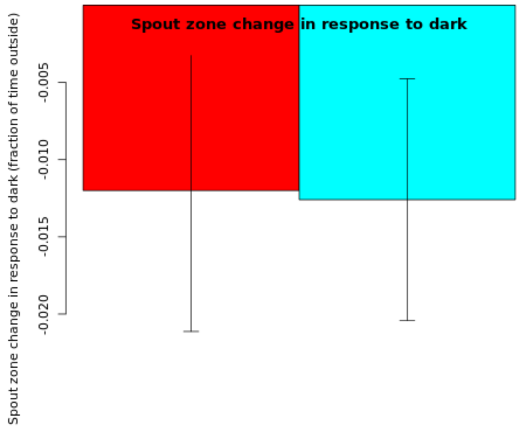

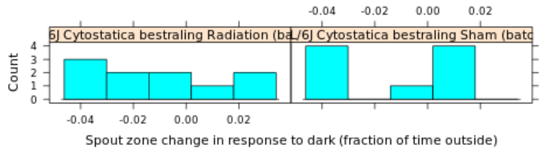

**Secondary plot:**

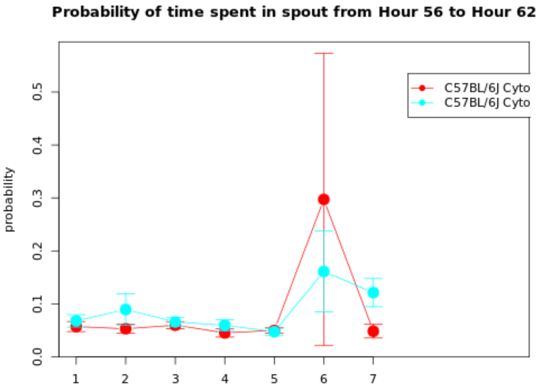

| Genotype | Mean | Median | SEM | N |
|---|---|---|---|---|
| (Gr 1): C57BL/6J Cytostatica bestraling Radiation (batch 22) | -0.012 | -0.0176 | 0.0091 | 10 |
| (Gr 2): C57BL/6J Cytostatica bestraling Sham (batch 22) | -0.0126 | -0.0109 | 0.0078 | 9 |

| | |
|---|---|
| **T-test:** | 0.0491 |
| **P-value:** | 0.9614 |
| **Df:** | 16.8545 |
| **Mann-Whitney U test:** | 0.9682 |

| Mice excluded by Quality Control |
|---|
| PH06469 (C57BL/6J - ) |

Spout zone change in anticipation light

*Spout zone change in anticipation light*: Change in Spout zone during the first 2 hours of the dark phase in response to the onset of the dark phase

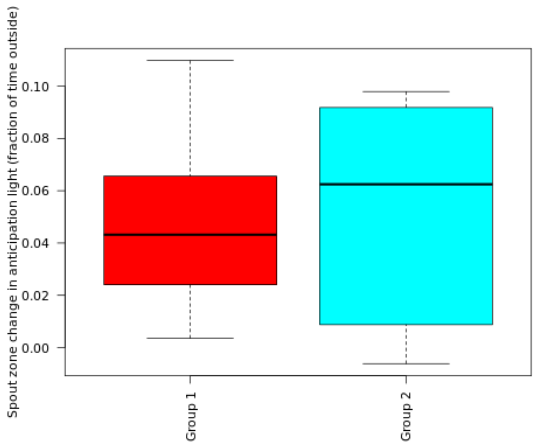

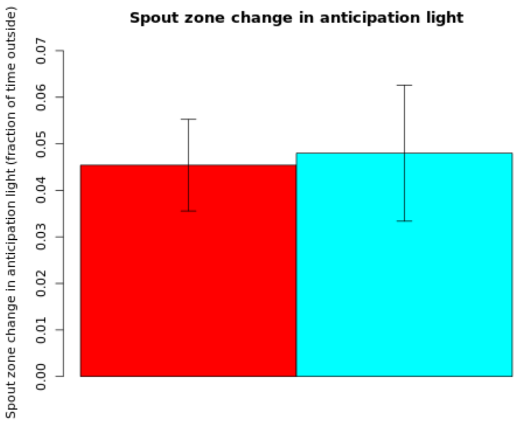

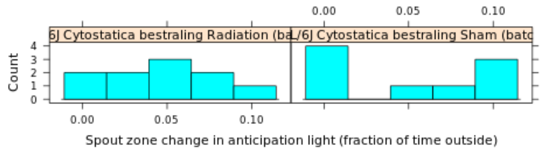

#### Secondary plot:

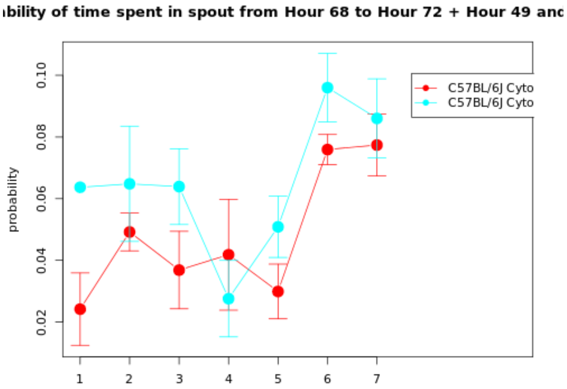

| Genotype | Mean | Median | SEM | N |
|---|---|---|---|---|
| (Gr 1): C57BL/6J Cytostatica bestraling Radiation (batch 22) | 0.0454 | 0.0432 | 0.0098 | 10 |
| (Gr 2): C57BL/6J Cytostatica bestraling Sham (batch 22) | 0.048 | 0.0625 | 0.0146 | 9 |

| | |
|---|---|
| **T-test:** | -0.1464 |
| **P-value:** | 0.8857 |
| **Df:** | 14.281 |
| **Mann-Whitney U test:** | 1 |

| Mice excluded by Quality Control |
|---|
| PH06469 (C57BL/6J - ) |

Spout zone change in response to light

*Spout zone change in response to light*: Change in Spout zone during the first 2 hours of the light phase in response to the onset of the light phase

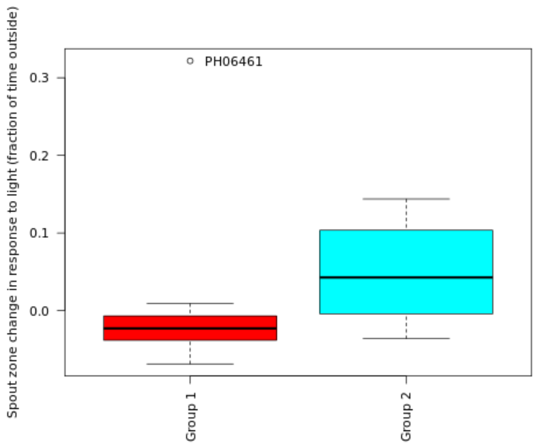

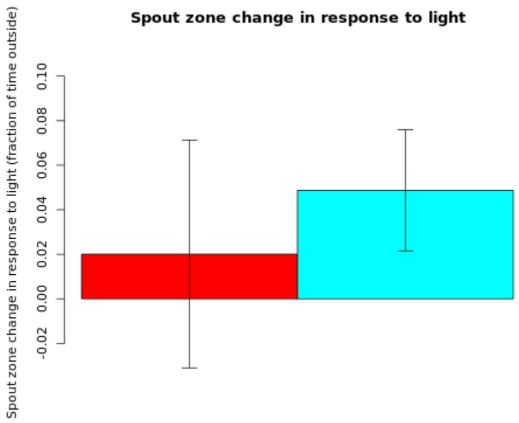

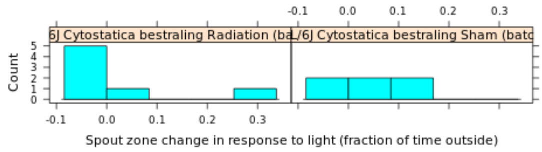

**Secondary plot:**

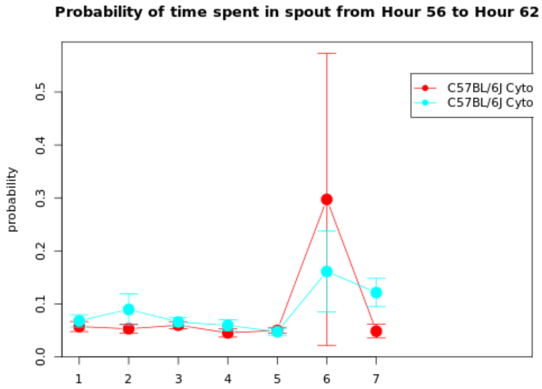

| Genotype | Mean | Median | SEM | N |
|---|---|---|---|---|
| (Gr 1): C57BL/6J Cytostatica bestraling Radiation (batch 22) | 0.0201 | -0.0229 | 0.0511 | 7 |
| (Gr 2): C57BL/6J Cytostatica bestraling Sham (batch 22) | 0.0488 | 0.0427 | 0.0272 | 6 |

| | |
|---|---|
| **T-test:** | -0.4957 |
| **P-value:** | 0.632 |
| **Df:** | 9.0177 |
| **Mann-Whitney U test:** | 0.1807 |

| Mice excluded by Quality Control |
|---|
| PH06469 (C57BL/6J - ) |

 *© Sylics BV*

### Spontaneous behavior (Group 6: activity)

With respect to spontaneous behaviors in the first three days in the PhenoTyper, 6 groups of behavioral parameters are defined as described below. The first two groups describe specific behavioral elements related to kinematics of mice (description of movement characteristics, group1) and sheltering behavior (group 2). These behavioral parameters were analyzed with respect to temporal aspects, in particular over 4 different time scales, i.e., habituation effects across multiple days (group 3), effects of DarkLight phase across 24h (group 4), differences in the pattern of behavior in the few hours before and after phase shifts (group5), and differences in activity bout properties on the sub-minute time scale (group 6).

**Group 6: activity**
Characteristics and quantity of individual activity bouts.

 *© Sylics BV*

Activity duration - dark

*Activity duration - dark*: Cummulative duration of activity during the dark phase

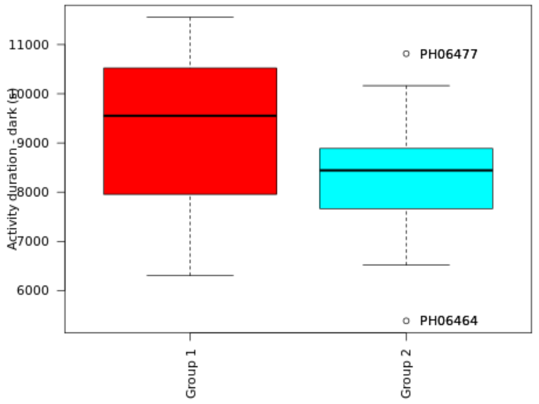

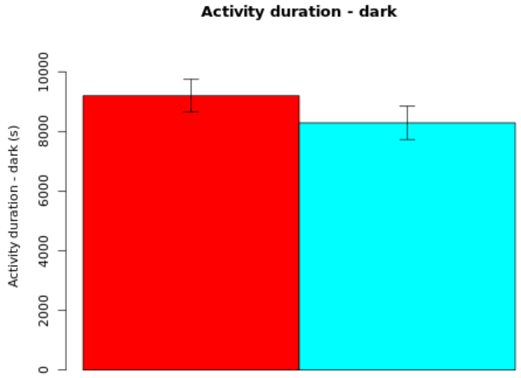

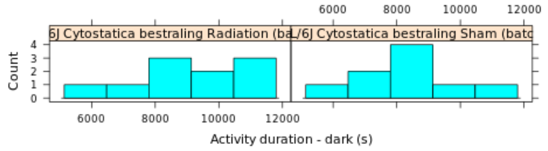

#### Secondary plot:

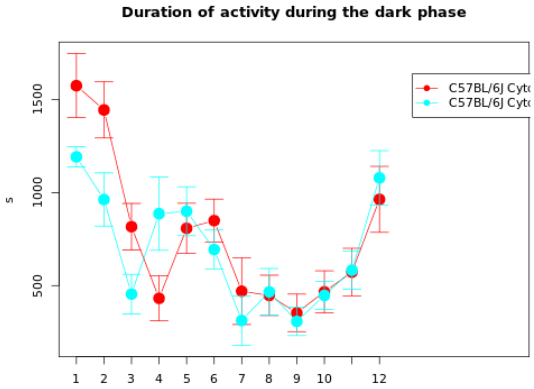

| Genotype | Mean | Median | SEM | N |
|---|---|---|---|---|
| (Gr 1): C57BL/6J Cytostatica bestraling Radiation (batch 22) | 9199.1907 | 9552.3133 | 546.3186 | 10 |
| (Gr 2): C57BL/6J Cytostatica bestraling Sham (batch 22) | 8287.8726 | 8443 | 558.3474 | 9 |

| | |
|---|---|
| **T-test:** | 1.1666 |
| **P-value:** | 0.2596 |
| **Df:** | 16.8899 |
| **Mann-Whitney U test:** | 0.3154 |

| Mice excluded by Quality Control |
|---|
| PH06469 (C57BL/6J - ) |

Mean activity duration - dark

*Mean activity duration - dark*: Mean duration per activity bout during the dark phase

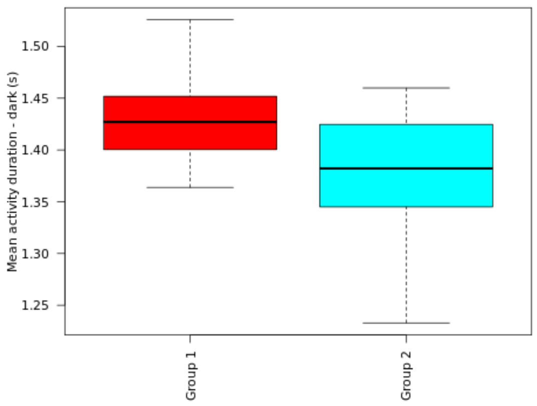

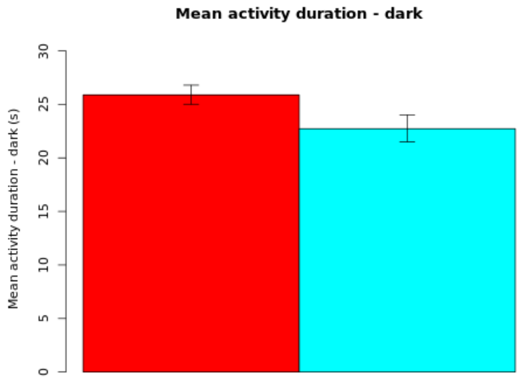

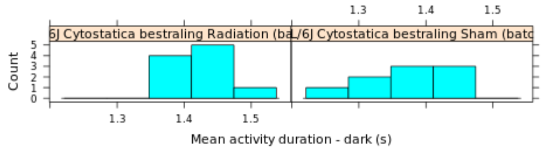

**NOTE: The statistics and boxplot are performed on log10 transformed data**

| Genotype | Mean | Median | SEM | N |
|---|---|---|---|---|
| (Gr 1): C57BL/6J Cytostatica bestraling Radiation (batch 22) | 25.8879 | 0 | Up: 0.9076<br>Down: 0.878 | 10 |
| (Gr 2): C57BL/6J Cytostatica bestraling Sham (batch 22) | 22.7262 | 0 | Up: 1.2879<br>Down: 1.2215 | 9 |

| | |
|---|---|
| **T-test:** | 2.0042 |
| **P-value:** | 0.0653 |
| **Df:** | 13.6663 |
| **Mann-Whitney U test:** | 0.0535 |

| Mice excluded by Quality Control |
|---|
| PH06469 (C57BL/6J - ) |

 *© Sylics BV*

Activity number - dark

*Activity number - dark*: Cummulative number of activity bouts during the dark phase

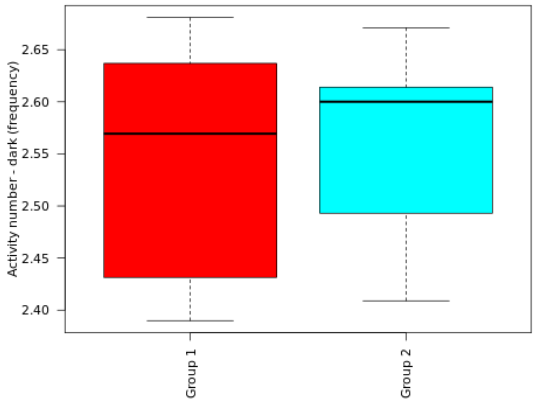

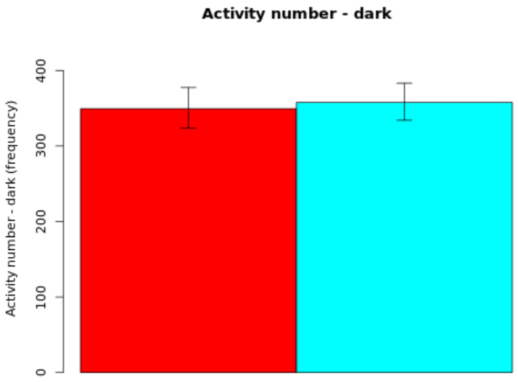

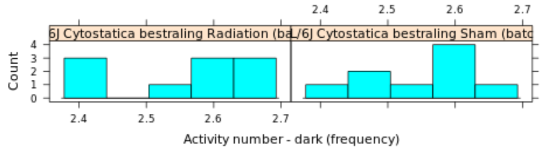

#### Secondary plot:

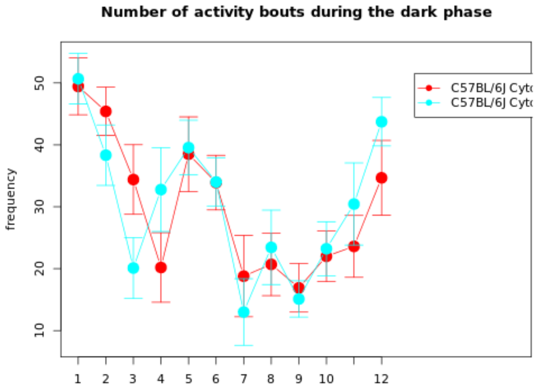

**NOTE: The statistics and boxplot are performed on log10 transformed data**

| Genotype | Mean | Median | SEM | N |
|---|---|---|---|---|
| (Gr 1): C57BL/6J Cytostatica bestraling Radiation (batch 22) | 349.4097 | 0 | Up: 28.0322<br>Down: 25.9558 | 10 |
| (Gr 2): C57BL/6J Cytostatica bestraling Sham (batch 22) | 357.8402 | 0 | Up: 25.4308<br>Down: 23.7478 | 9 |

| | |
|---|---|
| **T-test:** | -0.2308 |
| **P-value:** | 0.8202 |
| **Df:** | 16.9437 |
| **Mann-Whitney U test:** | 0.8421 |

| Mice excluded by Quality Control |
|---|
| PH06469 (C57BL/6J - ) |

 © Sylics BV

Feeding zone duration - dark

*Feeding zone duration - dark*: Cummulative duration in feeding zone during the dark phase

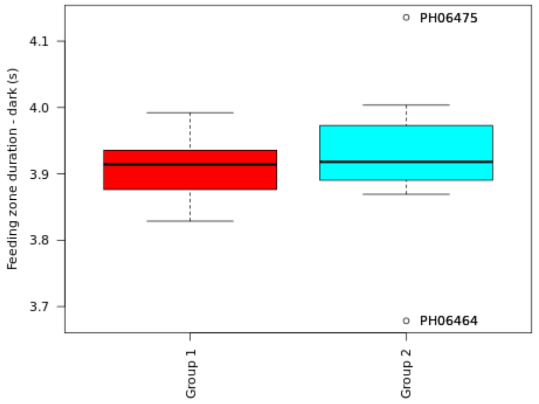

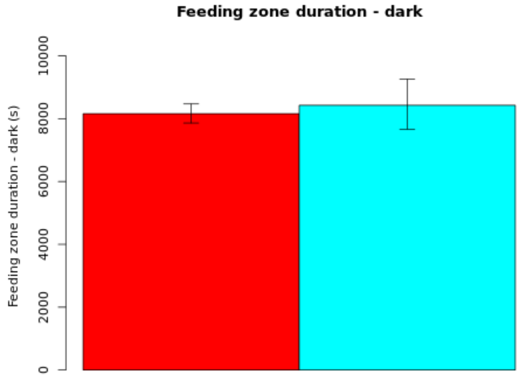

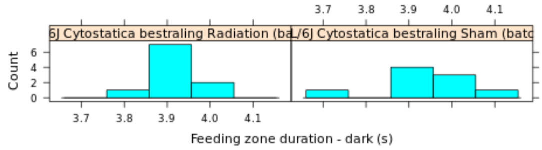

**Secondary plot:**

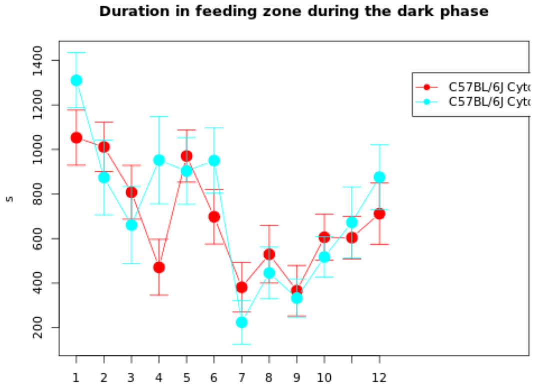

**NOTE: The statistics and boxplot are performed on log10 transformed data**

| Genotype | Mean | Median | SEM | N |
|---|---|---|---|---|
| (Gr 1): C57BL/6J Cytostatica bestraling Radiation (batch 22) | 8159.3751 | 0 | Up: 308.4847<br>Down: 297.2479 | 10 |
| (Gr 2): C57BL/6J Cytostatica bestraling Sham (batch 22) | 8424.8164 | 0 | Up: 835.2005<br>Down: 759.8783 | 9 |

| | |
|---|---|
| **T-test:** | -0.3153 |
| **P-value:** | 0.7588 |
| **Df:** | 10.4358 |
| **Mann-Whitney U test:** | 0.6038 |

| Mice excluded by Quality Control |
|---|
| PH06469 (C57BL/6J - ) |

© Sylics BV

Sylics

OnShelter zone duration - dark

*OnShelter zone duration - dark*: Cummulative duration in OnShelter zone during the dark phase

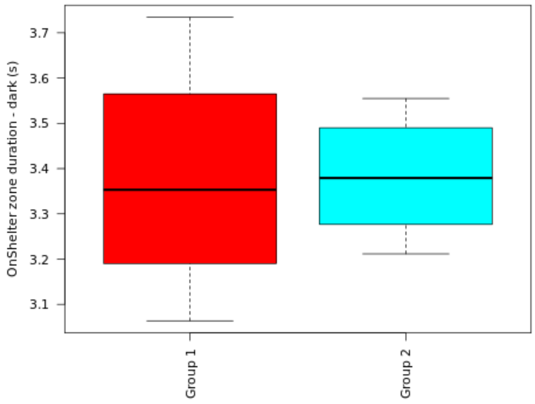

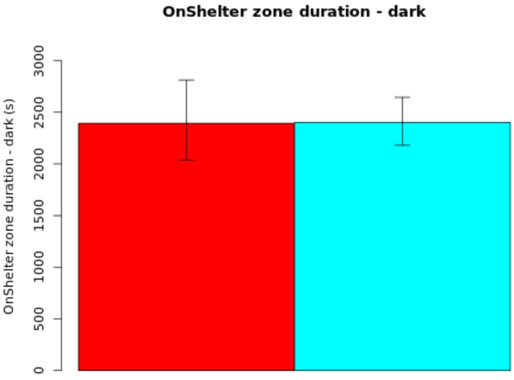

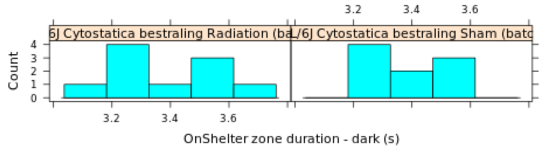

#### Secondary plot:

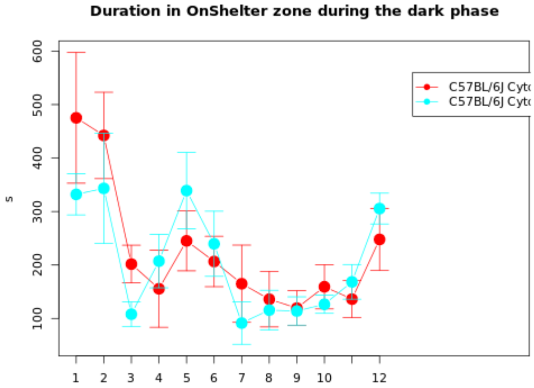

**NOTE: The statistics and boxplot are performed on log10 transformed data**

| Genotype | Mean | Median | SEM | N |
|---|---|---|---|---|
| (Gr 1): C57BL/6J Cytostatica bestraling Radiation (batch 22) | 2393.1509 | 0 | Up: 419.1457<br>Down: 356.6983 | 10 |
| (Gr 2): C57BL/6J Cytostatica bestraling Sham (batch 22) | 2400.9962 | 0 | Up: 242.8496<br>Down: 220.5512 | 9 |

| | |
|---|---|
| **T-test:** | -0.0174 |
| **P-value:** | 0.9863 |
| **Df:** | 14.4881 |
| **Mann-Whitney U test:** | 0.9682 |

| Mice excluded by Quality Control |
|---|
| PH06469 (C57BL/6J - ) |

 © Sylics BV

OnShelter zone number - dark

*OnShelter zone number - dark*: Cummulative number of visits to OnShelter zone during the dark phase

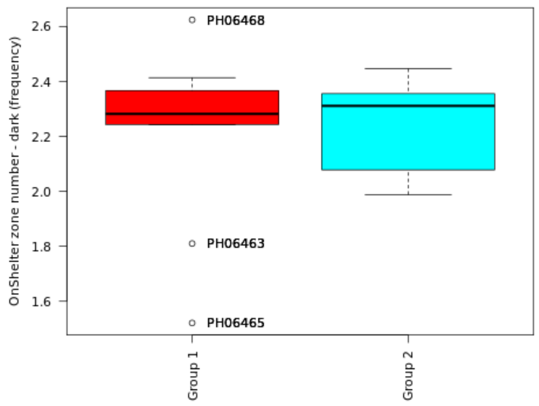

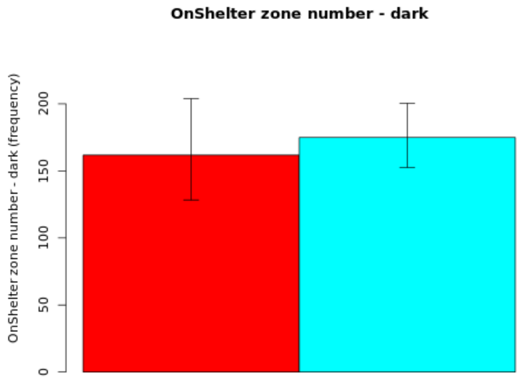

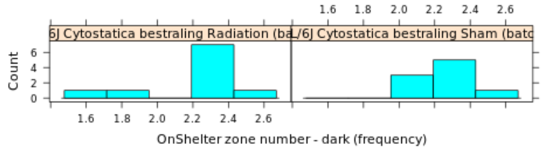

**Secondary plot:**

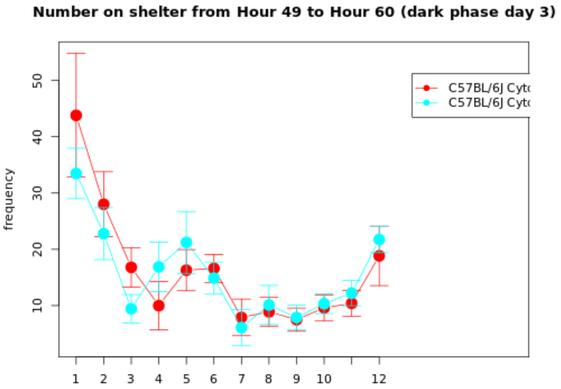

**NOTE: The statistics and boxplot are performed on log10 transformed data**

| Genotype | Mean | Median | SEM | N |
|---|---|---|---|---|
| (Gr 1): C57BL/6J Cytostatica bestraling Radiation (batch 22) | 161.7885 | 0 | Up: 42.0686<br>Down: 33.4296 | 10 |
| (Gr 2): C57BL/6J Cytostatica bestraling Sham (batch 22) | 174.9288 | 0 | Up: 25.4856<br>Down: 22.2608 | 9 |

| | |
|---|---|
| **T-test:** | -0.291 |
| **P-value:** | 0.7752 |
| **Df:** | 14.3746 |
| **Mann-Whitney U test:** | 0.7197 |

| Mice excluded by Quality Control |
|---|
| PH06469 (C57BL/6J - ) |

Sylics

Spout zone duration - dark

*Spout zone duration - dark*: Cummulative duration in Spout zone during the dark phase

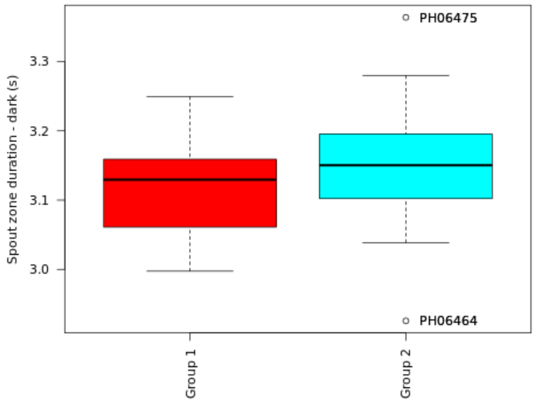

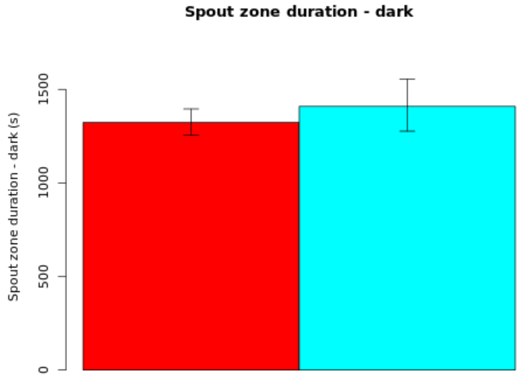

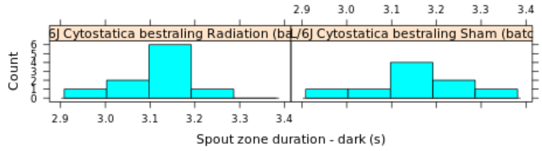

#### Secondary plot:

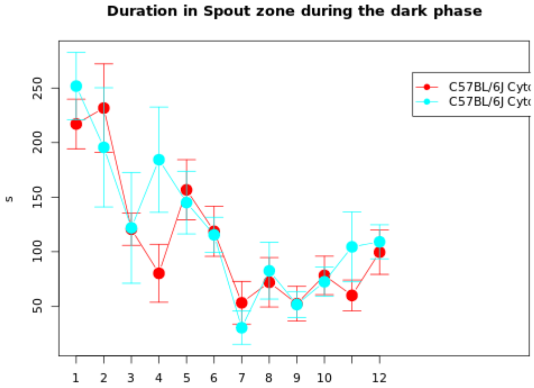

**NOTE: The statistics and boxplot are performed on log10 transformed data**

| Genotype | Mean | Median | SEM | N |
|---|---|---|---|---|
| (Gr 1): C57BL/6J Cytostatica bestraling Radiation (batch 22) | 1323.6769 | 0 | Up: 71.8074<br>Down: 68.1151 | 10 |
| (Gr 2): C57BL/6J Cytostatica bestraling Sham (batch 22) | 1408.9966 | 0 | Up: 146.0111<br>Down: 132.3099 | 9 |

| | |
|---|---|
| **T-test:** | -0.5584 |
| **P-value:** | 0.5866 |
| **Df:** | 12.3471 |
| **Mann-Whitney U test:** | 0.7197 |

| Mice excluded by Quality Control |
|---|
| PH06469 (C57BL/6J - ) |

© Sylics BV

Activity duration - light

*Activity duration - light*: Cummulative duration of activity during the light phase

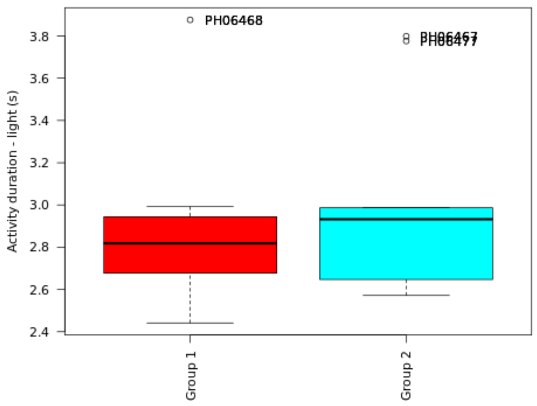

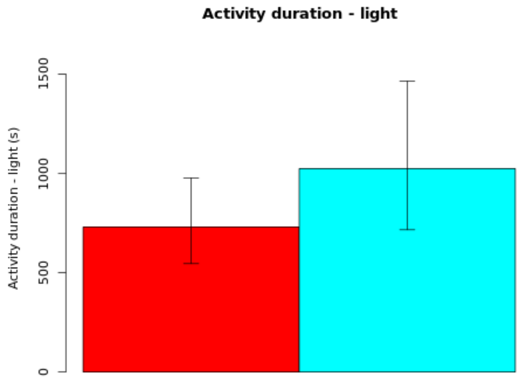

**Secondary plot:**

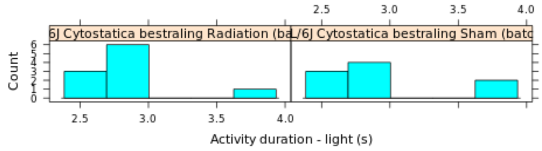

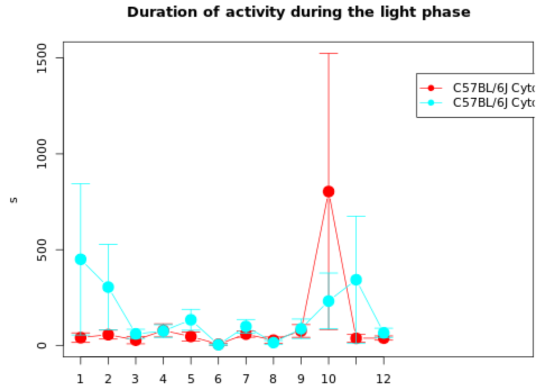

**NOTE: The statistics and boxplot are performed on log10 transformed data**

| Genotype | Mean | Median | SEM | N |
|---|---|---|---|---|
| (Gr 1): C57BL/6J Cytostatica bestraling Radiation (batch 22) | 730.1952 | 0 | Up: 246.9801<br>Down: 184.62 | 10 |
| (Gr 2): C57BL/6J Cytostatica bestraling Sham (batch 22) | 1024.0687 | 0 | Up: 440.3872<br>Down: 308.0455 | 9 |

| | |
|---|---|
| **T-test:** | -0.733 |
| **P-value:** | 0.4742 |
| **Df:** | 15.9084 |
| **Mann-Whitney U test:** | 0.6038 |

| Mice excluded by Quality Control |
|---|
| PH06469 (C57BL/6J - ) |

 © Sylics BV

Mean activity duration - light

*Mean activity duration - light*: Mean duration per activity bout during the light phase

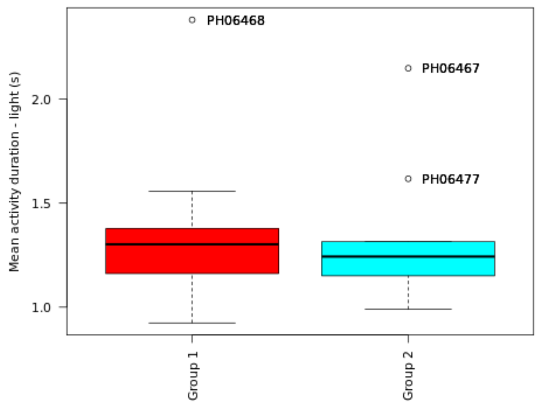

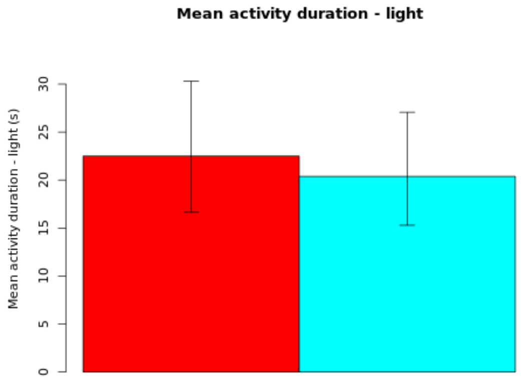

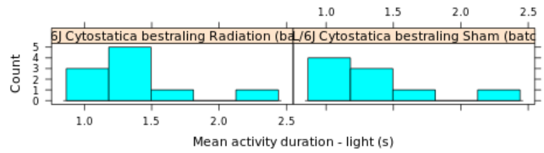

**NOTE: The statistics and boxplot are performed on log10 transformed data**

| Genotype | Mean | Median | SEM | N |
|---|---|---|---|---|
| (Gr 1): C57BL/6J Cytostatica bestraling Radiation (batch 22) | 22.5087 | 0 | Up: 7.8045<br>Down: 5.8593 | 10 |
| (Gr 2): C57BL/6J Cytostatica bestraling Sham (batch 22) | 20.382 | 0 | Up: 6.6832<br>Down: 5.0917 | 9 |

| | |
|---|---|
| **T-test:** | 0.24 |
| **P-value:** | 0.8132 |
| **Df:** | 16.9993 |
| **Mann-Whitney U test:** | 0.549 |

| Mice excluded by Quality Control |
|---|
| PH06469 (C57BL/6J - ) |

Activity number - light

*Activity number - light*: Cummulative number of activity bouts during the light phase

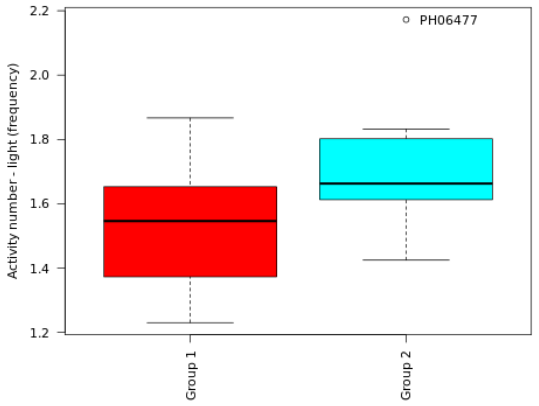

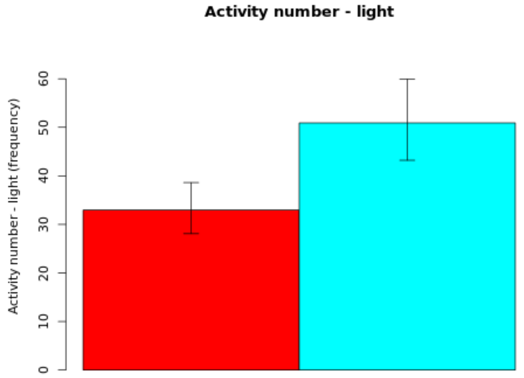

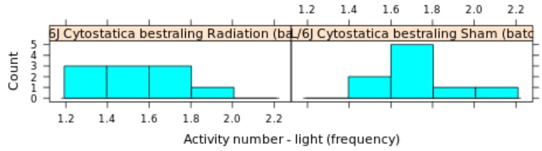

**Secondary plot:**

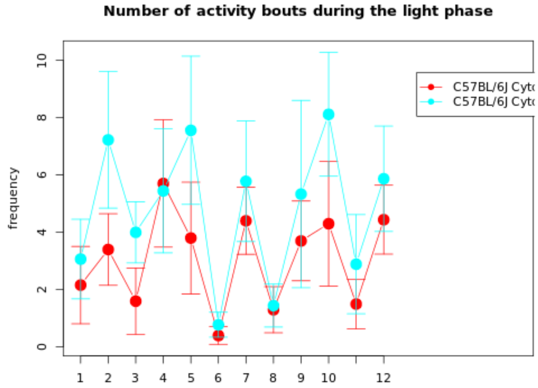

**NOTE: The statistics and boxplot are performed on log10 transformed data**

| Genotype | Mean | Median | SEM | N |
|---|---|---|---|---|
| (Gr 1): C57BL/6J Cytostatica bestraling Radiation (batch 22) | 32.9562 | 0 | Up: 5.6699<br>Down: 4.8586 | 10 |
| (Gr 2): C57BL/6J Cytostatica bestraling Sham (batch 22) | 50.905 | 0 | Up: 9.0349<br>Down: 7.6954 | 9 |

| | |
|---|---|
| **T-test:** | -1.9054 |
| **P-value:** | 0.0739 |
| **Df:** | 16.8401 |
| **Mann-Whitney U test:** | 0.094 |

| Mice excluded by Quality Control |
|---|
| PH06469 (C57BL/6J - ) |

 *© Sylics BV*

Feeding zone duration - light

*Feeding zone duration - light*: Cummulative duration in feeding zone during the light phase

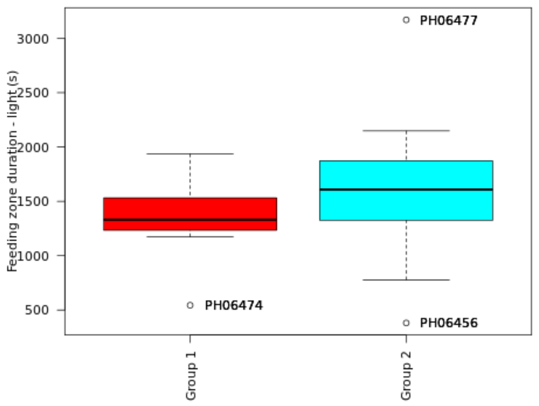

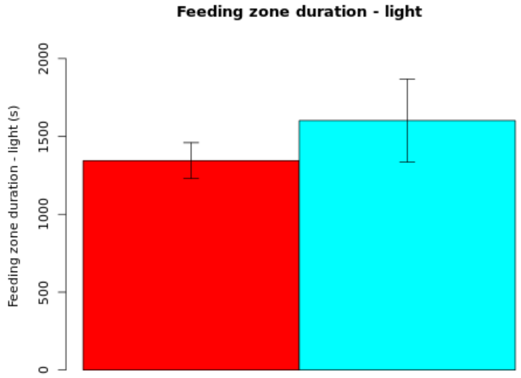

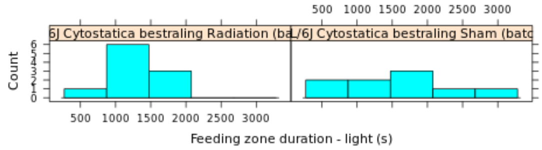

#### Secondary plot:

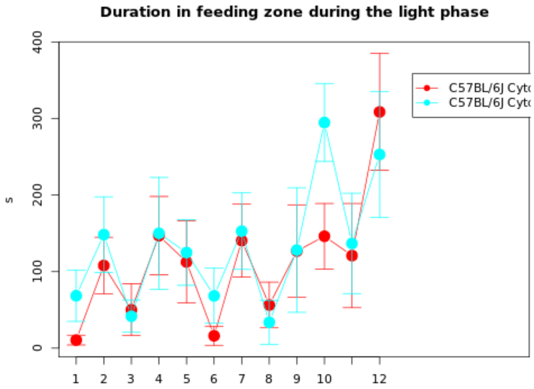

| Genotype | Mean | Median | SEM | N |
|---|---|---|---|---|
| (Gr 1): C57BL/6J Cytostatica bestraling Radiation (batch 22) | 1344.6013 | 1331.06 | 115.3727 | 10 |
| (Gr 2): C57BL/6J Cytostatica bestraling Sham (batch 22) | 1602.0281 | 1607.9067 | 266.3285 | 9 |

| | |
|---|---|
| **T-test:** | -0.8869 |
| **P-value:** | 0.3942 |
| **Df:** | 10.9418 |
| **Mann-Whitney U test:** | 0.3154 |

| Mice excluded by Quality Control |
|---|
| PH06469 (C57BL/6J - ) |

 © Sylics BV

OnShelter zone duration - light

*OnShelter zone duration - light*: Cummulative duration in OnShelter zone during the light phase

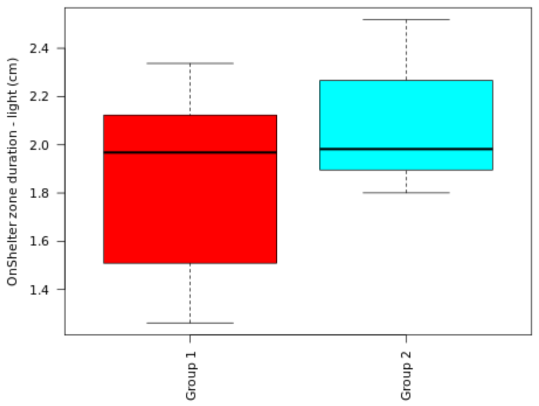

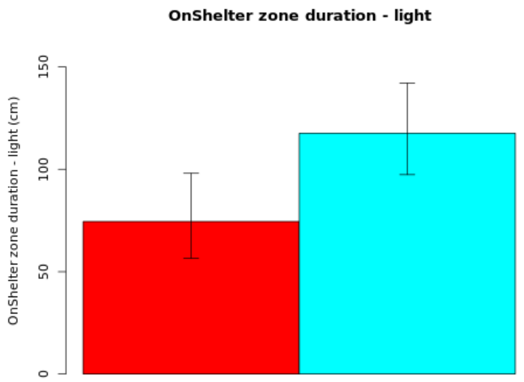

#### Secondary plot:

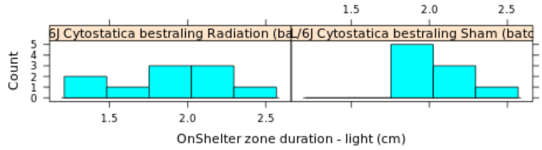

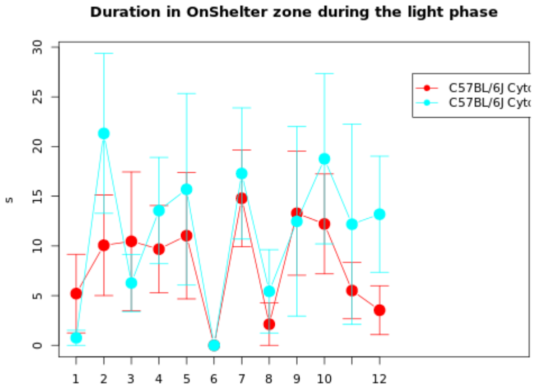

**NOTE: The statistics and boxplot are performed on log10 transformed data**

| Genotype | Mean | Median | SEM | N |
|---|---|---|---|---|
| (Gr 1): C57BL/6J Cytostatica bestraling Radiation (batch 22) | 74.4986 | 0 | Up: 23.5301<br>Down: 17.9391 | 10 |
| (Gr 2): C57BL/6J Cytostatica bestraling Sham (batch 22) | 117.6592 | 0 | Up: 24.4518<br>Down: 20.274 | 9 |

| | |
|---|---|
| **T-test:** | -1.3713 |
| **P-value:** | 0.1896 |
| **Df:** | 15.6321 |
| **Mann-Whitney U test:** | 0.4967 |

| Mice excluded by Quality Control |
|---|
| PH06469 (C57BL/6J - ) |

Sylics

OnShelter zone number - light

*OnShelter zone number - light*: Cummulative number of visits to OnShelter zone during the light phase

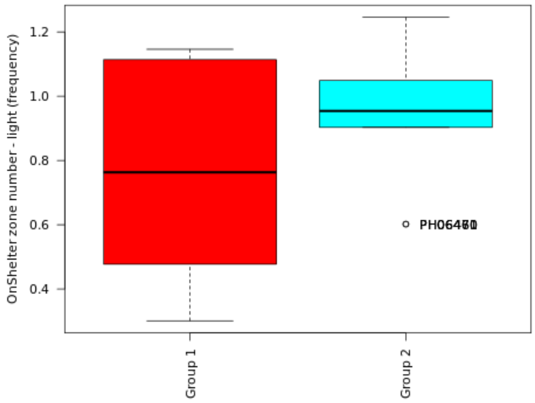

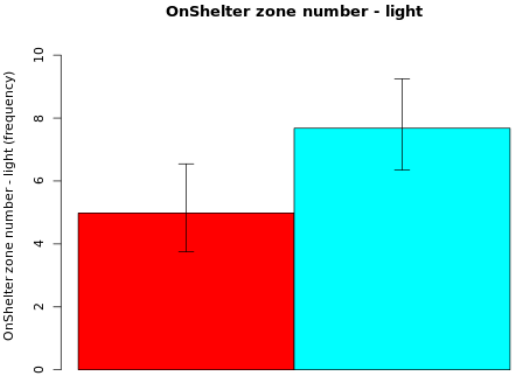

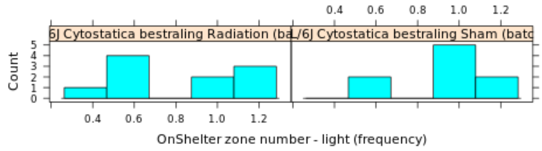

**NOTE: The statistics and boxplot are performed on log10 transformed data**

| Genotype | Mean | Median | SEM | N |
|---|---|---|---|---|
| (Gr 1): C57BL/6J Cytostatica bestraling Radiation (batch 22) | 4.9853 | 0 | Up: 1.5607<br>Down: 1.2379 | 10 |
| (Gr 2): C57BL/6J Cytostatica bestraling Sham (batch 22) | 7.6788 | 0 | Up: 1.5693<br>Down: 1.329 | 9 |

| | |
|---|---|
| **T-test:** | -1.3031 |
| **P-value:** | 0.2111 |
| **Df:** | 15.9065 |
| **Mann-Whitney U test:** | 0.3244 |

| Mice excluded by Quality Control |
|---|
| PH06469 (C57BL/6J - ) |

© Sylics BV

Sylics

Spout zone duration - light

*Spout zone duration - light*: Cummulative duration in Spout zone during the light phase

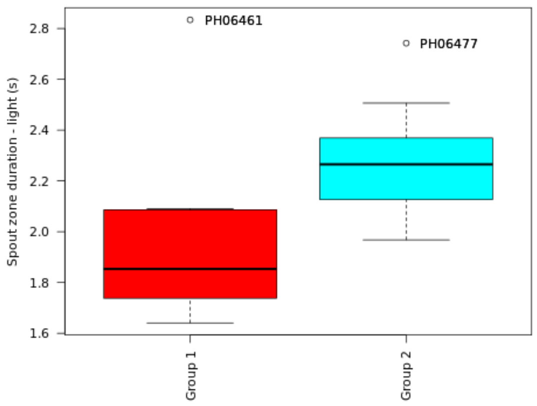

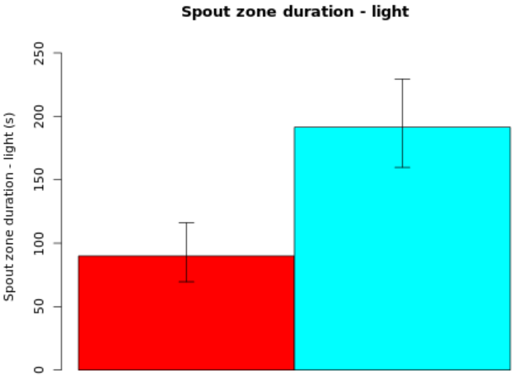

##### Secondary plot:

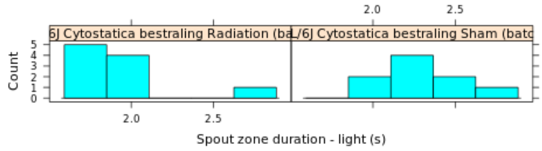

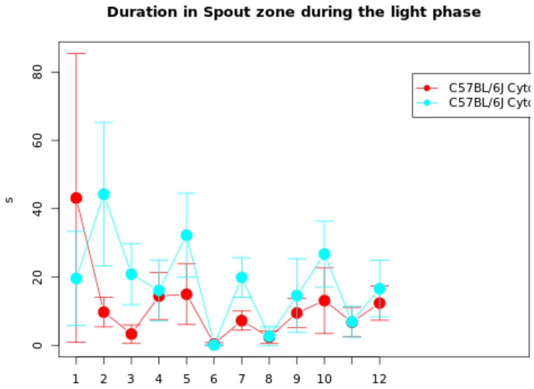

**NOTE: The statistics and boxplot are performed on log10 transformed data**

| Genotype | Mean | Median | SEM | N |
|---|---|---|---|---|
| (Gr 1): C57BL/6J Cytostatica bestraling Radiation (batch 22) | 89.9094 | 0 | Up: 26.1636<br>Down: 20.3165 | 10 |
| (Gr 2): C57BL/6J Cytostatica bestraling Sham (batch 22) | 191.4149 | 0 | Up: 37.9337<br>Down: 31.6868 | 9 |

| | |
|---|---|
| **T-test:** | -2.4155 |
| **P-value:** | 0.0282 |
| **Df:** | 15.8483 |
| **Mann-Whitney U test:** | 0.0101 |

| Mice excluded by Quality Control |
|---|
| PH06469 (C57BL/6J - ) |

© Sylics BV
